# Learning molecular traits of human pain disease via voltage-gated sodium channel structure renormalization

**DOI:** 10.1101/2025.02.19.639033

**Authors:** Markos N. Xenakis, Angelika Lampert

**Affiliations:** Institute of Neurophysiology, Uniklinik RWTH Aachen, Pauwelsstraße 30, Aachen, 52074, NRW, Germany; Scientific Center for Neuropathic Pain Research Aachen, SCN^AACHEN^, Uniklinik RWTH Aachen, Roermonderstraße 110a, Aachen, 52072, NRW, Germany

**Keywords:** Voltage-gated Sodium Channel, Renormalization, Criticality, Human Pain Disease, Machine Learning

## Abstract

Mammalian neurophysiology vitally depends on the stable functioning of transmembrane, pore-forming voltage-sensing proteins known as voltage-gated sodium channels (NaVChs). Deciphering the principles of NaVCh spatial organization can illuminate fundamental structure-function aspects of pore-forming proteins and offer new opportunities for pharmacological treatment of associated diseases such as chronic pain. Here, we introduce a renormalization group flow paradigm permitting a formal investigation of NaVCh thermostability properties. Our procedures are solidified by deriving a *q* -deformed statistical mechanical entropy and validated over 121 experimentally resolved NaVCh structures of prokaryotic and eukaryotic origin. We uncover the universality of a critical inflection point regulating the thermostability of the pore domain relative to the voltage sensors, summarized in terms of a generalized Widom scaling law. A machine learning algorithm, rationalized in terms of the ‘loosening’ of inertia and conductivity channel constraints, identifies pain-disease-associated mutation hotspots in the human NaV1.7 channel. Our work illustrates how first-principles-based machine learning approaches can deliver accurate insights for human pain medicine and clinicians at a reduced computational cost, while clarifying the self-organized critical nature of NaVChs.

## 1 Introduction

The primary neurophysiological role of voltage-gated sodium channels (NaVChs) is to initiate and propagate action potentials along neuronal tissue [1]. Accordingly, NaVChs act as ‘traffic controllers’ for sodium ions crossing the cell membrane, thereby shaping the upstroke phase of action potentials [2, 3].

The key property that makes sodium ion transport entropically favorable across the hydrophobic membrane environment is hydrophilicity [4]. Hydrophilic groups coordinate the dehydration of sodium ions along a NaVCh pore, facilitating the ion’s passage through the membrane [5–7]. The physical principle underlying NaVCh selectivity is thought to arise from a delicate balancing of strong and potentially long-range interactions between the selectivity filter (SF) and surrounding residue clusters [8], as inferred from early electrophysiological studies [9].

Members of the NaVCh superfamily follow a common structural organization principle: they form a porous-like environment composed of four radially arranged homologous domains (labeled DI-IV) [10]. The first NaVCh structure to be crystallized was the prokaryotic sodium channel from Arcobacter butzleri (NaVAb), captured in a pre-open state [11]. As anticipated from earlier crystallographic studies on potassium channels [12, 13], the NaVAb crystal verified that each domain comprises a pore module (PM) interlinked with a voltage sensor domain (VSD) [11]. The PM is composed of two antiparallel alpha-helical segments (labeled S5-6) connected by an extracellular loop and the SF [11]. The spatial arrangement of the four PMs inside a NaVCh gives rise to the pore domain (PD) sub-architecture, where a narrow, sieve-like extra-cellular SF-lined environment, followed by a hydrophobic central cavity, leads to an intracellular pore narrowing where the putative activation gate (AG) is located [11]. On the other hand, the VSD consists of four transmembrane alpha-helices (labeled S1-4) connected to the PM by a linker bridging S4 with S5 [11]. Positively charged residues in the S4 helix can sense the membrane depolarization stimuli [14]. Their outward movement exerts a pulling force that opens the pore [14–17]. In contrast to their prokaryotic ancestors, eukaryotic NaVChs display a more diverse structural and functional profile, mirroring the breaking of radial structural symmetries [10]. Consequently, the dynamic repertoire of eukaryotic NaVChs is richer, yet more specialized and allosterically efficient compared to their prokaryotic counterparts [18–20].

Complex biomolecules, such as NaVChs, have evolved to balance sensitivity and robustness to perturbations of environmental and genetic origin [21, 22]. Concretely, a search in the gnomAD [23] database reveals thousands of benign variants associated with genes encoding NaVChs in humans (e.g., 729 benign variants are found in the *SCN9A* gene [24]), encoding the human NaV1.7 channel molecule which is crucial for human pain perception [25]. This implies that NaVChs, albeit complex, exhibit substantial structural and functional tolerance to random variations in the encoding genetic material. What physical principle guarantees this?

It turns out that protein systems leverage self-similarity (or scale-invariance [26]) to optimally distribute internal energies and external stresses throughout their structure [27, 28]. This phenomenon is known as self-organized criticality (SOC) [29]. SOC is believed to be evolution’s answer to the need for maintaining an internal molecular state of high functional sensitivity without sacrificing mutational robustness [21, 22, 27, 28, 30, 31]. In logical connection with early studies demonstrating the universal characteristics of hydropathy plots in globular proteins [32–35], SOC signatures manifest in protein molecules as a pattern of extrema (peaks and valleys) of intrinsic thermostability cost functions associated with water-mediated interactions [30, 31].

Scaling analysis of the hydropathic dipole field acting along the pore shed light on SOC phenomenology in the NaVAb molecule captured at a pre-open state [36] and in a computationally optimized NaV1.7 molecule found in humans [37]. It was demonstrated that the distribution of atoms around the pore of the NaVAb molecule is predominantly unimodal, meaning the corresponding cumulative distribution function is characterized by a prominent inflection point [36, 37]. Also, the amplitude of the hydropathic dipole field exhibited a clear self-similar increment about the pore axis, leading to a distinct peak near the inflection point [36, 37]. Moreover, the radial location of this prominent inflection point marked the characteristic size of the PD, i.e., the distance from the current pore point where the structural transition from the PD to the VSDs takes place [36, 37]. Based on this triple coincidence, it is compelling to conceptually parallel the spatial organization of atoms around the pore with the archetypal SOC sandpile model [29] as a means to develop a toy model for NaVCh molecular complexity [38].

Accordingly, residue structural positions are abstracted as lattice sites, with mutations analogous to externally added sand grains perturbing the lattice structure. Slowly adding sand grains to the lattice leads to the formation of a theoretical sandpile. Here, ‘theoretical’ indicates that the sandpile itself is not a direct observable. Rather, its defining characteristic, defined by its slope, can be inferred from the scaling (power-law) exponents [30, 31, 36, 37], characterizing the self-similar incrementing of the underlying hydropathic dipole field about a pore point. Each pore point thus serves as an initial condition for traversing the sandpile at a varying slope determined by the self-similar property of the locally acting field [36, 37]. If a sand grain lands on a site where the slope of the sandpile is too steep, it is expected to adversely affect the stability of the molecule. Stated differently, a steep sandpile slope prescribes a high likelihood of mutation-induced destabilization. This insight provided a rationale for distinguishing between pain-disease-associated and benign structural locations in NaV1.7 [37].

A self-consistent scaling theory for proteins should be grounded in the renormalization group (RG) framework [39]. Here, we examine this concept within the NaVCh superfamily and demonstrate how it can yield biomedically relevant insights concerning inherited forms of human pain disease. Specifically, starting from the simplest possible radial differential equation that can rationally support a pore-forming molecular architecture with a PD/VSD interface, we derive the RG flow equation facilitating a substantial yet biophysically meaningful reduction of molecular degrees of freedom (DoF) about a NaVCh pore. This yields universal insights into dimensionality reduction, entropy constraints, and symmetry breaking of hydropathic dipole fields across the NaVCh superfamily, as revealed by atomic-level analysis of 121 sufficient-resolution NaVCh structures of prokaryotic and eukaryotic origin. At a single-NaV1.7-molecule level, we rigorously describe mutation clustering relative to the PD/VSD interface, establishing a transparent machine learning framework for detecting NaV1.7 structural locations where the steepening of the sandpile slope correlates with a human pain disease phenotype.

## 2 Methods

### 2.1 Analytical procedures

We assume that the NaVCh molecule has settled into a reasonably long-lived or frequently visited metastable state.

#### 2.1.1. Spatial organization principles

We focus on a pore point, **p** = (p_*x*_, p_*y*_, p_*z*_ ≡ P_⊥_) ∈ 𝒫 (SI Eq. (S1) and Fig. 1(a)), and introduce the open ball:

**Fig. 1.**
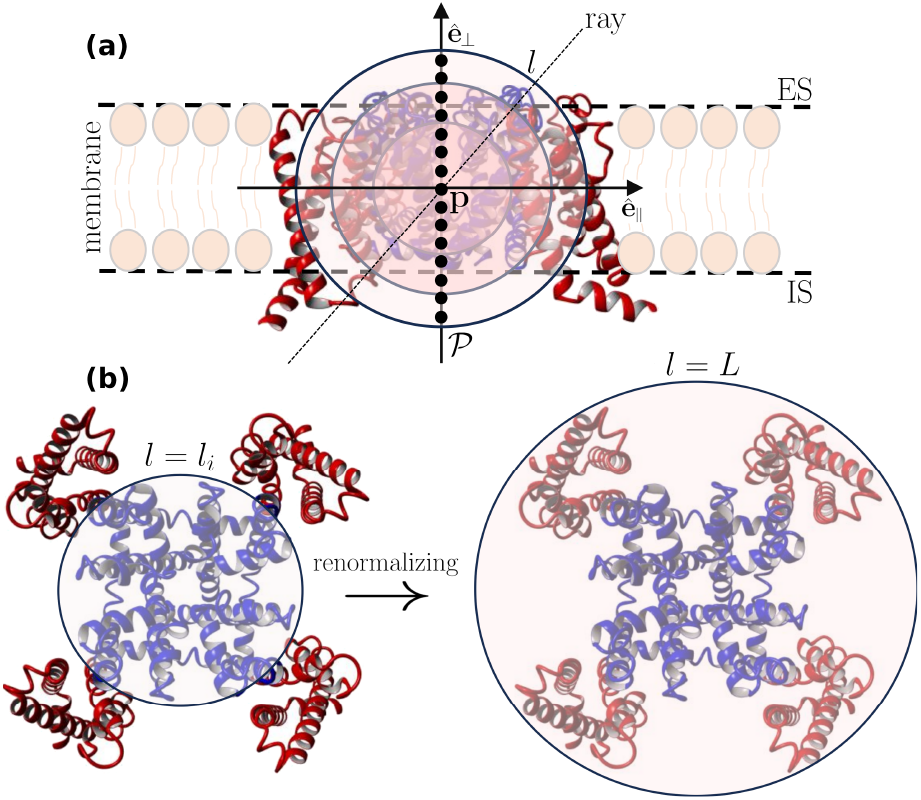
Geometry and renormalizability of the prototype NaVAb channel. **(a)**, The illustrated molecular side view corresponds to a pre-open NaVAb molecule (PDB code: 3rvy). The pore domain (PD) and voltage sensor domains (VSDs) are illustrated in blue and red, respectively. **ê**_∥_ and **ê**⊥ are the membrane-parallel and membrane-perpendicular unit vectors, respectively. We introduce consecutive pore points, **p** ∈ 𝒫 (SI Eq. (S1)), forming a path through the NaVAb pore. Each pore point serves as the center of an ensemble of nested balls, each of them characterized by a radius *l* [Å] (Eq. (1)). An infinite number of paths (rays) can pass through a pore point, radiating in all possible directions. Analyzing the properties of the atomic environment along individual radial paths is impractical due to their infinite number. The renormalization group procedure solves this problem by collapsing radial paths into *l*, rendering atomic environment properties inherently dependent on *l*. This is equivalent to *coarse-graining* [85], and enables the computation of relevant scaling exponent. For example, substituting ln *n* for 𝒪 in Eq. (30), returns via (31) the order parameter scaling exponent, 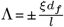 (SI Eq. (S15)). **(b)**, Top view, from the extracellular side (ES). *l*_*i*_ [Å] represents the characteristic size of the PD, also marking the inflection point of the cumulative atom number (Eqs. (5),(6)). *L* [Å] is the radius of the smallest ball that contains the entire molecule (SI Eq. (S4)). IS stands for intracellular side.

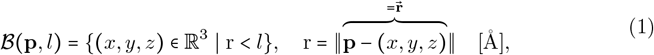

where *l* [Å] is the probe radius and 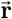 is the vector from **p** to a candidate atom coordinate, (*x, y, z*). ∥ ⋅ ∥ is the Euclidean norm. *∂*ℬ = {(*x, y, z*) S r = *l*} denotes the spherical surface of ℬ. In a structural biology context, *∂*ℬ is called an *interface*.

Mathematical functions and associated parameters appearing henceforth depend on (**p**, *l*) and **p**, respectively. For clarity, this dependence is omitted.

##### 2.1.1.1 Atom packing in the continuum

The number of atoms residing on *∂*ℬ is continuously approximated with the following generalized logistic differential equation [36, 40]:

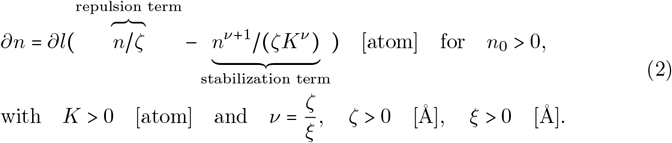

*n* is the number of atoms residing in ℬ. *ζ* represents the effective range over which atoms push each other apart (i.e., repel each other). On the other hand, *ξ* represents the effective range over which atoms pull each other together (i.e., attract each other). *n*_0_ = *n* (*l*_0_) is the atom packing initial condition realized for some initial *l*-value, *l*_0_. Meaning *n*_0_ is the number of atoms residing in the initial ball, 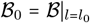. *K* [atom] is the carrying capacity of the NaVCh structure. It delimits the number of atoms residing in ℬ for *l* →∞ (see also (3)), thereby setting an upper bound on the number of atoms the structure can accommodate.

We propose an effective phenomenology supporting (2) based on the interplay between hydrophobic attraction and hydrophilic repulsion [41]. The nonlinear term in the right hand side (RHS) of (2) accounts for attractive, hydrophobicity-driven atomic interactions stabilizing an *n*-cluster. Namely, it reflects the tendency of hydrophobic constituents to ‘hide’ inside ℬ. The term *n*^*ν*+1^ indicates that any group of *ν* + 1 atoms can form an interaction subnetwork. Since no spatial constraints apply to the formation of such a group, *n*^*ν*+ 1^ accounts also for long-range interatomic interactions. Hence, *ν*= *ζ* /*ξ* explains the emergence of long-range interactions through bond length adjustment driven by atomic repulsivity.

Accordingly, the linear term appearing in the RHS of Eq. (2) accounts for repulsive, hydrophilicity-driven interactions pushing atoms toward *∂*ℬ. This is attributed to space-filling water structuring effects that increase the energy required for efficient atom packing, according to the thermodynamic self-assembly principles outlined in [42]. Additionally, *ζ* incorporates the Pauli exclusion principle, as discussed in Ref. [43].

##### 2.1.1.1 The characteristic size of a pore domain

Solving (2) for *n* gives:

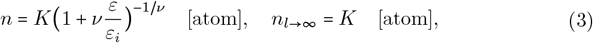

where

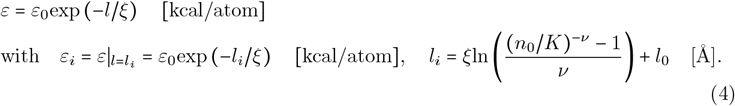

*ε* imposes an upper bound on the average unsigned hydropathic energy of an individual atom. On the basis of simplicity, *ε*_0_ [kcal/atom] is chosen to be a constant. Accordingly, *n*_0_*ε*_0_ [kcal] delimits the absolute hydropathic energy stored in the initial ball, ℬ_0_. 1/*ξ* [Å^− 1^] can then be interpreted as the rate at which hydrophobic energy surpluses, stored in ℬ _0_, are consumed in useful (stabilizing) interactions as the *n*-cluster size increases. We note that this interpretation is simply a reformulation of our initial understanding about 1 /*ξ* deduced from (2). Specifically, decreasing *ξ* increases *ν*, thereby increasing the likelihood of distant atoms interactions, reflecting the faster depletion of hydrophobic energy surpluses into stabilizing, potentially long-range bonds.

We do not seek a biophysical interpretation for the logarithmic argument, 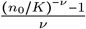. Instead, we focus on *l*.

We emphasize that *l*_*i*_ marks the *l*-value for which the second-order derivative of *n* with respect to *l*, i.e., the curvature of *n* along the radial direction,

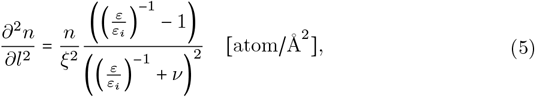

becomes zero. Stated differently, *l*_*i*_ is an inflection point of *n*. It follows that for *l* = *l*_*i*_, (2) is globally maximized, i.e.,:

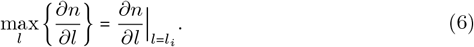

The biophysical significance of (6) becomes apparent when considering empirical evidence suggesting that 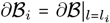 serves as an approximation of the interface mediating the structural transition from the PD to the VSDs [36, 37]. Accordingly, we treat *l*_*i*_ as the characteristic length scale of the PD sub-architecture, i.e., the maximum size at which the PD retains its essential functional and structural characteristics, independent of VSD influence (for an illustration see Fig. 1(b), left). Namely, beyond *l*_*i*_ (i.e., for *l* >*l*_*i*_), the coupling interactions between a pore module and a radially succeeding voltage sensor cannot be neglected, driving the structural transition from the former towards the latter.

The irregular cylindrical surface emerging from the dense arrangement of ℬ_*i*_ balls along 𝒫 is parameterized by:

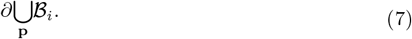

(7) serves as the characteristic cover for the PD, in the sense that it incorporates essential structural elements of the PD sub-architecture.

##### 2.1.1.3. Mean-field conditioning of the atomic environment

The length scale

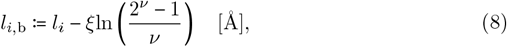

denotes the *l*-value for which the atom packing condition *n*/*K* = 0.5 is satisfied, meaning exactly half out of the *K* atoms are residing in 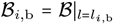. ‘b’ stands here for ‘bound’, since *l*_*i*,b_ imposes an upper and lower bound on *l*_*i*_, depending on whether the attractive or repulsive interaction range prevails, i.e., whether *ν*< 1 or *ν*>1, respectively. Accordingly, the negative and positive sign of ln 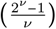 decides whether the length scale bound applies from below or from above, i.e., whether *l*_*i*,b_ > *l*_*i*_ or *l*_*i*,b_ < *l*_*i*_, respectively.

If the attractive interaction range equals the repelling interaction range (i.e., *ν*= 1), then ln 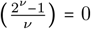 and *l*_*i*,b_= *l*_*i*_, and (3) becomes isomorphic with the Fermi-Dirac(i.e., logistic) distribution. Hydrophobicity-driven attraction and hydrophilicity-driven repulsion are then delicately balanced, imposing a mean-field conditioning on the atomic environment around a pore point. Across an evolutionary time scale, *ν =* 1 promotes isotropic space exploration in both the PD and the VSDs, as an equal number of atoms is distributed in the PD and the VSDs (i.e., half of the *K* atoms are found in ℬ_*i*_ = ℬ_*i*,b_ and the other half outside of it). Generally, this favors the emergence of globular-like molecular shapes, as demonstrated in Ref. [43], which, in the context of this study, is driven by a roughly equal and uniform allocation of masses inside and outside ℬ_*i*_.

##### 2.1.1.4 Unit mass fractal dimension

Expressing (2) in terms of (3), reveals that packing of atoms around a pore point satisfies the self-similarity relationship:

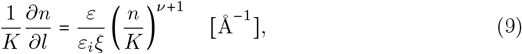

which implies that:

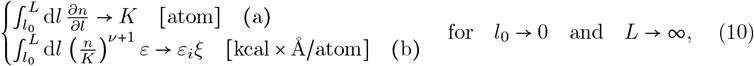

where ℬ_*L*_ *=* ℬ|_*l* = *L*_ is the smallest ball that accommodates all NaVCh structure atoms (SI Eq. (S4) and Fig. 1(b) right part).

(10) introduces two distinct yet interrelated constraints. Specifically, (10)(a) and (10)(b) ensure that even if the molecular radial size, *s* [Å] (SI Eq. (S5)), becomes infinitely large, the total number of atoms cannot exceed *K*, and the hydropathic interaction energy per atom converges to *ε*_*i*_*ξ* [kcal×Å/atom], respectively.

The intrinsic dimension (i.e., the unit mass fractal dimension [44]) of an *n*-cluster is continuously measured with:

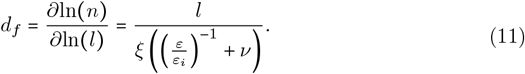

(11) describes how ‘intensively’ (or ‘compactly’, as discussed in Ref. [45]) *n* atoms fill space in ℬ. Generally, the fraction of empty space in ℬ increases with decreasing *d*_*f*_.

#### 2.1.2 Statistical mechanical entropy

Let us now consider the following discretization of *l*:

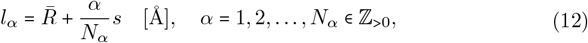

where *α* is a scale index [36, 40]. *N*_*α*_ is the total number of scale iterations, determining the resolution of the discretization. 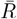 [Å] is the distance separating **p** from its nearest-neighbor atom, while *s* [Å] remains the molecular radial size (first considered below (10) and given by SI Eq. (S5)). Note that 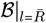 is empty of atomsThe probability that an atom resides in 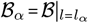 is given by:

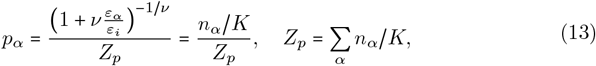

where (1 + *νx*)^−1/*ν*^, *x* ≥ 0, is identified with the survival function of a Pareto Type-II distribution [46]. *Z*_*p*_ is the partition sum guaranteeing that ∑_*α*_ *p*_*α*_ = 1

The probability that (*ν*+ 1)-atoms reside in ℬ_*α*_ is given by the Escort probability [47]:

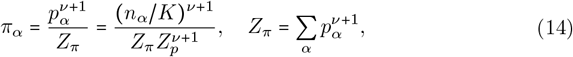

where *Z*_*π*_ is the corresponding partition sum.

*π*’s biophysical importance stands out when we consider the following finite-size corrected version of (10(b)):

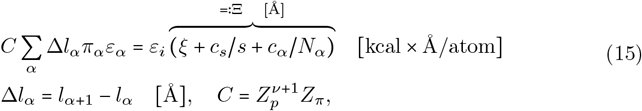

where Ξ is a finite-size corrected version of *ξ. c*_*s*_ [Å^2^] and *c*_*α*_ [Å] account for interfacial and radial geometric distortions due to *s*’s and *N*_*α*_’s finiteness, respectively. Note that Δ*l*_*α*_ is constant. *C* is a normalization factor whose specific value is irrelevant. Crucially, for *N*_*α*_ →∞, the constraint (10)(b) can be recovered:

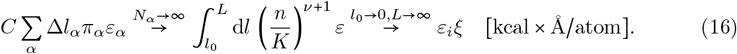

An entropy functional whose maximization explicitly incorporates (13)-(15) is given by:

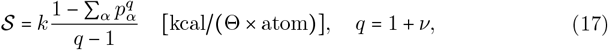

where *q* is known as the nonextensivity index [48, 49] and *k* [kcal/(Θ × atom)] is the equivalent of Boltzman’s constant. Note that [Θ] is the physical unit at which the temperature of the current pore point is measured (SI 1.4).

From an information theoretical viewpoint, 𝒮 can be trivially interpreted as the average amount of surprise (or ‘unexpectedness’) associated with determining how many atoms reside in ℬ._*α*_

From an evolutionary perspective, 𝒮 illustrates how the competition between hydrophobicity-driven attraction and hydrophilicity-driven repulsion determines the information content of the atomic environment around a pore point.

Local scarcity of hydrophilicity can reduce associated repulsion effects (i.e., *ζ* → 0), promoting the formation of a hydrophobic core. 𝒮 approaches then the following upper bound:

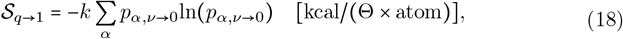

where 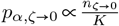 is a common exponential (SI Eqs. (S6),(S10)).

Maximizing 𝒮 toward 𝒮 _*q* →1_ expands the NaVCh configuration space volume, since the amount of water required to solvate a hydrophobic *n*-cluster packed in ℬis typically less than that required for solvating *n* individually dispersed hydrophobic atoms. The remaining, non-solvating water molecules can then uncoordinately engage in hydrogen bond interactions with NaVCh constituents on *∂*ℬ, effectively increasing the NaVCh configuration DoF as water structuring effects diminish. Simultaneously, we notice that the intrinsic dimension of the PD,

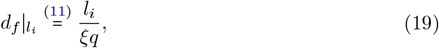

is decreased, since

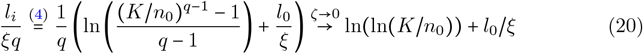

is monotonically and positively correlated with *ζ* for fixed parameter values {*l*_0_, *n*_0_, *K, ξ*} (SI Fig. S1 inset).

(20) illustrates how the PD sub-architecture can mitigate the potentially disordering effect of excess entropy by lowering its intrinsic dimension.

#### 2.1.3 Thermostability

##### 2.1.3.1 Empirical measure

The thermostability of an *n*-cluster is probed with the hydropathic moments toolbox [32–37, 40]:

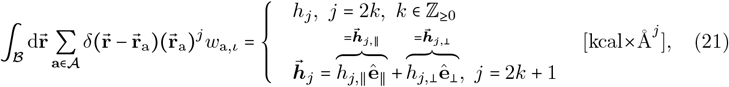

where *δ* (⋅) is a generally defined Dirac delta function, **a** ∈ **𝒜** is an atom coordinate with 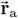 denoting the vector from **p** to **a**, and *w*_a,*ι*_ = *w*_a_ + *ι* [kcal], *ι* ∼𝒩 (0, *σ*_*ι*_ = 1e − 03), are noise-perturbed hydropathic weights originating from the Kapcha-&-Rossky atomic hydropathic scale [50]. Noise accounts here for randomly occurring water density fluctuations.

Note that 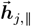 and 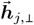 act parallel and perpendicularly, respectively, to the membrane, surface with **ê**_∥_ and **ê**_⊥_ being the corresponding unit vectors (SI S1.2.3). Conventionally, **ê**_⊥_ points towards the extracellular side (ES).

The subscript ‘*j*’ indicates the moment order determining the dimension of the space within which water-mediated interactions are probed. *j*-parity determines whether we probe inertia (*j* = 2*k*) or conductivity (*j* = 2*k* + 1) constraints, concerning rotational and translational DoF expressed around and along the pore point path, 𝒫, as detailed in SI 1.5.1-2, respectively.

##### 2.1.3.2. Theoretical model

Normalizing *h*_*j*_ over *n* yields the *n*-cluster hydropathic density [32–35] whose oscillatory behavior is described by:

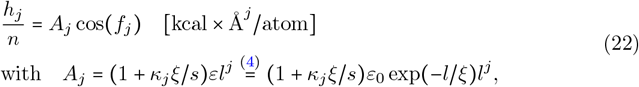

where *f*_*j*_ determines the sign changing behavior of *h*_*j*_ and *A*_*j*_ envelopes the underlying wave packet. The factor (1 + *κ*_*j*_*ξ* /*s*) > 0 introduces a finite-size correction with *κ*_*j*_*ξ* [Å] being the adsorption bond length. *κ*_*j*_ is dimensionless and purely phenomenological, as it can only be deduced by fitting *A*_*j*_ cos(*f*_*j*_) to experimental *h*_*j*_ /*n*-traces. It generally describes the degree of adhesion of atoms on *∂*ℬ in response to interfacial tensions.

Notably, *A*_*j*_ preserves the general form that the hydropathic energy has been assigned in Refs. [51–53]. Moreover, consistent with Eq. (4), for *j* = 0, we verify that 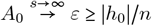, illustrating that *ε* serves as an upper bound for the average unsigned hydropathic energy of an individual atom.

##### 2.1.3.3 Scaling of hydropathic dipole field amplitude

Hydropathic energy wave packet self-modulation effects persisting over sufficiently large *l*-intervals covering in the PD and the VSDs are described by [36, 37]:

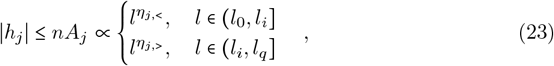

where *l*_*q*_ [Å] denotes the *l*-value for which the *n*-curvature (Eq. (5)) is negatively maximized.

(*η*_*j*, <_, *η*_*j*, >_) ∈ ℝ^2^ are the corresponding scaling exponents. |*η*_*j*, <_| and |*η*_*j*, >_| set an upper bound on how ‘intensively’ *j*-th-order dipole-dipole hydropathic interactions can fill space in the PD and the VSDs, respectively.

The sign of *η*_*j*_ indicates the direction of hydropathic interaction network intensification, either inwards (i.e., *η*_*j*_ < 0) or outwards (i.e., *η*_*j*_ >0).

#### 2.1.4 Mutational robustness

##### 2.1.4.1 Decomposability

Sign-changes of *h*_*j*_ mark *n*-clusters of diminished thermostability. Mechanofunctional properties of these *n*-clusters are expected to be sensitive to perturbations.

To understand how the multiscale competition between hydrophobic attraction and hydrophilic repulsion can lead to varying thermostability and emergent mechanofunctional properties, we consider the following decomposition ansatz:

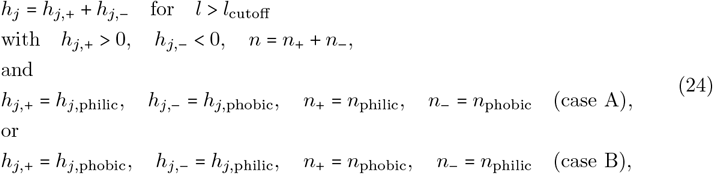

where *l*_cutoff_ is the ansatz cutoff scale.

We verify that for vanishingly small noise, i.e., *σ*_*ι*_ → 0 (see (22)), and *j* = 2*k*, case A holds. Namely, *h*_*j*,+_ = *h*_*j*,philic_ and *h*_*j*,−_ = *h*_*j*,phobic_ are the contributions of only hydrophilic and hydrophobic atoms, respectively, with *n*_+_ = *n*_philic_, *n*_−_= *n*_phobic_ and *l*_cutoff_→ *l*_0_.

Whether case A or B holds for the pair {*h*_2*k*+1,⊥,philic_, *h*_2*k*+ 1,⊥,phobic_} requires computational investigation, as there is no general argument supporting this operation. Since *h*_*j*,_ + and *h*_*j*,_ − are cumulative, and thus slowly changing, sign-preserving functions over the *l*-range, we can straightforwardly compute their continuously varying scaling exponents:

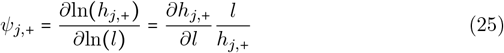

and

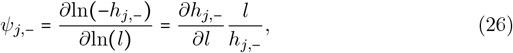

respectively.

The physical meaning of *ψ*_*j*,+_ and *ψ*_*j*,−_ complements that of *η*. Namely, |*ψ*_*j*,+_| and |*ψ*_*j*, −_| account for the intensification of the *j*-th-order dipole-dipole interaction network stabilizing the subgroups of *n*_+_ and *n*_−_ atoms, respectively, along the radial direction. The signs of *ψ*_*j*,+_ and *ψ*_*j*,−_ indicate the direction of interaction network intensification, being either inwards (if *ψ*_j_ < 0) or outwards (if *ψ* > 0)._*jj*_

##### 2.1.4.2 Logarithmic composite susceptibility

The difference of *ψ*_*j*,−_ from *ψ*_*j*,+_,

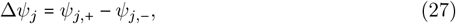

illustrates how the interaction networks, stabilizing the subgroups of *n*_+_ and *n*_−_ atoms, ‘compete’ to fill space in ℬ.

We postulate that when Δ*ψ*_*j*_ = 0, the two networks are in a state of balance, where perturbations affecting the *n*_+_ and *n*_−_ subgroups tend to induce oppositely directed *∂*ℬ-responses, effectively canceling each other out. On the other hand, Δ*ψ* ≠ 0_*j*_ implies an interaction network imbalance, determining also the prevailing direction of a *∂*ℬ -response. Accordingly, the sign of Δ*ψ*_*j*_ informs about the direction of perturbation-induced *∂*ℬ-responses.

Dividing Δ*ψ*_*j*_ by *l* defines the following size-normalized measure of interaction network imbalance:

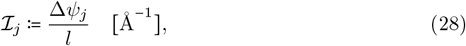

assuming that the interaction network imbalance is uniformly distributed over the radial extent of an *n*-cluster.

Integrating ℐ_*j*_ over an *l*-range, yields the logarithm of the *j*-th-order composite susceptibility, i.e.,:

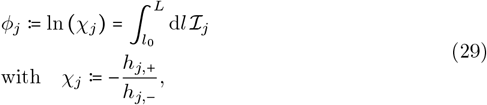

with *χ*_*j*_ being identified with the *j*-th-order composite susceptibility. Generally, *χ*_*j*_ can be interpreted as a descriptor of how internal hydrophobicity-generated fields couple with internal hydrophilicity-generated fields and vice versa.

Treating 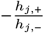 as an effective temperature ratio reveals that |*φ*_*j*_| is the equivalent of a thermostability geodesic (i.e., shortest distance measure) (SI S1.6.1). Simply put, |*φ*_*j*_| quantifies how ‘far’ an *n*-cluster’s thermostability state is from a state of minimal thermostability (or, maximal sensitivity to perturbations).

We emphasize that although both *φ*_*j*_ and *h*_*j*_ qualify as thermostability ‘distance’ measures (since *h* _*j*_= 0 =⇒ (24) *φ* _*j*_= 0), the geodesic property applies exclusively to |*φ*_*j*_|. In fact, while *η*_*j*_ (Eq. (23)) provides information at an envelope level, Δ*ψ*_*j*_ (27) reflects the actual sculpting (i.e., the radial interplay of successive peaks and valleys) of the hydropathic energy wave packet.

On the ground of consistency, ℐ _*j*_ is considered to provide equivalent information with *φ*_*j*_ but at an interface level. Namely, | ℐ _*j*_ | measures how ‘far’ an *∂n*-shell’s thermostability state is from a state of minimal thermostability. We term ℐ_*j*_ the ‘interfacial coupling strength’.

#### 2.1.5 Renormalizability

The RG flow equation has the general form [54]:

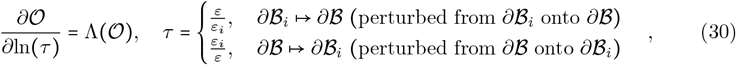

where 𝒪 := 𝒪 (**p**, *l*) is a NaVCh structure log-observable (referred to as the *effective coupling* [54]). *τ* is the normalized hydropathic energy *scale* [54], representing the relative temperature change in response to PD/VSD interface fluctuations (SI Eq. (S13)). Λ describes the dependence of 𝒪 on *τ* and is the equivalent of the *beta function* [54]. It represents a continuously varying scaling exponent attaining its critical value precisely at *τ* = 1 (SI 1.7).

The conceptual basis upon which (30) is built, is illustrated in Fig. 1(b) along with its caption.

From (30), we straightforwardly obtain the renormalizability relationship:

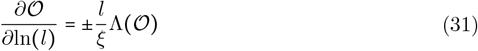

Eq. (31) establishes that changes 𝒪 in over a logarithmic radial range (on the left hand side of (31)) can be equivalently obtained by considering changes in 𝒪 driven by PD-subarchitecture deformations reflected upon PD/VSD interface fluctuations (on the right hand side of (31)).

### 2.2 Computational Procedures

#### 2.2.1 Structure preparation

NaVCh structures are ‘cleaned’, protonated, and their orientation is fixed, following the procedures outlined in SI S1.2.

#### 2.2.2 Curve fitting and model selection

Candidate *n*-models were fitted to the dimensionless empirical cumulative atom number:

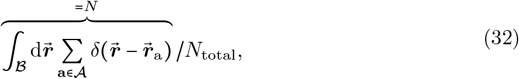

where *N*_total_ = card (𝒜) is the total number of atoms found in a ‘clean’, protonated NaVCh structure. Algorithmic implementation details are enclosed in SI 1.11.

## 3 Results

### 3.1 NaVCh spatial organization

We compiled a structure dataset comprising 71 prokaryotic NaVCh structures (sub-types: NaChBac, NaVAb, NaVMs, NaVAe, NaVRh) and 50 eukaryotic structures (subtypes: NaVPas, NaVEe1, NaVEh, NaV1-8) of sufficient resolution (SI S1.1).

#### 3.1.1 Geometry

##### 3.1.1.1 Inflection point universality

We report an excellent agreement between the empirical cumulative atom number, *N* (Eq. (32)), and its theoretical counterpart, *n* (Eq. (3)), across all 121 structures (Figs. 2(a),(g), SI Figs. S5-S10(a), S12-S22(a)), supported by small mean absolute fitting errors (SI S2.1). P-values remained inconclusive, as expected, due to the coarsegrained nature of our model and their sensitivity to noise (SI S2.1).

**Fig. 2.**
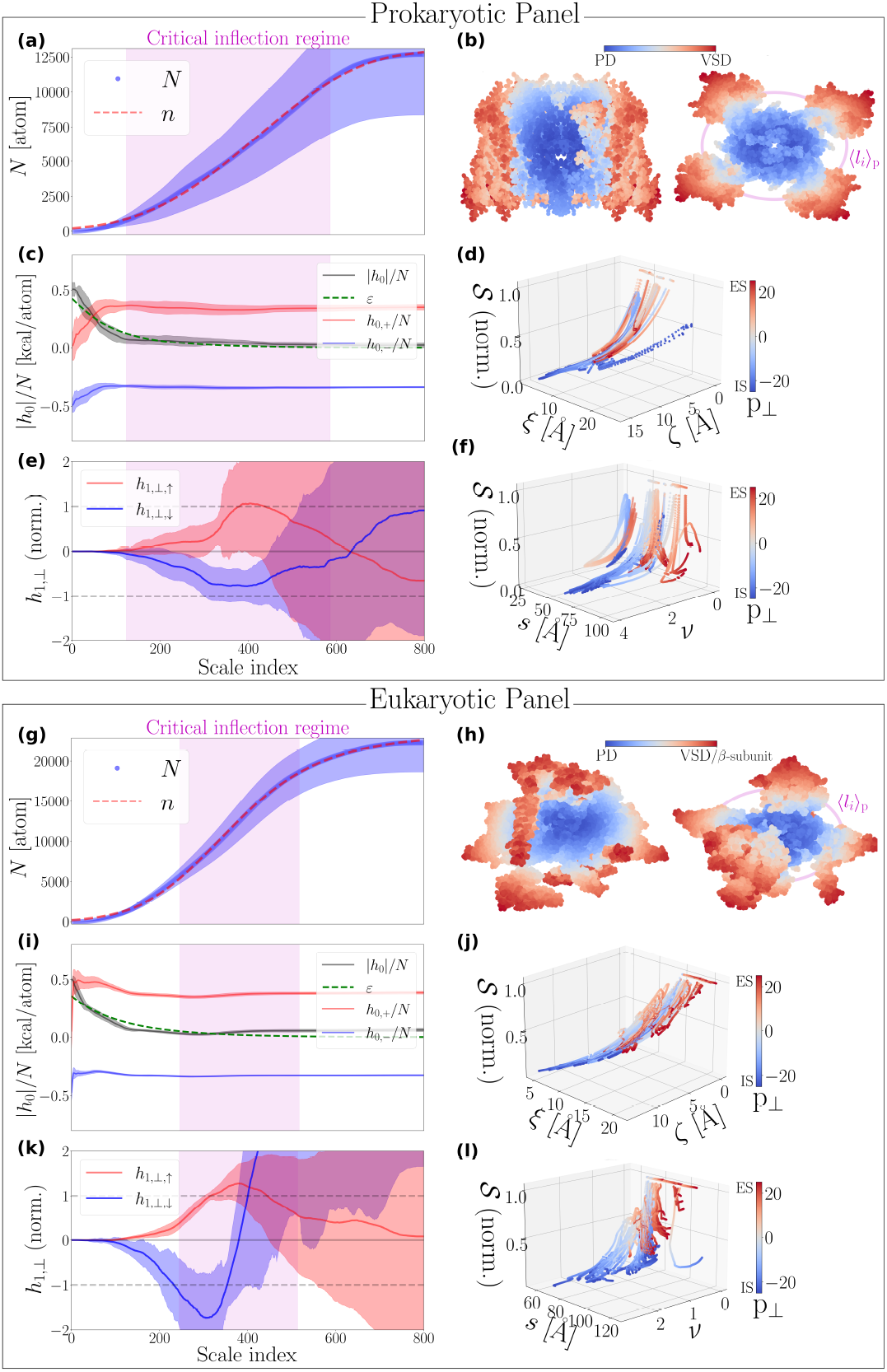
Spatial organization features of the NaVCh superfamily. Statistical summary of the spatial organization features of 71 prokaryotic (subtypes: NaChBac, NaVAb, NaVMs, NaVAe, NaVRh) and 50 eukaryotic (subtypes: NaVPas, NaVEe1, NaV1-8, NaVEh) NaVCh atomic structures (SI Tab. S1), for pore points, **p** ∈ 𝒫, and scale indices, *α* = 1, 2, …, 800. **(a)**,**(g)**, Collapsed traces of the empirical, *N* [atom], and best-fitted theoretical, *n* [atom], cumulative atom numbers. **(b)** and **(h)** illustrate the prototype NaVAb channel (PDB code: 3rvy) and the human NaV1.7 channel (PDB code: 7w9k), respectively. Atoms are colored according to their PD/VSD ordering score (SI S1.8.1.2), and projected onto a plane perpendicular (left side) and parallel (right side) to the membrane. ⟨*l*_*i*_⟩_p_ is the mean value (computed along P) of the characteristic size of the pore domain (PD). VSD stands for voltage sensor domain. **(c)**,**(i)**, Collapsed traces of the empirical,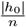 [kcal/atom], and theoretical, *ε* [kcal/atom], absolute atomic hydropathic energies.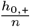 is the hydrophobic component, and 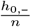 is the hydrophilic component of 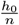, respectively. **(d)**,**(j)**, The interplay between the normalized statistical mechanical entropy 𝒮, and the pair of interaction ranges, {*ξ, ζ*}. **(e)**,**(k)**, Collapsed traces of ‘pointing extracellularly (ES)’ and ‘pointing intracellularly (IS)’ instances of the normalized membrane-vertical HDF amplitude, *h*_1,⊥_ [kcal×A /atom]. *h*_1,⊥_ is normalized over 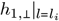. If 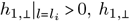 is labeled as a ‘pointing ES’ (↑) instance; otherwise, ‘pointing IS’ (↓). **(f)**,**(l)**, Nonextensivity of 𝒮 [kcal/(Θ×atom)], as revealed from its weak and strong dependence on the molecular radial size, *s* [Å], and the interaction range ratio,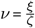, respectively. Note that 𝒮 is normalized over 𝒮_max_ (Eq. (18)). The critical inflection regime marks the interval from which all *l*_*i*_ values are drawn. Trace collapse procedures are described in SI S1.8.1.1.

To appreciate the statistical significance of this finding, we emphasize that our fitting algorithm had to converge approximately 121 × 690 = 83,490 times, where 121 is the total number of NaVCh structures considered and 690 gives the average number of investigated pore points per NaVCh structure, separated by a sampling distance of 0.1 Å (SI Tab. S2).

These results indicate that NaVCh atom packing physics can be compressed into the set of parameters {*ζ, ξ, K*} (Eqs. (2),(3)), in a pore-point-specific manner. Key to our understanding is that *ξ* and *ζ* are indirect measures of molecular attraction and repulsion, influencing molecular compression and expansion, respectively. Accordingly, the NaVCh molecule is conceptualized as a compressible, fluid-like material, whose mechanofunctional properties vary along its principal pore axis, approximated with the pore point path 𝒫. In turn, this implies that the atomic environment around each pore point requires a distinct statistical mechanical description or, equivalently, admits a local temperature linked to the PD hydropathic energy levels (SI S1.4).

The formation of interfacial geometries in compressible materials requires the spontaneous cancellation of interfacial tensions, manifested as minima in the underlying interfacial free energy [55], a principle that also applies to the spatial organization of NaVChs. Structurally, this prompts the emergence of the PD/VSD interface, denoted as *∂*ℬ _*i*_ (Eq. (6)), mediating two qualitatively different – in terms of their thermostability character – atomic environments. This insight enables the structural annotation of atoms, determining whether they belong to the PD or the VSDs, as *n K* serves as an order parameter distinguishing between two coexisting material phases (Figs. 2(b),(h), SI Figs. S5-S10(b), S12-S22(b)).

Under mean-field conditions (when *ξ* = *ζ*, see below Eq. (8)), *n*/*K* admits the standard critical exponent, *β* = 0.5 (SI Eqs. (S15),(S16)). Moreover, it can be readily shown that *β* is related to other standard critical exponents through a generalized version of Widom’s scaling law – whose standard form can be recovered when the characteristic renormalization scale matches the correlation length (represented by *l*_*i*_ and *ξ*, respectively) (SI S1.6.4).

##### 3.1.1.2 Pore domain intrinsic dimensionality

Dimensionality analysis reveals that prokaryotic and eukaryotic PDs exhibit a preference for flatness, as their intrinsic dimensions (Eq. (19)) are, on average, values only slightly greater than 2 (Figs. 3(a),(b)).

**Fig. 3.**
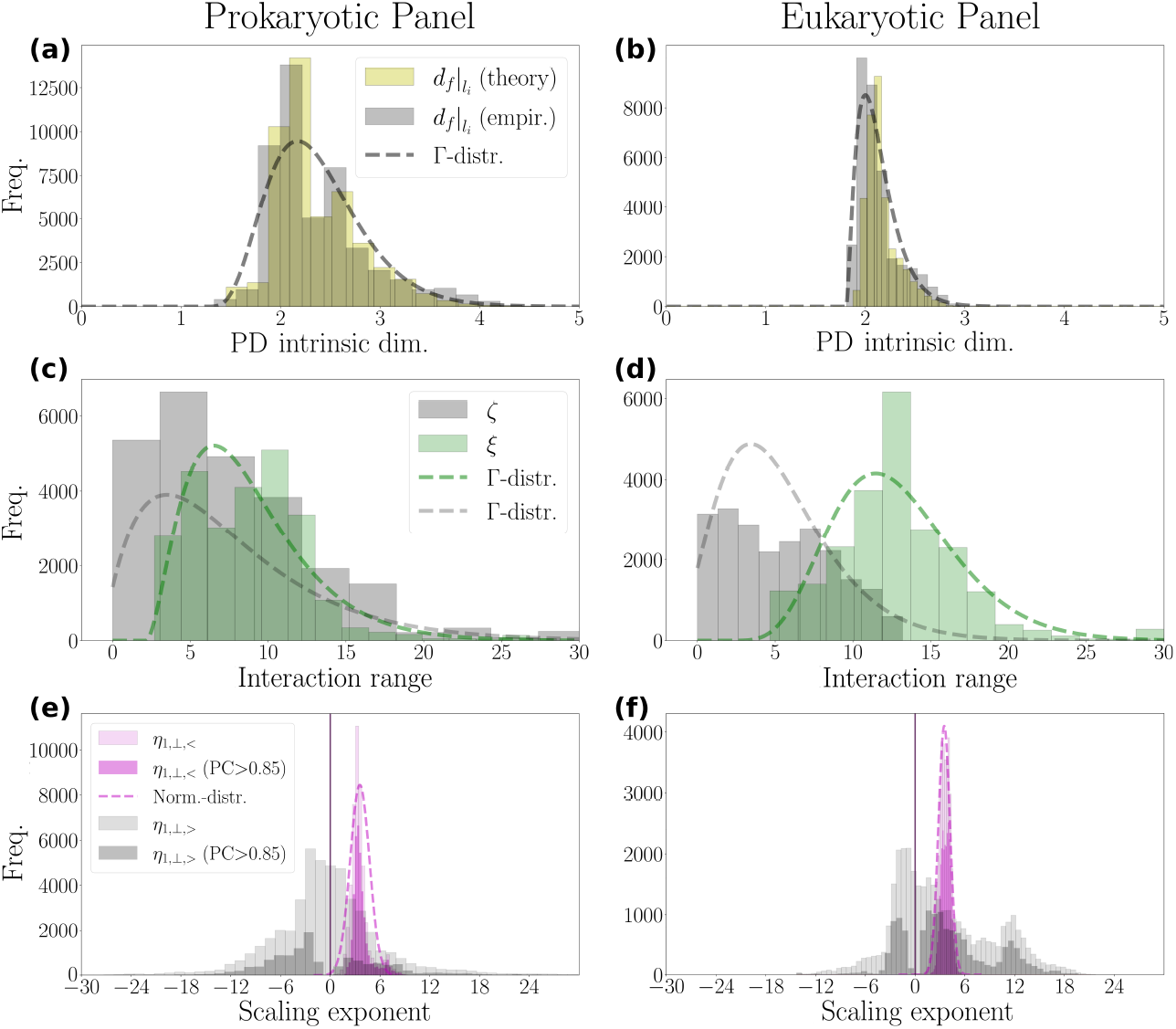
Intrinsic dimension of a pore domain and attractive-vs.-repulsive interaction range statistics. Statistical compilation of the pore domain (PD) intrinsic dimension and interaction range characteristics for 71 prokaryotic (subtypes: NaChBac, NaVAb, NaVMs, NaVAe, NaVRh) and 50 eukaryotic (subtypes: NaVPas, NaVEe1, NaV1-8, NaVEh) NaVCh atomic structures (SI Tab. S1). **(a)**,**(b)**, Histograms of the empirical and theoretical measures of the PD intrinsic dimension. The Γ-distribution mean value of the empirical intrinsic dimension measure is 2.4 and 2.1, with corresponding standard deviations of 0.5 and 0.2, for prokaryotic and eukaryotic PDs, respectively. **(c)**,**(d)**, Histograms of the attractive and repulsive interaction ranges given by *ζ* [Å] and *ξ* [Å], respectively. The Γ-distribution mean value of *ξ* is 8.8 Å for prokaryotes and 12.9 Å for eukaryotes, while the mean value of *ζ* is 7.5 Å for prokaryotes and 5.2 Å for eukaryotes. **(e)**,**(f)**, Histograms of the scaling exponents of the membrane-perpendicular dipole field component, *h*_1, ⊥_. The exponents *η*_1,⊥,<_ and *η*_1,⊥,>_ account for the scaling behavior of *h*_1,⊥_ over *l*-intervals covering the PD and the VSDs, respectively (Eq. (23)). They are computed according to procedures described in SI S1.8.3. PC stands for Pearson coefficient.

By tuning the intrinsic dimension of the PD near 2, PD sub-architecture is effectively mapped onto consecutive membrane-parallel planes, with roughness reciprocally related to 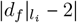, favoring a smooth cylindrical shape. As inferred from Eq. (20), this reflects a self-regulation mechanism that ensures the PD maintains its functional and structural integrity, despite the excess entropy generated by the tightening of the atomic environment, driven by hydrophilic deficiencies (or equivalently, a surplus of hydrophobicities) causing *ζ* → 0. Assuming *ξ* is fixed at a value well above zero, decreasing *ζ* effectively reduces interatomic distances, which, in turn, counteracts the expansion of the configuration space caused by excess entropy. Hence, by lowering its intrinsic dimension, the PD increases its spatial organization efficiency, as the likelihood for PD-forming atoms to adopt a nearly water-free, planar configuration increases.

Eukaryotic PDs exhibit greater efficiency in spatial organization, as evidenced by their smaller mean and standard deviation values for intrinsic dimensions compared to prokaryotic PDs (see Fig. 3 caption). This finding is consistent with the general trends reported in Ref. [56], where lower intrinsic dimensions were found to correlate with higher organismal complexity.

#### 3.1.2 Energy and entropy

##### 3.1.2.1 An exponentially decaying hydrophobic ‘force’ stabilizes the structure

As illustrated in Figs. 2(c),(i), a decrease in *n*-cluster size implies an increase in its average hydrophobicity, *h*_0,−_ /*N* (Eqs. (22),(24)). Specifically, *h* /*N* converges towards the value −0.5 kcal for *l* → *l*_0_ (decreasing scale index), denoting the hydropathic score that an individual hydrophobic atom is assigned with based on the Kapcha&Rossky hydropathic scale [50]. In contrast, the *n*-cluster average hydrophilicity, *h*_0,+_/*N*(Eqs. (22),(24)), tends to vanish for *l* → *l*_0_ (Figs. 2(c),(i)).

This trend showcases the entropically favorable arrangement where hydrophobicities ‘hide’ inside ℬ rather than expose on its surface, *∂*ℬ. As hydrophobic constituents fill the available space in ℬ, hydrophilic ones are displaced outwards, preferentially settling near *∂*ℬ. This redistribution creates an initial surplus of hydrophobic energy available to pore-lining constituents, as evidenced by the peak in the experimentally observed absolute hydropathic energy, |*h*_0_|/*N*, when *l* → *l*_0_ (Figs. 2(c),(i), SI Figs. S5-S10(c), S12-S22(c)).

However, as the *n*-cluster size grows (increasing scale index), the initial hydrophobic energy surplus is exponentially depleted at a rate governed by *ξ*, converted into stabilizing interactions that bind the nested ‘s together. Intriguingly, this would imply that interfacial adsorption effects do not substantially distort atom packing conditions (Eq. (22)). In turn, this would imply that NaVChs have evolved near a thermodynamic state where they mimic the spatial organization traits of larger, bulkier systems. This maximizes NaVCh mechanofunctional efficiency by allowing perturbations to be processed nearly adiabatically (SI S1.6.1). Our analysis supports this intuition, as shown by the good agreement between |*h*_0_|/*N* and *ε* [kcal/atoms] (Figs. 2(c),(i), SI Figs. 5-10(c), 12-22(c)).

Evidently, if the repulsive interaction range, *ζ*, grows too large, structural integrity could be compromised due to excessive interatomic spacing. To avoid that, the histogram of *ζ* is located closer to zero compared to that of *ξ* (Figs. 3(c),(d)). Also, in contrast to *ξ, ζ* accepts a smaller mean value for eukaryotic NaVChs compared to prokaryotic NaVChs (Fig. 3 caption), supporting that eukaryotic NaVChs, despite being more diverse – both structurally and functionally –, are spatially organized in a more efficient manner.

##### 3.1.2.2 Entropy constraints along the pore

Moving from the IS to the ES along 𝒫, we observe that the statistical mechanical entropy, 𝒮 (Eq. (17)), is smoothly up-regulated in both prokaryotes and eukaryotes (Figs. 2(d),(j), SI Figs. S5-S10(d), S12-S22(d)), as a result of the general trend that *ζ* surpasses *ξ* at the IS. Atomic environments at the IS and ES are thus subject to qualitatively entropic constraints, likely also experiencing asymmetric perturbation responses. To sustain such a delicate metastable equilibrium, even instantaneously, requires the contribution of external forces, since otherwise the structural integrity of the IS could be compromised. The observed nonextensivity, where the entropy is largely indifferent to changes in the radial molecular size while being slaved to the interaction range ratio, 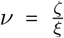 (Eq. (2), Figs. 2(f),(l) and SI Figs. S5-S10(f), S12-S22(f)), is likely the key property ensuring that the structure can quickly react to environmental changes and visit unconventional metastable states.

𝒮 maximizes in response to reduction of repulsive effects (*ζ* → 0), indicating a tightly organized, prevalently hydrophobic atomic environment (Eqs. (17),(20)). This is typically the case in the pore center, where the equivalent of a hydrophobic core exists, namely, a prevalently hydrophobic central cavity (e.g., see Fig. 1 from [40]). To resist structural disorder caused by excessive, hydrophobicity-induced configurational freedom, the PD reduces its characteristic size in order to decrease its intrinsic dimension (SI S2.2), which, in turn, guarantees greater efficiency in spatial organization.

#### 3.1.3 The PD/VSD structural transition as an order/disorder phase transition

Implications for the NaVCh functional architecture arising from the cancellation of interfacial tensions in the vicinity of the PD/VSD interface are best summarized by the membrane-perpendicular first-order hydropathic dipole field (HDF) amplitude, *h* [kcal× Å] (as explained in SI 1.5.2).

The sign of 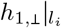 indicates whether the *h*_1,⊥_**ê**_⊥_-field induced by the PD at the current pore point is oriented extracellularly (↑) or intracellularly (↓) (Fig. 2 caption and SI 2.4.2, 2.5.2).

As shown in Figs. 2(e),(k) and SI Figs. S5-S10(e), S12-S22(e), distinguishing between *h*_1,⊥↑_, and *h*_1,⊥↓_, highlights the critical [30, 31] nature of the inflection point.

Specifically, *h*_1,⊥,↓_ and *h*_1,⊥,↑_ exhibit clear negative and positive peaks, respectively, for *l* ≈ *l*_i_, indicating that |*h*_1,⊥_ | is globally maximized. Implying that the rate of change of *h*_1, ⊥_ has stagnated, causing the interfacial free energy pair, {*∂h*_1,⊥↑_/, *∂l, ∂h*_1, ⊥↓_/, *∂l*} [kcal], to nearly vanish. This resembles a second-order phase transition, though smeared out by NaVCh finiteness.

Regardless of its orientation, |*h*_1, 1,⊥_| increments in a power-law fashion in both prokaryotic and eukaryotic PDs. The corresponding scaling exponents, *η*_1,⊥,<_ (Eq. (23)), are narrowly distributed with 3.2 ± 0.6 and 3.5 ± 1.3 (see Figs. 3(e) and (f), respectively). High Pearson coefficient values verify that these observations reflect a genuine self-similar increment behavior (e.g., see Fig. 3(c) from [36]). Microscopically, this necessitates that dipole-dipole hydropathic interaction networking radial atom neighbors are nonrandomly aligned [36], as random alignment would make self-similar incrementation of their cumulative index an extraordinary evolutionary accident.

The narrowness of the *η*_1, ⊥<,_ -distribution (Figs. 3(e),(f)), further necessitates a highly specialized nature for the water-mediated interactions established between the PD and the ion/water pore mixture. The PD operates under a narrow-banded dehydration protocol, through which ion selectivity can naturally emerge, as the radial alignment of dipoles around a pore point guarantees that water reorganization energies optimally dissipate into the atomic structure. Perturbation amplification is strictly unidirectional: perturbation amplitude attenuates inwards (towards the pore-lining interface, *∂*ℬ_0_), while amplifies outwards (towards the PD/VSD interface), irrespective of the current pore point. This implies the existence of a directed (or ordered) allosteric network inside the PD, where any pair of perturbed pore-lining constituents can potentially influence the same distant neighbor, unlike the other way around.

Once the characteristic PD size is surpassed (i.e., for *l* > *l*_*i*_), *h*_1, ⊥ ↑_, bends upwards, while *h*_1,_⊥_,↓_ bends downwards, thereafter exhibiting transient behaviors that cannot be described by a single scaling rule, as can be inferred by the broad, zero-centered *η*_1,⊥, >_ -exponent distribution (Figs. 3(e),(f)). |*h*_1,⊥_| may thus increment, decrement, or stagnate beyond the PD (note how percentile clouds increase for *l* >*l*_*i*_ in Figs. 2(e),(k) and SI Figs. S5-S10(e), S12-S22(e)). Given the high Pearson coefficient values associated with distributional *η*_1,⊥,>_-patterns on both sides of zero (Figs. 3(e),(f)), the observed diversification of the the scaling behavior of |*h*_1,⊥_| beyond the PD, supports that allosteric pathways coupling the PD to the VSDs are bidirectional: perturbations can be either attenuated or amplified from a VSD towards the PD/VSD interface in a pore-point-specific manner (SI 2.4.3, 2.5.3). This implies the multidirected (or disordered) nature of the allosteric network coupling the PD with the VSDs.

### 3.2 NaV1.7 mutational robustness in the context of human pain disease

Let us assume that the NaV1.7 molecule exhibits a uniform mutational robustness profile, implying that its ability to tolerate mutations is approximately the same across all of its constituent parts. Given that the PD/VSD interface is the most occupied molecular surface, it then follows that it is also a mutation hotspot, i.e., a geometric site where the likelihood of residue mutation is highest.

To scrutinize the validity of the above assumption, we examine mutation clustering in the NaV1.7 structure with the highest available resolution (PDB code: 7w9k). We consider all *SCN9A*-gene-related mutations found in gnomAD [23] and ClinVar [57] databases, yielding *N*_mut_ = 1705 events spanning three main categories: pathogenic (6.5%), non-pathogenic (11.3%), and variants of unclassified (or unknown) significance (VUS) (82.2%). The set of pathogenic mutations contains the subset of pain-disease-associated mutations (2.7%) and a substantial portion of paindisease-unrelated pathogenic mutations (3.8%). Pain-disease-associated mutations are categorized as either gain-of-function (GoF) (2.1%) or loss-of-function (LoF) (0.5%) (SI S1.9). Non-pathogenic mutations comprise two subsets: benign and neutral, with the latter consisting of carefully selected negative controls for human pain disease, sourced from Refs. [40, 58].

To probe mutation clustering relatively to the PD/VSD interface, we consider the dataset, {*l*_mut_ −*l*_*i*_}, where 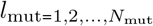 is the Euclidean distance between a pore point and the geometric center of the residue undergoing mutation (referred to as ‘structural location’). The triple vanishing of mean, mode, and median of {*l*_mut_ − *l*_*i*_} indicates that the likelihood for observing a mutation maximizes for *l* =*l*_*i*_ and, moreover, that the underlying mutation distribution is roughly bell-shaped (Fig. (4) insets), implying the uniformity of the mutational robustness profile.

**Fig. 4.**
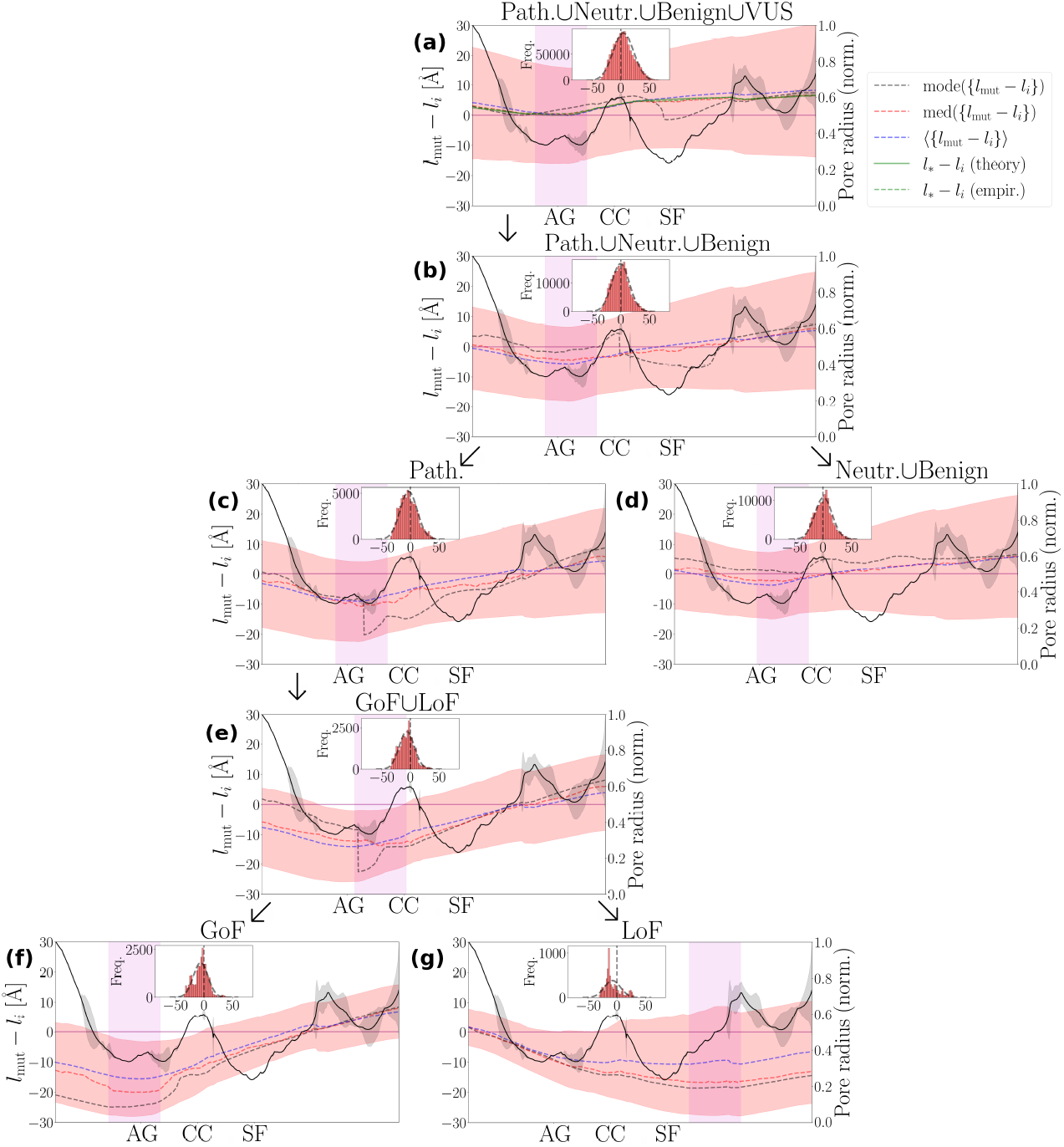
Distributional characteristics of mutations inside a NaV1.7 structure. We summarize the statistical properties of the {*l*_mut_ − *l*_*i*_} dataset in terms of the median med and the mean, ⟨⋅⟩, for pore points, **p** ∈ 𝒫. *l*_mut_ represents the Euclidean distance between **p** and the geometric center of the residue being mutated. We break down {*l*_mut_ − *l*_*i*_} to different subsets each of them corresponds to a different mutation set. Insets visualize the histogram of the {*l*_mut_− *l*_*i*_} dataset, i.e., the collapsed distribution incorporating contributions from all pore points. In **(a)**, we show the {*l*_mut_ −*l*_*i*_} -distribution of the parent set of all (i.e., Path.∪Benign∪Neutr.∪VUS) *SCN9A*-gene mutations. *l*_*_ identifies the structural location where the coin-flipping entropy is maximized. Note that if *n*_*L*_ ≈ *K, l*_*_ can be approximated with *l*_*i*,b_ (Eq. (8)). The empirical and theoretical instance of *l*_*_ is determined by the *l*-value for which *N/ N*_total_ =0.5 and *n /n*_*L*_ =0.5 is satisfied, respectively. The parent dataset is broken down to the following: **(b)**, pathogenic and non-pathogenic (Path.∪Benign∪Neutr.). **(c)**, pathogenic (Path). **(d)**, non-pathogenic (Benign Neutr.). **(e)**, Gain-of-function (GoF) and loss-of-function (LoF) (GoF∪LoF). **(f)**, GoF. **(g)**, LoF. Note that a negative and positive sign of *l*_mut_ − *l*_*i*_ indicate that the residue being mutated is likely residing in the PD and the VSDs, respectively. The areas highlighted in light magenta represent the pore regions where the median is minimized.

The uniform mutational robustness profile hypothesis for the NaV1.7 molecule is rejected across 𝒫, except at the AG pore region (Fig. 4(a)). While the mean and median of {*l*_mut_−*l*_*i*_} closely agree, and the mode oscillates around them within reasonable bounds (Fig. 4(a)), their deviation from zero reflects random genetic fluctuations governed by a flipping-coin maximum entropy principle.

Accordingly, the probabilities of a mutation occurring inside or outside ℬ are given by *P*_in_ = *N* /*N*_total_ ≈ *n*/*n*_*L*_ and *P*_out_ = 1 − *P*_in_, respectively. Maximizing the flipping-coin entropy, *H* ∶ = −*P*_in_ log_2_(*P*_in_) − *P*_out_ log_2_(*P*_out_), requires that *P*_in_ = *P*_out_ = *n*_∗_/*n*_*L*_ = 0.5 with *l*_∗_ denoting the *l*-value for which *n*_∗_ *n*_*L*_ 0.5 is satisfied.

As shown in Fig. 4(a), the line *l* − *l* coincides with the {*l*_mut_ − *l*_*i*_}-median, verifying that mutation clustering in the NaV1.7 is predominantly shaped by randomness. Across an evolutionary time-scale, this results in a bell-shaped mutation distribution approximately centered at 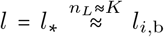 (Eq. (8)), ensuring that molecular diversity is explored inside and outside ℬ_∗_ in an unbiased manner. Because the range of attractive interactions prevails over the range of repulsive interactions (SI Fig. S23), the *∂*ℬ_∗_ -interface encapsulates the PD/VSD interface (Fig. 5(a)), acting as a buffer, dissipating mutation-induced perturbations and mitigating their impact on the PD.

**Fig. 5.**
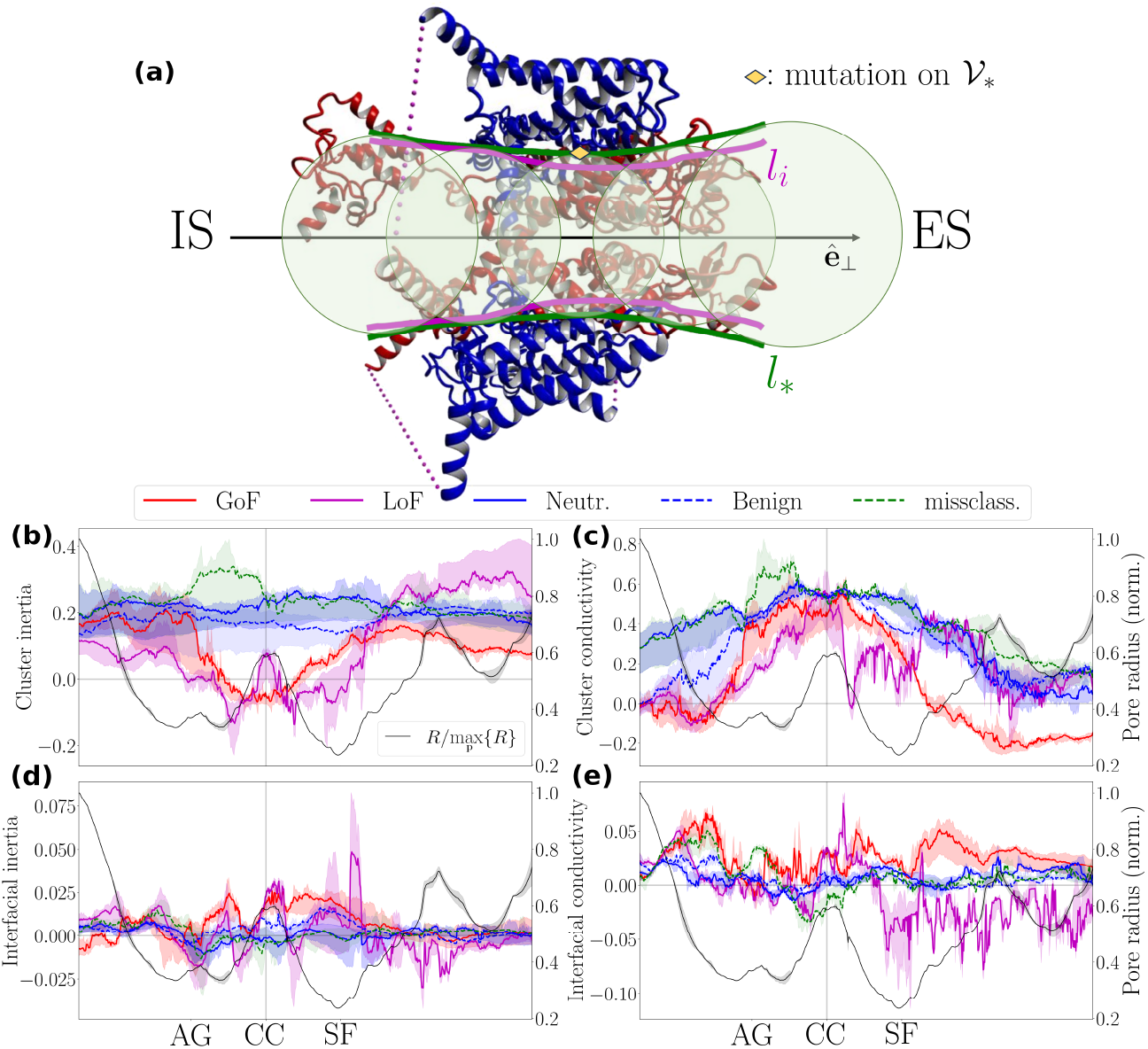
Inertia and conductivity profile of structural locations attracting pain-disease-associated mutations. **(a)**, Side view of a human NaV1.7 channel (PDB code: 7w9k). The PD and VSDs are illustrated in red and blue, respectively. For clarity, the *β* subunits are not shown. The dense arrangement of ℬ_*_-balls along 𝒫 creates the smooth cylindrical-like surface (in green), denoted as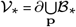. For comparison, the characteristic size of the PD, *l*_*i*_, is also illustrated (in magenta). Note that the inequality *l* > *l*_*_ arises from the atom packing condition *ν*< 1 (SI Fig. S23). At the AG pore region, where *ν↗* 1, *l*_*_ ≈ *l*_*i*_ implies *l*_*_ ≈ *l*_*i*,b_ with *n*_*L*_ ≈ *K* (see Eq. (8)). In **(b)** and **(c)** we statistically summarize the *n*-cluster inertias,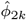, and the *n*-cluster conductivities,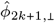, characterizing structural locations where the gain-of-function (GoF), loss-of-function (LoF), Neutral (Neutr.) and Bening mutations appear for *k* = 0, 1, … 5 **(d)** and **(e)** provide analogous information at an interface level: we statistically summarize the *∂n*-shell inertias,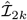, and the *∂n*-shell conductivities,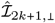,. Clouds around the index traces represent the min-max range of the underlying data points. The misclassified (missclass.) subset contains pain-disease-associated mutations systematically misclassified by the machine-learning algorithm (see Fig. 6(b)). The conductivity index derived from the first-order hydropathic dipole field is separately shown in SI Fig. S4, as the pair {*h*_1,+,⊥_, *h*_1,−,⊥_} satisfies (24) only partially (SI S2.3.1). The construction of the statistical summary indices 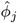 and 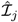 is described in SI S1.8.2. Molecular illustrations were generated using the Yasara software [86].

This is not the case at the AG pore region, where all the statistical indices considered above tend to vanish (Fig. 4(a)), promoting a uniform mutational robustness profile as *∂B* roughly coincides with the PD/VSD interface (Fig. 5(a)). Although the PD loses its protective buffer, it gains evolutionary momentum, allowing for greater exploration of molecular diversity. Also, mutations around the AG pore region are equally weighing on the PD and VSDs, preventing either from being disproportionately stressed.

Collapsing {*l*_mut_ − *l*_*i*_} into a histogram yields a roughly symmetric bell shape, well-described by the normal distribution, 4.46 ± 18.04 (inset in Fig. 4(a)). Excluding VUS, shifts the distribution to the left, with −0.47 ± 15.18 (Fig. 4(b)). Pathogenic and non-pathogenic mutation histograms admit −2.89 ± 14.9 and 0.9 ± 15.1 indicating their slight preference for the PD and the VSDs, respectively (Figs. 4(c),(d)). The pain-disease-associated mutation histogram is further left-shifted with −6.2 ± 13.0 (Fig. 4(e)), indicating a strong preference for the PD.

Distinguishing between GoF and LoF mutations reveals opposite statistical trends (Figs. 4(f),(g)). GoF mutations minimize their distance from the AG (Fig. 4(f)). On the other hand, LoF mutations converge towards the ES of the SF, where the extra-cellular funnel is formed (Fig. 4(g)). The molecular basis for GoF and LoF pain phenotypes can thus be underscored by qualitatively different perturbation modes. Positive *η*_1,⊥,>_ -exponents at the AG pore region favor GoF-mutation-triggered perturbations to amplify and propagate further into the VSDs (SI Fig. S23). On the other hand, negative *η*_1,⊥, >_ -exponents at the ES side of pore disfavor further amplification of LoF-mutation-triggered perturbations into the VSDs (SI Fig. S23), potentially causing them to be locally absorbed with detrimental impact on the PD.

#### 3.2.1 Violation of inertia and conductivity constraints underpins human pain disease at the molecular level

We hypothesize that pain-disease-associated mutations exploit interaction network imbalances at an interface level to perturb the *n*-cluster beneath.

To explore this hypothesis for the NaV1.7, we utilize the Decomposition ansatz (24) (SI S2.3), to derive the logarithmic composite susceptibility, *φ*_*j*_ (Eq. (29)), and combine it with the interfacial coupling strength, ℐ_*j*_ (Eq. (28)), in the form of the ratio 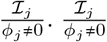 establishes an upper bound for the number of available perturbation modes encoded into a structural location (SI Eq. (S24)). Simply put,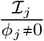 measures how many ways there are to destabilize an *n*-cluster, thus serving as a perturbation potential (i.e., as a measure of the sandpile slope) associated with the structural location under scrutiny.

A mechanical analogy rationalizing 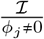is that of a maximum torque/force principle. When tightening a bolt, applying force at a stable far-end grip, where the inertia is minimum, maximizes torque. Analogously, residues distributed over strongly coupled interfaces covering clusters of vanishingly small inertia (probed with *φ*_2*k*_, *k* 0, 1, …, 5) are ideal mutagenesis sites; they can effortlessly perturb *n*-cluster rotation profile. A similar mechanism governs the perturbation of the *n*-cluster conductivity profile, characterized by the transverse odd-parity moments |*φ*_2*k* + 1, ⊥|_ (*k* = 0, 1, …, 5). A vanishingly small HDF indicates that the *n*-cluster conductivity becomes highly susceptible to perturbations, such that even minor surface fluctuations can induce disproportionately large reorganization in both amplitude and orientation of local dehydration forces.

Contrary to non-pathogenic mutations, pain-disease associated mutations prefer interfaces covering *n*-clusters whose inertia decreases, as we approach the CC pore region from left and right, and, additionally, at the SF and the AG pore regions (Fig. 5(b)). Specifically, the inertia of the *n*-clusters whose surface acts as a hotspot for GoF and LoF mutations vanishes twice and four times, respectively, at roughly symmetric locations along 𝒫. The arrangement and number of inertia zero-crossings may detrimentally perturb the NaV1.7 functional architecture via excess rotational DoF. Whether this leads to an increased or decreased pore-open probability and, consequently, results in a GoF- or LoF-like electrophysiological signature, is likely thresholded by the number of inertia zero-crossings, determining the tolerable excess rotational DoF before the gating cycle collapses. Targeting binding sites in the CC with anchoring molecules, e.g., local anesthetics, can thus offer a reasonable option for restoring inertia and mitigating the effects of the perturbation.

Additionally, GoF and LoF mutations prefer interfaces covering *n*-clusters whose conductivity is diminished past the AG pore region toward the IS (Fig. 5(c)). Also, GoF mutations cause a zero-crossing in conductivity at the ES of the SF pore region, while LoF mutations reduce conductivity at the IS of the SF pore region, without inducing a definitive sign change (Fig. 5(c)). Pain-disease-associated mutations may thus maximally perturb HDFs in the vicinity of the SF and AG, thereby potentially altering ion/water fluxes exactly at bottleneck pore regions on the left and right of the CC, where the pore radius is minimized.

At an interface level, inertia and conductivity amplitudes characterizing pain-disease-related mutations fluctuate stronger than those characterizing non-pathogenic mutations (Figs. 5(d),(e))). This suggests that GoF and LoF surface hotspots engage in substantially stronger coupling interactions with their environment compared to surfaces attracting non-pathogenic mutations. GoF and LoF surface hotspots are characterized by oppositely signed interfacial conductivities at the ES of the CC, suggesting that, once perturbed, these interfaces could generate oppositely directed ion/water flux responses (Figs. 5(e))).

#### 3.2.2 Verification of Mechanistic Rationalization via Machine Learning Experimentation

We introduce a machine learning strategy to differentiate between pathogenic and non-pathogenic NaV1.7 locations, both locally (i.e., at **p**) and globally (i.e., across 𝒫), based on 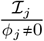 (SI S1.10). To avoid divergent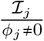 -values, we treat *φ*_*j*_ and ℐ_*j*_ as two separate feature inputs and we also investigate the significance of inertia and conductivity constraints separately, ending up with four feature inputs, {*φ*_2*k*_, I_2*k*_, *φ*_2*k*+1,⊥_, I_2*k*+1,⊥_}.

##### 3.2.2.1 Machine learning experiment I: pain-disease-associated vs. neutrals (PDB: 7w9k)

Following previous works [37, 58], we apply our algorithm to the well-balanced dataset of pain-disease-associated mutations and neutrals.

The local performance of the classifier, evaluated in terms of the area under the curve (AUC) and f1 scores, remains generally stable along 𝒫, indicating that the classifier can adapt well to local atom packing conditions (SI S2.6.1).

A yes/no variant classification scheme based on a linear threshold, is illustrated in Fig. 6(a). Its performance, evaluated in terms of the median of the AUC scores, reaches 0.85 (Fig. 6(e), Tab. 1).

**Fig. 6.**
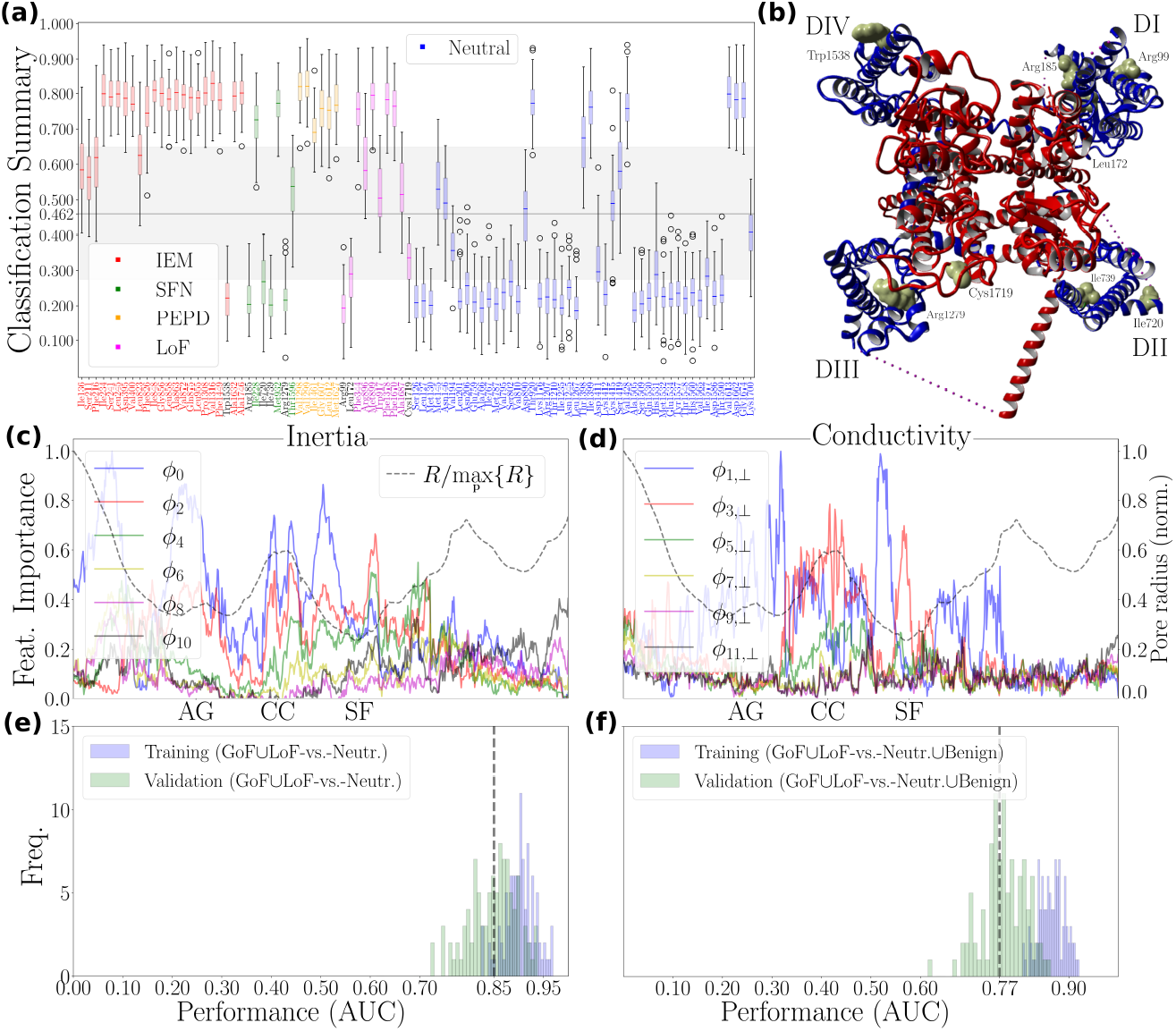
Machine-learning-assisted verification: pain-disease-associated mutation hotspots exhibit substantially distinct perturbation potential profile. **(a)**, Summary of a machine learning experiment evaluating the distinctiveness of the pain-disease-associated class (`class_ 0:` GoF LoF) in relation to the neutral class (`class 1`: Neutr.). Gain-of-function (GoF) subclass incorporates structural locations whose mutability is phenotypically associated with inherited erythromelalgia (IEM), small fiber neuropathy (SFN), and paroxysmal extreme pain disorder (PEPD) (SI Tab. S3). Loss-of-function (LoF) subclass incorporates structural locations whose mutability is phenotypically associated with insensitivity to pain (SI Tab. S3). Neutrals are structural locations whose mutability is not likely to be phenotypically associated with human pain disease, sourced from [37, 58]. The optimal threshold at 0.462 corresponds to the median of the best thresholds obtained through bootstrapping; each bootstrap sample of `class 0` probabilities yields a threshold that maximizes the f1-score. X-axis ticks indicate the location of a mutation event in the NaV1.7 structure. Note that x-axis ticks of misclassified pain-disease-associated mutations (i.e., those that do not pass the linear threshold) are highlighted in black. **(b)**, Top-view illustration of the human NaV1.7 channel (PDB code: 7w9k). For clarity, the *β*-subunits are not shown. The van der Waals surface of misclassified structural locations is highlighted in yellow. The PD and the VSDs are shown in red and blue color, respectively. In **(c)** and **(d)** we plot the importance score for the features *ϕ*_2*k*_ and *ϕ*_2*k*1_, *k =* 0, 1, …, 5, accounting for inertia and conductivity constraints, respectively. In **(e)** and **(f)** we illustrate the distribution of the final area under the curve (AUC) scores obtained from the final classification round for two different machine learning experiments, namely, the one summarized in (a) and the another one with `class _0:` GoF LoF vs. `class _1:` Neutr.∪Benign, respectively. Benign are structural locations whose mutability is not likely to be associated with disease (SI 1.9). Details concerning machine learning algorithm design and parameter selection can be found in SI 1.10. The Yasara software [86] was used for molecular illustrations.

*φ*_2*k*_-feature importance shows that maintaining higher-order interactions becomes increasingly important around the SF pore region. The importance of the second and fourth-order inertia features exceed that of the zero-th-order, suggesting that rotational and vibrational modes up to at least fourth order may influence ion selectivity (Fig. 6(c)). Corroboratively, the importance of higher-order interfacial inertia features rises sharply at the SF pore region, while, strikingly, the zero-th-order term, ℐ_0_, losses significance (SI Fig. S26(a)).

Additionally, at the SF pore region, higher-order conductivity features, particularly those of third and fifth order, become increasingly dominant, even surpassing the importance of first-order features (Fig. 6(d)). This indicates that volumetric, potentially asymmetric interactions may be crucial for the dehydration of sodium ions. In the ℐ_2*k* 1, ⊥_-feature domain, we observe similar interdependences at the ES entry point of the SF (SI Fig. S26(b)).

To clarify whether these results depend on the feature sign encoding information concerning the direction of the perturbation response (Eqs. 22,27), we repeat the machine learning experiment with the unsigned feature input, {| *ϕ*_2*k*_|, |*ℐ*_2*k*_|,*ϕ*_2*k* 1,⊥_ ℐ|, |*ℐ*_2*k* 1,⊥|_}. Classification results become only marginally worse (Tab. 1 vs. SI Tab. S5), suggesting that NaV1.7 functional architecture is affected by the violation of inertia and conductivity constraints shown in Fig. 5 in a largely direction-independent manner.

##### 3.2.2.2 Machine learning experiment II: pain-disease-associated vs. non-pathogenic (PDB: 7w9k)

To demonstrate that our findings are not biased by the specific choice of neutrals, we repeat our machine learning experiment, this time focusing on the non-pathogenic mutation subset containing both neutrals and benign mutations.

Obtained AUC scores admit a median of 0.77 (Fig. 6(f), Tab. 1). Despite the considerable class imbalance, as non-pathogenic mutations greatly outnumber pain-disease-associated ones with a class ratio as low as 0.19, the algorithm maintains both accuracy and stability, showing no detrimental effects (SI Figs. S24(c), S25(c)).

**Table 1.** Machine-learning experiments summary. The medians of the AUC and f1 scores were obtained during the final classification round (SI 1.10.2). The first and second number of each (⋅, ⋅) -pair are median values obtained during training and testing the final classification model, respectively. Inertia (iner.) and conductivity (cond.) constraints are probed by the feature inputs {*ϕ*_2*k*_, ℐ_2*k*_} and {*ϕ*_2*k*+1,⊥_, ℐ_2*k*+1,⊥_}, respectively. *SCN9A*-gene mutation datasets I, II, and III contain the classes {`class 0`: GoF∪LoF, `class _1:` Neutr.}, {`class_ 0:` GoF∪LoF, `class _1:`Neutr.∪Benign}, and {`class_0:` Path., `class_1:` Neutr.∪Benign}, respectively. GoF and LoF stand for gain-of-function and loss-of-function, respectively, and represent mutations associated with increased or diminished pain sensation (listed in SI Tab. S3). Neutrals (neutr.) are carefully selected human pain disease negative controls, sourced from [37, 58]. Benign are generally not expected to be associated with a disease phenotype. Note that the pathogenic (path.) class contains both pain-disease-associated and pain-disease-unrelated pathogenic mutations.

| Pert. modes | PDB: 7w9k, res.: 2.2 Å |  |  | PDB: 6j8j, res.: 3.2 Å |  |  |
| --- | --- | --- | --- | --- | --- | --- |
|  | iner./cond. | iner. | cond. | iner./cond. | iner. | cond. |
| AUC (dataset I) | (0.90, 0.85) | (0.89, 0.83) | (0.87, 0.84) | (0.90, 0.83) | (0.87, 0.82) | (0.87, 0.83) |
| f1 (dataset I) | (0.81, 0.78) | (0.78, 0.76) | (0.81, 0.80) | (0.80, 0.78) | (0.79, 0.76) | (0.80, 0.78) |
| AUC (dataset II) | (0.87, 0.77) | (0.80, 0.74) | (0.85, 0.77) | (0.84, 0.72) | (0.79, 0.70) | (0.79, 0.72) |
| f1 (dataset II) | (0.75, 0.72) | (0.76, 0.74) | (0.74, 0.72) | (0.79, 0.73) | (0.80, 0.78) | (0.74, 0.70) |
| AUC (dataset III) | (0.77, 0.63) | (0.71, 0.60) | (0.71, 0.64) | (0.79, 0.65) | (0.74, 0.64) | (0.71, 0.61) |
| f1 (dataset III) | (0.71, 0.62) | (0.67, 0.61) | (0.68, 0.64) | (0.72, 0.62) | (0.68, 0.61) | (0.67, 0.59) |

These findings reinforce that pain-disease-associated mutations appear in molecular neighborhoods of significantly different perturbation potential profile compared to non-pathogenic mutations.

##### 3.2.2.3 Machine learning experiment III: pathogenic vs. non-pathogenic (PDB: 7w9k)

Running our algorithm on the entire *SCN9A*-gene mutation dataset results in a drop of approximately 25% in performance, with the median AUC score dropping to 0.63 (Tab. 1). The reason for the observed drop is the inclusion of pain-disease-unrelated pathogenic mutations. The perturbation potential profile of structural locations whose mutability is associated with a disease phenotype distinct from painful or painless neuropathies, resembles that of non-pathogenic mutations – at least up to a first-order *ϕ*_*j*_ derivative. The level of confidence in this finding is high, as less than 6% of all pain-disease-unrelated pathogenic mutations are characterized as ‘Pathogenic/Likely pathogenic’ or ‘Likely pathogenic’.

Enhancing classifier performance would involve higher-order derivatives of *ϕ*_*j*_, alongside with refined scaling exponents calculation techniques as managing noise becomes progressively more challenging with higher derivative orders and greater distance from the pore. In fact, pain-disease-associated mutations affecting the VSDs are more likely to be misclassified (Fig. 6(b)). Four out of five misclassified GoF mutations are associated with an SFN phenotype and occur at VSD locations Arg185 (DI), Ile720 (DII), Ile739 (DII), and Arg1279 (DIII). The fifth misclassified GoF mutation, associated with an IEM phenotype, is found at the VSD location Trp1538 (DIV). Among the three misclassified LoF mutations, Arg99 (DI) and Leu172 (DI) are in the VSD, while Cys1719 (DIV) is in the extracellular loop connecting S5 and S6.

##### 3.2.2.4 Repeating machine learning experiments I-III using a lower-resolution structure (PDB: 6j8j)

To assess how lower resolution impacts algorithm performance, we repeat the analysis using a widely studied NaV1.7 structure with lower resolution (PDB code: 6j8j. Despite a slight drop in performance, the results remain largely consistent (Tab. 1 vs. SI Tab. S5). Given that both the high- and low-resolution structures likely represent an inactivated state of the channel, the observed decrease in performance is due to low-resolution limitations.

##### 3.2.2.5 Inertia-vs-conductivity: what matters more for NaV1.7 physiological functioning? (PDB: 7w9k, 6j8j)

Exchanging the feature inputs, {*ϕ*_2*k*_, ℐ_2*k*_, *ϕ*_2*k*+1,⊥_, ℐ_2*k*+1,⊥_}, with either {*ϕ*_2*k*_, ℐ_2*k*_} or *ϕ*_2*k*+1,⊥_, ℐ_2*k*+1,⊥_}, for the 7w9k or 6j8j structures, induces an algorithm’s performance drop that is always less then 10% (Tab. 1, SI Tab. S5). Additionally, the AUC scores achieved with {*ϕ*_2*k*_, ℐ_2*k*_} are close to those obtained with {*ϕ*_2*k* 1,⊥_, ℐ_2*k*+1,⊥_}. Together, these results suggest that the feature inputs {*ϕ*_2*k*_, ℐ_2*k*_} and {*ϕ*_2*k*+1,⊥_, ℐ_2*k*+1,⊥_} contain redundant information, implying that rotational and translational NaV1.7 DoF are coupled. The corresponding perturbations modes are thus likely intertwined through feedback, with molecule rotations around the pore influencing the flow of ions through the pore, and vice versa.

## 4 Discussion

The NaVCh functional architecture represents the latest product of an evolutionary process initiated nearly three billion years ago [59]. It embodies a rich metastable dynamics repertoire emerging from the instantaneous coordination of thousands of atoms assembled into several hundred amino acids and structurally organized into distinct domains around a central axis. Parsimonious models of the NaVCh functional architecture must thus rely on the derivation of evolutionary conserved laws that connect the microstructure (single atom) with the macrostructure (multi-domain molecule) in a manner that allows molecule functionality to emerge *a priori*, i.e., by cvirtue of the relationships established among these evolutionary conserved laws. Since an atom is separated from a domain by orders of magnitude in length scale, our quest finds its natural place in the RG framework clarifying how (i.e., through which scaling operations) one can transition from one molecular scale to the next one without losing the ability to reconstruct essential functional molecule characteristics. Since the evolutionary conserved laws are precisely defined by the underlying scaling operations, the two concepts are interchangeable and are both encompassed by the term ‘scaling law’. These considerations are summarized in the key analytical result of the RG flow equation (Eq. (30)), which is applied in a pore-point-specific manner, thus establishing a connection between the porous microenvironment and the surrounding molecular macroembedding context.

Importantly, we arrived at Eq. (30) by starting from the simplest possible equation of state (Eq. (2), describing the number of atoms within an infinitely thin shell *∂*ℬ, with a repulsion and stabilization term), that can rationally support a pore-forming macroenvironment where a PD is radially succeeded by four VSDs. To unbiasedly identify the PD/VSD interface, we focused on NaVCh shape characteristics: specifically, the sign change in the curvature of the ‘slow’ [60] state variable *n*, which marks a prominent inflection point. We are therefore confident that the logical thread connecting Eq. (2) with Eq. (30) is well-founded, suggesting that renormalizability is a fundamental property of NaVCh protein molecules. It is thus unsurprising that our theoretical framework can recapitulate Widom’s scaling law [61] under mean-field-like conditions and exactly at a molecular scale matching the inflection point (SI S1.7.4), suggesting that NaVChs belong to the same universality class as other multi-domain complex systems with some degree of fluidity, whose constituents interact via magnetic-like fields [62].

Experimental proof comes from the analysis of sufficient-resolution, full-atom NaVCh structures, shown to adhere to Eq. (2) within reasonable structural constraints – compromised only by extending the PD with a C-terminal domain while removing the VSDs (SI Fig. S7). The NaVCh structure can thus be treated as an ‘extended’ [63] (or ‘transient’ [64]) self-similar object whose intrinsic dimension changes continuously with molecular scale (Eq. (11)). A statistical mechanical description of ‘extended’ fractals is available within the framework of nonextensive statistical mechanics [48, 49], hallmarked by the *q*-deformed statistical mechanical entropy given by Eq. (17). Juxtaposing the notions of intrinsic dimension and statistical mechanical entropy suggests that the formation of the central cavity is a structural consequence of hydrophobicities being buried inside the NaVCh core over evolutionary timescales (3.1.1 and SI S2.2). Accordingly, the excess entropy generated by increasing core hydrophobicity is compensated by a reduction in intrinsic dimension, thereby preserving the structural integrity of the PD sub-architecture, as described by Eq. (19). This phenomenon is generally interpreted within the evolution-driven dimensionality reduction framework [65], which favors more efficient spatial organization in eukaryotic NaVChs, as verified both here across the NaVCh superfamily and across thousands of proteins in Ref. [56]. The physical basis of this phenomenon lies in the long-range nature of water-mediated interactions allowing spatially distant residues to influence one another [66–68]. Consistent with previous experimental findings reported in Refs. [51–53, 69], we found that the amplitude of the effective hydrophobic ‘force’ holding distant components together decays exponentially with increasing molecular scale, exhibiting a characteristic decay length, given by *ξ*, on the order of magnitude of 10 Å (Figs. 3(c),(d)).

Self-similarities imply long-range interactions, which in turn lead to synergistic constituent functioning via allostery [70]. The NaVCh field is well-positioned to begin contemplating the role of synergies within NaVCh molecules and their significance in both physiological function and mutation-perturbed disease-related contexts (e.g., see[71] and [72], respectively). Our results suggest that the *modus operandi* of the PD primarily relies on synergy built upon a narrow-width hydropathic dipole field exponent distribution (3.1.3). We argue that this evolutionary trend effectively reduces interaction modes for peeling off waters from a sodium ion while simultaneously increasing available pathways through which dehydration free energies can dissipate rapidly and deeply, i.e., in a nearly adiabatic manner, into the PD sub-architecture. Synergies are thus expected to arise from the unidirected (i.e., outwards-radiating) nature of allosteric interactions within the PD, enabling ‘collective’ responses to perturbations, as observed in molecular dynamics simulations [73]. Beyond the PD, the directionality of allosteric interactions is diversified, supporting the notion that VSD constituents may engage in asymmetric interactions both among themselves and with the PD. This renders analysis of VSD information processing capabilities challenging, calling for a more detailed future investigation that examines each VSD individually. Nevertheless, even with our current coarse-grained approach, we were able to infer that eukaryotic VSDs possess more specialized information-processing capabilities compared to their prokaryotic counterparts, enabling differential perturbation processing at the ES and IS (SI S2.5.3).

Our understanding of the molecular basis of human pain disease has been built upon painstaking experimental efforts, primarily relying on cell-based electrophysiology assays that screen single-amino-acid mutations in the human NaV1.7 molecule (for a list, see SI Tab. S3). Resolving the structure-function relationship riddle for the human NaV1.7 molecule in the context of human pain disease seems like assembling an impossible-to-ever-complete puzzle: each puzzle piece encodes only a tiny fraction of biophysical and neurophysiological possibilities. To demonstrate how our theoretical framework can simplify this task, we mapped all *SCN9A* gene mutations onto a wild-type NaV1.7 molecule and attempted to explain the observed mutation clustering patterns (3.2). We identified the most frequently mutated interface and showed that it becomes indistinguishable from the PD/VSD interface around the AG, where mean-field conditions are established as *q↗* 2. This finding demonstrates that the NaV1.7 mutation landscape has not emerged due to pure chance, but rather it is subject to NaV1.7 shape constraints that can vary along the pore, effectively captured by *q*. In fact, a tendency towards a more globular-like shape lowers the risk of experiencing a detrimental perturbation as the PD/VSD interface becomes the primary mutagenesis site and mutations tend to land between the PD and VSD, thereby sparing either domain from direct damage. Pain-disease-associated mutations appearing at the AG in the PD most likely result in a GoF phenotype, consistent with the mutation clustering patterns reported in Ref. [74]. This pattern is reversed for mutations linked to a LoF pain phenotype (Fig. 4, panel (g) vs. (f)); however, the limited number of available LoF pain mutations precludes any definitive conclusions. Nevertheless, when viewed through the RG lens, these observations suggest that the primary molecular distinction between GoF and LoF pain phenotypes lies in whether mutation-induced perturbation shocks are more likely to be distantly propagated or locally absorbed, as determined by the scaling properties of VSD-induced hydropathic fields (3.2 and SI Fig. S23).

Just as adding a grain of sand to a steep slope can trigger a cascading avalanche, mutations can exploit allosteric shortcuts [75] to augment NaVCh DoF, exerting a substantial destabilizing effect on the functional architecture. We conceive the NaVCh functional architecture as a hierarchically organized mechanical system centered around a principal pore axis, where hinges [76–79] are arranged in nested levels – smaller sub-hinges exist within or regulate parts of larger hinges – enabling multi-scale coordinated motion. Mechanistically, this implies a hierarchy among residues, with some more likely than others to significantly alter channel conductivity and inertia upon mutation-induced perturbation, analogous to how a force applied farther from a hinge generates a larger rotational effect about the axis, which can, in turn, induce large ion/water fluxes along the axis. An in-depth investigation of this idea was undertaken in accordance with the analytical procedures detailed in 2.1.4. The explanatory strength of this approach is illustrated in Fig. 5 and further verified via a metalearning algorithm, i.e., a machine learning paradigm that learns globally (i.e., across the pore) from prior, local (i.e., pore-point-specific) learning processes [80]. Our algorithm achieved state-of-the-art AUC scores (compared to previous efforts [37, 58, 81]), demonstrating that a standard support vector machine classifier suffices to learn the biomechanical constraints that differentiate pain-disease-associated mutation hotspots from any other benign and/or neutral site. However, it yields only mediocre results when applied to the entire dataset of pathogenic versus non-pathogenic *SCN9A* gene mutations. We argue that this is primarily due to low-resolution artifacts, which affect feature observability, and only secondarily due to the inefficacy of the features themselves.

In summary, relaxing inertia constraints that ‘lock’ the transmembrane helical segments around the CC in a ‘screwed-in’ state is a key perturbation mode through which GoF electrophysiological signatures can be generated in the context of human pain disease. This also applies to LoF electrophysiological signatures but in a more spatially extended form: enhancing rotational freedom at sites along the pore other than the CC can diminish open-state probability. It is hence reasonable to consider that the general mechanistic principle for shifting from a GoF-associated electrophysiological phenotype regime towards a LoF one is through desynchronization. Weakly desynchronizing rotational motions of transmembrane helical segments augments channel activation dynamics by increasing PD/VSD coupling configurations, ultimately increasing the probability of the open state. Strong desynchronization, however, has the opposite effect. Different parts of the transmembrane helical segments start to behave as asynchronous rotors, effectively decoupling the PD from VSD, thereby diminishing open-state probability. Drugging NaV1.7’s CC to correct the effects of GoF mutations by restoring inertial dynamics could therefore serve as an experimental validation strategy. Dismantling orientation and amplitude constraints of HDFs along the extra-cellular funnel towards the SF is an equally important perturbation mode. Our data suggest that whether a mutation results in gain-of-function (GoF) or loss-of-function (LoF) may depend on the orientation of dehydration forces exerted upon incoming ions (Figs. 5(c),(d)), a hypothesis that can be tested through molecular dynamics simulations. How these predictions will generalize to larger datasets of GoF versus LoF mutations remains, however, to be seen.

Several limitations apply to our work. First, the structure dataset considered here does not account for the full configuration repertoire that a NaVCh molecule can adopt. Most NaVCh of eukaryotic origin represent snapshots of an inactivated state. Second, we acknowledge that the high-resolution (< 3 Å) NaV1.7 structures with PDB codes 8s9b [82], 8xmm [83], and 8thh [84] were not available at the time of structure dataset assembly and are therefore not included in the present analysis. Third, misclassification of VSD hotspots remains an inherent challenge, which could potentially be mitigated by either partitioning the flow into four subflows, each sampling a single VSD, or by initiating RG flows along axes that traverse each VSD individually. As long as Eq. (2) is satisfied, the latter option introduces an axis translation, shifting the perspective from the central pore axis to peripheral VSD-centered axes simulating the direction along which gating charges move. Lastly, our work does not address dynamic NaVCh aspects, which, in turn, further limits our understanding of hydropathic interaction network intensification characteristics in the VSDs.

As a kind of Swiss knife that can cut through protein structure complexity, the RG offers a coarse-grained yet not oversimplified description of key mechanistic NaVCh aspects. On this basis, we investigated how machine learning paradigms that can efficiently (i.e., with minimal computational costs) and transparently (i.e., with maximal explainability) deliver biomedical insights for the pain clinic while enhancing our understanding of physicochemical NaVCh properties. Because our procedures are completely general, they could provide a foundation for analyzing various gene-protein relationships in a pathophysiological context, particularly for large globular or pore-forming systems being hierarchically organized around a principal point set.

## Supplementary information

This article includes supplementary information (SI), which can be accessed in the accompanying PDF file: SI RGNaVChPain.pdf.

## Acknowledgements

The authors would like to sincerely acknowledge the helpful discussions and support of M.Sc. Vishal Sudha Bhagavath Eswaran and Dr. Annette Lischka. Their contributions in identifying pain-disease-associated variants and providing valuable insights during discussions have been greatly appreciated and have contributed to the development of this study.

## Declarations

- Funding: This work was funded by the Deutsche Forschungsgemeinschaft (DFG, German Research Foundation 363055819/GRK2415 Mechanobiology of 3D epithelial tissues (ME3T) to A.L.; 368482240/GRK2416, MultiSenses-MultiScales to A.L., LA2740/6-1 to A.L.).
- Conflict of interest/Competing interests: The authors declare no competing interests.
- Ethics approval and consent to participate: do not apply.
- Consent for publication: do not apply.
- Data availability:
  – The data generated from analyzing 71 prokaryotic NaVCh structures are available at: https://doi.org/10.5281/zenodo.14617204
  – The data generated from analyzing 50 eukaryotic NaVCh structures and machine learning experiments in the context of human pain disease are available at: https://doi.org/10.5281/zenodo.14628099
- Materials availability: do not apply
- Code availability: The code used throughout is available at: https://github.com/mnxenakis/NaVCh_Scaling
- Author contributions: M.N.X. designed the study, conceived the mathematical structure, wrote the code, analyzed the data, and prepared the manuscript. A.L. codesigned the study, contributed to the interpretation of the findings, assisted with data analysis, reviewed the manuscript, and secured the funding.

