## Supplementary material for "Learning molecular traits of human pain disease via voltage-gated sodium channel structure renormalization": Suppl. Info.

#### S1 Supplementary Methods

##### S1.1 NaVCh structure collection

We searched in the Protein Bank Europe [1] by using the following keywords:

> Text: 'voltage-gated sodium channel'
> AND
> Text: 'X', where X ← NaChBac, NaVAb, NaVMs, NaVRh, NaVCt, NaVAe, NaV1-8, NaVEh, NaVEe1, NaVPas
> Resolution: smaller than or equal to 4 Å

The collection of voltage-gated sodium channel (NaVCh) structures is listed in Tab. S1.

##### S1.2 NaVCh structure preprocessing

###### S1.2.1 Cleaning

Removal of heteroatoms was performed in YASARA software [2] (YASARA).

**Table S1**: Collected NaVCh protein structures. With asterisk * we denote pore-only structures.

| Prokaryotes | | | Eukaryotes | | |
|---|---|---|---|---|---|
| subtype (*species*) | PDB code | resolution [Å] | subtype (*species*) | PDB code | resolution [Å] |
| NaChBac (*Bacillus Halodurans*) | 6vwx | 3.1 | NaVPas (*Periplaneta Americana*) | 6a95 | 2.6 |
| | 6vx3 | 3.7 | NaVEe1 (*Electrophorous Electricus*) | 5xsy | 4.0 |
| | 6vxo | 3.7 | NaVEh (*Emiliania Huxleyi*) | 7x5v | 2.83 |
| | 6w6o | 3.2 | NaV1.1 (*Homo Sapiens*) | 7dtd | 3.3 |
| NaVAb (*Arcobacter butzleri*) | 3rvy | 2.7 | NaV1.2 (*Homo Sapiens*) | 6j8e | 3.0 |
| | 3rvz | 2.8 | NaV1.3 (*Homo Sapiens*) | 7w77 | 3.3 |
| | 3rw0 | 2.95 | | 7w7f | 3.3 |
| | 4kew | 3.21 | NaV1.4 (*Homo Sapiens*) | 6agf | 3.2 |
| | 5kmh | 3.2 | NaV1.5 (*Homo Sapiens*) | 6lqa | 3.3 |
| | 5vb2 | 3.2 | | 7dtc | 3.3 |
| | 5vb8 | 2.85 | NaV1.5 (*Rattus Norvegicus*) | 6uz3 | 3.5 |
| | 5yua | 2.8 | | 6uz0 | 3.24 |
| | 5yub | 3.4 | | 7fbs | 3.4 |
| | 5yuc | 3.4 | | 7k18 | 3.3 |
| | 6c1e | 2.86 | | 8f6p | 3.2 |
| | 6c1m | 2.52 | NaV1.6 (*Homo Sapiens*) | 8fhd | 3.1 |
| | 6mvv | 2.9 | | 8gz1 | 3.4 |
| | 6mvw | 3.2 | | 8gz2 | 3.3 |
| | 6mvx | 2.46 | NaV1.7 (*Homo Sapiens*) | 5ek0 | 3.53 |
| | 6mvy | 3.0 | | 6j8g | 3.2 |
| | 6mwa | 2.4 | | 6j8h | 3.2 |
| | 6mwb | 2.6 | | 6j8i | 3.2 |
| | 6mwd | 2.33 | | 6j8j | 3.2 |
| | 6mwg | 2.5 | | 6ntq | 3.6 |
| | 6p6w | 4.0 | | 7w9k | 2.2 |
| | 6p6x | 2.75 | | 7w9l | 3.5 |
| | 6p6y | 2.89 | | 7w9m | 3.0 |
| | 7k48 | 3.6 | | 7w9p | 2.9 |
| | 8diz | 2.75 | | 7w9t | 3.0 |
| | 8dj0 | 2.7 | | 7xm9 | 3.0 |
| | 8dj1 | 3.1 | | 7xmf | 3.07 |
| | 8h9o | 3.3 | | 7xmg | 3.09 |
| | 8h9w | 2.7 | | 7xve | 2.7 |
| | 8h9x | 3.4 | | 7xvf | 2.8 |
| | 8h9y | 3.4 | | 8i5b | 2.7 |

#### Table S1 (continuation)

| Prokaryotes | | | Eukaryotes | | |
|---|---|---|---|---|---|
| subtype (*species*) | PDB code | resolution [Å] | subtype (*species*) | PDB code | resolution [Å] |
| NaVAb (*Arcobacter butzleri*) (continuation) | 8ha1 | 3.5 | NaV1.7 (*Homo Sapiens*) (continuation) | 8g1a | 2.8 |
| | 8ha2 | 3.3 | | 8i5g | 2.7 |
| NaVAe (*Alkalilimnicola Ehrlichi*) | 4lto | 3.46 | | 8i5x | 2.9 |
| | 4ltp | 3.8 | | 8i5y | 2.6 |
| | 5hj8 | 3.7 | | 8j4f | 3.0 |
| | 5hk7 | 2.95 | | 6nt3 | 3.4 |
| | 5hkd | 3.8 | | 6nt4 | 3.5 |
| | 5iwn | 3.75 | | 8f0p | 2.2 |
| | 5iwo | 3.33 | | 8f0q | 2.5 |
| NaVMs (*Magnetococcus marinus MC-1*) | 3zjz* | 2.92 | | 8f0r | 2.9 |
| | 4cbc* | 2.66 | | 8f0s | 3.1 |
| | 4oxs* | 2.80 | NaV1.8 (*Homo Sapiens*) | 7we4 | 2.7 |
| | 4p2z* | 3.08 | | 7wel | 3.2 |
| | 4p9o* | 2.89 | | 7wfr | 3.0 |
| | 4p9p* | 2.91 | | 7wfw | 3.1 |
| | 4p30* | 3.31 | | | |
| | 4pa3* | 3.25 | | | |
| | 4pa4* | 3.02 | | | |
| | 4pa6* | 3.36 | | | |
| | 4pa7* | 3.02 | | | |
| | 4pa9* | 3.43 | | | |
| | 4x8a* | 3.02 | | | |
| | 4x88* | 3.5 | | | |
| | 4x89* | 2.62 | | | |
| | 5hvd | 2.6 | | | |
| | 5hvx | 2.45 | | | |
| | 6sx5 | 2.2 | | | |
| | 6sx7 | 2.5 | | | |
| | 6sxc | 2.5 | | | |
| | 6sxe | 2.6 | | | |
| | 6sxf | 2.84 | | | |
| | 6sxg | 2.4 | | | |
| | 6yz0 | 2.3 | | | |
| | 6yz2 | 2.2 | | | |
| | 6z8c | 3.2 | | | |
| NaVRh (*Alpha Proteobacterium HIMB114*) | 4dxw | 3.05 | | | |

##### S1.2.2 Protonating

Addition of hydrogen atoms was performed using the Reduce software [3].

##### S1.2.3 Orienting

Using the VMD software [4], we computed the molecular principal pore axis, i.e., the axis passing through the pore and perpendicular to the membrane surface, and aligned it with the $z$-axis of the global coordinate system $(\hat{\mathbf{e}}_x, \hat{\mathbf{e}}_y, \hat{\mathbf{e}}_z)$. This procedure automatically rendered an $xy$-plane as membrane-parallel.

The molecular mass center, $\mathbf{c} = (\mathrm{c}_x, \mathrm{c}_y, \mathrm{c}_z) = \frac{1}{N\mathrm{total}} \sum_{\mathbf{a} \in \mathcal{A}} \mathbf{a}$, where $\mathbf{a} = (\mathrm{a}_x, \mathrm{a}_y, \mathrm{a}_z) \in \mathcal{A} \subset \mathbb{R}^3$ is a three-dimensional atom coordinate, was set to coincide with the coordinate system origin. $N_{\mathrm{total}} = \mathrm{card}(\mathcal{A})$ is the total number of atoms in the 'clean', protonated structure.

To emphasize its relationship with the membrane surface, the $z$-coordinate is referred to as the $\perp$-coordinate in the Main Text and thereafter, i.e., $\hat{\mathbf{e}}_\perp \equiv \hat{\mathbf{e}}_z$.

##### S1.2.4 Navigation through the pore environment

We call the HOLE routine [5] $N_{\mathrm{HOLE}} = 50$ times with parameter values shown in Tab. S2.

Once the $k$-th HOLE call returns, preliminary pore points, $\mathbf{q}_{k,j} = (\mathrm{q}_{x,k,j}, \mathrm{q}_{y,k,j}, j\Delta z)$, are collected, where $j = 0, 1, \ldots$ is an index for the current pore point where the HOLE routine is operating. The parameter $\Delta z$ [Å] determines the distance between consecutive membrane-parallel planes where pore points are scattered (Tab. S2).

Assuming that the scattering of $(\mathrm{q}_{x,k,j}, \mathrm{q}_{y,k,j})$ on the $j$-th membrane-parallel plane is – or at least tends to be as $N_{\mathrm{HOLE}} \to \infty$ – random, the expected pore point location is estimated by:

$$\mathbf{p}_j = (\overbrace{\frac{1}{N_{\mathrm{HOLE}}} \sum_{k=1}^{N_{\mathrm{HOLE}}} \mathrm{q}_{x,k,j}}^{=\mathrm{p}_{x,j}}, \overbrace{\frac{1}{N_{\mathrm{HOLE}}} \sum_{k=1}^{N_{\mathrm{HOLE}}} \mathrm{q}_{y,k,j}}^{=\mathrm{p}_{y,j}}, \underbrace{j\Delta z}_{=\mathrm{p}_{\perp,j}}). \tag{S1}$$

A statistical estimation of the pore radius is obtained by:

$$R_j = \frac{1}{N_{\mathrm{HOLE}}} \sum_{k=1}^{N_{\mathrm{HOLE}}} r_{k,j} \quad [\text{Å}], \tag{S2}$$

where $r_{k,j}$ is the pore radius corresponding to $\mathbf{q}_{k,j}$ returned by the HOLE routine.

The standard deviation in $R_j$ serves as a pore structure uncertainty measure:

$$\sigma_{R_j} = \sqrt{\frac{1}{N_{\mathrm{HOLE}}} \sum_{k=1}^{N_{\mathrm{HOLE}}} (r_{k,j} - R_j)^2} \quad [\text{Å}]. \tag{S3}$$

The radius of the smallest $\mathbf{p}_j$-centered ball that contains all $N_{\text{total}}$ atoms including their van der Waals radii is given by:

$$L_j = \max_{\mathbf{a}\in\mathcal{A}}\{\|\mathbf{a}-\mathbf{p}_j\| + \text{vdW}_{\text{a}}\} \quad [\text{Å}], \tag{S4}$$

where $\text{vdW}_{\text{a}}$ [Å] represents the van der Walls radius of the **a**-atom (Tab. S2). (S4) leads to the following definition for the radial molecular size about $\mathbf{p}_j$:

$$\begin{aligned} & s_j := L_j - \bar{R}_j \quad [\text{Å}] \\ & \text{with} \quad \bar{R}_j = \min_{\mathbf{a}\in\mathcal{A}}\{\|\mathbf{a}-\mathbf{p}_j\|\} \quad [\text{Å}], \end{aligned} \tag{S5}$$

where $\bar{R}_j$ is the smallest Euclidean distance between $\mathcal{A}$ and $\mathbf{p}_j$.

Elsewhere, the subscript '$j$' is omitted for clarity to avoid unnecessary complexity in the notation. The subscript '$j$' appearing here should not be confused with the subscript '$j$' appearing in the Main Text and in the subsequent discussion, where it specifically denotes the dimension of the space within which water-mediated interactions are probed. In the Main Text and thereafter, $R$ is referred to as the pore radius.

#### S1.3 Tight atom packing: the ($\zeta \to 0$)-case for $n$

Using $\lim_{y\to 0}(1+yx)^{-1/y} = \exp(-x)$, $x \in \mathbb{R}$, we obtain for $n$:

$$\begin{aligned} & n_{\zeta\to 0} = K\exp\left(-\frac{\varepsilon}{\varepsilon_{i,\zeta\to 0}}\right) \\ & \text{with} \quad \varepsilon_{i,\zeta\to 0} = \varepsilon_0\exp(-l_{i,\zeta\to 0}/\xi) \quad [\text{kcal/atom}], \quad l_{i,\zeta\to 0} = \lim_{\zeta\to 0} l_i = \xi\ln(\ln(K/n_0)) + l_0 \quad [\text{Å}], \end{aligned} \tag{S6}$$

where $\varepsilon$ [kcal/atom] is given by Main Text Eq. (4) and $\ln(K/n_0) = \lim_{\zeta\to 0}\left(\frac{(n_0/K)^{-\nu}-1}{\nu}\right)$.

In the growth modeling literature, (S6) is known as the Gompertz model (for a comprehensive review see Ref. [6]).

In Fig. S1, we illustrate that while keeping parameters $\xi$ and $K$ fixed, taking $\zeta \to 0$, translocates the inflection point to the left. This event signifies the tightening of the atomic environment as the radial atom distribution is displaced towards the left while also the atom packing rate, $1/\zeta$, diverges, causing $n_{\zeta\to 0}$ to approach $K$ 'sooner' than $n$. In fact, as shown in Fig. S1, $n_{\zeta\to 0}$ works as an upper bound for $n$.

In Fig. S1 inset, we depict the intrinsic dimension of a pore domain (PD), $d_f|_{l_i}$ (Main Text Eqs. (19),(20)). We notice how $d_f|_{l_i}$ increases monotonically for increasing the nonextensivity index, $q$, controlled by $\zeta$.

#### S1.4 Hydropathicity and temperature: different faces of the same coin?

The functional form of the $p_\alpha$-probability is deduced by maximizing the $\mathcal{S}$-entropy (Main Text Eq. (17)) under the constraints:

$$\begin{cases} \sum_\alpha p_\alpha = 1 \quad \text{(Main Text Eq. (13))} \\ \sum_\alpha \pi_\alpha \varepsilon_\alpha = \overbrace{\frac{\Xi}{C\Delta l_\alpha}}^{=:\rho} \varepsilon_i \quad \text{[kcal/atom]} \quad \text{(Main Text Eq. (15))} \end{cases}, \tag{S7}$$

where $\rho$ is introduced solely for the reader's convenience, as it improves the compactness of the formulation.

Following [7, 8], we proceed by formulating the variational problem:

$$\begin{aligned} &\frac{\delta}{\delta p_\alpha}\Big[\frac{\mathcal{S}}{k} - \mathcal{L}_1 \sum_{\alpha^*} p_{\alpha^*} - \mathcal{L}_2 \sum_{\alpha^*} \pi_{\alpha^*}\varepsilon_{\alpha^*}\Big] = 0 \\ &\text{with} \quad \mathcal{L}_2 \equiv \frac{1}{kT}, \end{aligned} \tag{S8}$$

where $\mathcal{L}_1$ and $\mathcal{L}_2$ are Lagrange multipliers. $T$ [$\Theta$] corresponds to the physical temperature that a molecular thermometer would measure if placed at a pore point, $\mathbf{p}$. $k$ [kcal/($\Theta\times$ atom)] is the equivalent of Boltzmann's constant.

From (S8), and following the procedures described in [8], we arrive at:

$$\begin{cases} p_\alpha = \frac{\left(1+(1-q)\frac{\varepsilon_\alpha}{\varepsilon_i}\right)^{-1/(1-q)}}{Z_p} \\ T = \frac{1-(1-q)\rho}{kZ_\pi}\varepsilon_i > 0 \quad [\Theta], \end{cases} \tag{S9}$$

where $\mathcal{L}_1$ has been eliminated from $p_\alpha$ via factorization.

(S9) indicates that $T$ depends on the unsigned hydropathic energy stored in a PD-forming atom, $\varepsilon_i$ [kcal/atom], suggesting a thermostatic role for the PD sub-architecture. To make this point even clearer, notice that for $\zeta \to 0$ (i.e., $q \to 1$), we have:

$$\begin{cases} p_{\alpha,\zeta\to 0} = \exp(-\frac{\varepsilon_\alpha}{\varepsilon_{i,\zeta\to 0}})/Z_{p_{\alpha,\zeta\to 0}}, \quad Z_{p_{\alpha,\zeta\to 0}} = \sum_\alpha \exp(-\frac{\varepsilon_\alpha}{\varepsilon_{i,\zeta\to 0}}) \\ T_{\zeta\to 0} = \varepsilon_{i,\zeta\to 0}/k \quad [\Theta] \end{cases}, \tag{S10}$$

indicating how $\varepsilon_{i,\zeta\to 0}$ (see Eq. (S6)) is now exclusively governing the thermal characteristics of the local atomic environment.

#### S1.5 Molecular degrees of freedom emerging from water-mediated interactions

We interpret Eq. (21) appearing in the Main Text in the context of NaVCh DoF expressed around and along $\mathcal{P}$. The former pertains to rotational, and the latter to translational molecular phenomena.

##### S1.5.1 Rotational degrees of freedom emerging from water-mediated interactions ($j = 2k$)

$h_{2k}$ informs about an $n$-cluster's resistance in changes of its hydropathic composition ($k = 0$), rotational motion ($k = 1$), and higher-order rotational and vibrational characteristics ($k > 2$).

The sign of $h_{2k}$ indicates the intrinsic hydropathic composition state ($k = 0$), the intrinsic rotation direction ($k = 1$), and the intrinsic directionality of higher order rotational and vibrational modes ($k > 2$), about $\mathcal{P}$.

For example, the sign of $h_2$ could potentially indicate the intrinsic rotational direction of helical segments relative to the membrane surface, whether clockwise or counterclockwise.

Generally, we consider $|h_{2k}|$ to be the equivalent of the $2k$-th order inertia of an $n$-cluster.

##### S1.5.2 Translational degrees of freedom emerging from water-mediated interactions ($j = 2k + 1$)

$\vec{\boldsymbol{h}}_{2k+1}$ informs about an $n$-cluster's resistance in $(2k + 1)$-dimensional deformations and how these contribute to the generation of dehydration force fields along $\mathcal{P}$. These forces are assumed to control pore conductivity, i.e., they regulate the ion/water flux through the pore.

Focusing on the membrane-perpendicular component, $h_{2k+1,\perp}$, offers a clearer picture about pore conductivity efficiency, since the membrane-parallel component, $h_{2k+1,\parallel}$, has been identified as a noise-sensitive friction force [9, 10].

The sign of $h_{2k+1,\perp}$ indicates the intrinsic conductivity direction along $\mathcal{P}$, i.e., whether the dominant ion/water flux is directed into or out of the cell. Generally, the larger $|h_{2k+1,\perp}|$, the larger the $2k + 1$-th order conductivity of an $n$-cluster would be.

$h_{1,\perp}$ accounts for the linear displacement of permeating species along $\mathcal{P}$. Linearizion of the ion transport implies a catalytic-like effect, where ions are transported through the pore along a nearly linear path (or, equivalently, at a diffusion limited rate [11]), short-circuiting the membrane. Simultaneously, $h_{1,\perp}$ renders an $n$-cluster polarizable thereby increasing its likelihood to interact with its surroundings, which, in turn contributes constructively to its thermostability. For these reasons, $h_{1,\perp}$ is a key index for investigating the potential emergence of critical phenomena in NaVChs.

#### S1.6 Adiabatic processing of perturbations

##### S1.6.1 Geometric structure of the thermostability state space

We introduce the Poincaré half-plane:

$$\mathbb{H} = \{(\mathrm{p}_\perp, \varepsilon) \mid \varepsilon > 0, \mathrm{p}_\perp \in \mathbb{R}\} \tag{S11}$$

endowed with the metric $(\mathrm{d}\omega)^2 = \frac{(\mathrm{dp}_\perp)^2 + (\mathrm{d}\varepsilon)^2}{\varepsilon^2}$.

The distance between the point $(\mathrm{p}_\perp, \varepsilon)$ and the point $(\mathrm{p}_\perp, \varepsilon_i)$ in the half-plane is given by:

$$\left|\ln\left(\frac{\varepsilon}{\varepsilon_i}\right)\right| = \left|\ln\left(\frac{\varepsilon_i}{\varepsilon}\right)\right| = \frac{|\delta l|}{\xi}, \quad \delta l = l - l_i \tag{S12}$$

Let us now suppose that $\varepsilon$ is a perturbed version of $\varepsilon_i$ or vice versa. We note the following points:

1. $\frac{|\delta l|}{\xi}$ can be interpreted as the shortest distance (geodesic) between an initial (unperturbed) and a final (perturbed) thermostability state represented by $(\mathrm{p}_\perp, \varepsilon_i)$ and $(\mathrm{p}_\perp, \varepsilon)$ (or vice versa), respectively.
2. $|\delta l|$ is the length of the shortest path connecting the PD/VSD interface, $\partial\mathcal{B}_i$, with any other interface, $\partial\mathcal{B}$.

The above points become relevant in the perturbation scenario, where constituents are instantaneously displaced from $\partial\mathcal{B}_i$ onto $\partial\mathcal{B}$ or vice versa. Thermodynamically, this is described with the following adiabatic process:

$$\begin{aligned} &\tau := \begin{cases} \frac{T_{\text{pert}}}{T} = \left(\frac{\varepsilon}{\varepsilon_i}\right)^{\gamma-1}, & (\mathrm{p}_\perp, \varepsilon_i) \mapsto (\mathrm{p}_\perp, \varepsilon) \\ \frac{T}{T_{\text{pert}}} = \left(\frac{\varepsilon_i}{\varepsilon}\right)^{\gamma-1}, & (\mathrm{p}_\perp, \varepsilon) \mapsto (\mathrm{p}_\perp, \varepsilon_i) \end{cases}, \quad T_{\text{pert}} := T + \delta T \\ &\text{with} \quad \gamma = 1 + \frac{2}{f} = 2, \quad \delta T = \frac{1-(1-q)\rho}{kZ_\pi}\delta\varepsilon, \quad \delta\varepsilon = \varepsilon - \varepsilon_i, \end{aligned} \tag{S13}$$

where $\tau$ represents the relative temperature change about a pore point, capturing perturbation-induced changes in the hydropathic composition and hydropathic interaction profile of the surrounding PD-forming atomic environment. $\gamma$ is the heat capacity ratio characterizing the adiabatic displacement of atoms from $\partial\mathcal{B}_i$ onto $\partial\mathcal{B}$ or vice versa. $f = 2$ is the number of effective DoF along the radial direction leaving only two atom displacement options: either inwards (i.e., $\delta\varepsilon \geq 0$) or outwards (i.e., $\delta\varepsilon < 0$). $\varepsilon$ works here as the equivalent of an ideal gas density.

Also note that the first definition case of $\tau$ implies atom displacement from $(\mathrm{p}_\perp, \varepsilon_i)$ onto $(\mathrm{p}_\perp, \varepsilon)$, i.e., from $\partial\mathcal{B}_i$ onto $\partial\mathcal{B}$. The second definition case of $\tau$ accounts for the reverse direction.

The following important comments applies to (S13). Obviously NaVChs are not ideal gases. However, operating near the thermodynamic limit described in (S13) is advantageous because it offers adiabatic control over perturbations. Accordingly, internal processing of perturbation involves minimizing heat exchanges between the NaVCh and its environment, while maximizing NaVCh mechanofunctional output via instantaneous stretching or compressing of its helical segments along the radial direction.

##### S1.6.2 A thermostability state geodesic measure for the composite of $n_+ + n_-$ atoms

Let now $\varepsilon_-$ and $\varepsilon_+$ be two arbitrary realizations of $\varepsilon$.

Since $\frac{\varepsilon_+}{\varepsilon_-} \in \mathbb{R}_{>0}$, there is always an $\{\varepsilon_+, \varepsilon_-\}$-pair satisfying the following equation:

$$\chi_j = \frac{\varepsilon_+}{\varepsilon_-} = \left(\frac{T_+}{T_-}\right)^{1/(\gamma-1)} \quad \text{with} \quad -\frac{h_{j,+}}{h_{j,-}} = \frac{\varepsilon_+}{\varepsilon_-}, \tag{S14}$$

where $\chi_j$ is the composite susceptibility (Main Text Eq. (29)). $T_+$ and $T_-$ represent the effective temperatures associated with the group of $n_+$ and $n_-$ atoms residing in $\mathcal{B}$, respectively.

Analogous to (S12), $|\ln(\chi_j)|$ returns the geodesic distance between an initial (unperturbed) state, $(\mathrm{p}_\perp, \varepsilon_-)$, and a final (perturbed) state, $(\mathrm{p}_\perp, \varepsilon_+)$. Accordingly, the smaller $|\ln(\chi_j)|$ is, the closer the composite system of $n_+ + n_-$ atoms is to a state of minimal thermostability.

#### S1.7 Critical exponents

$\tau$ is identified with the reduced NaVCh temperature: $\tau = 1$ signifies the location of the prominent critical inflection point.

For the critical exponents derived below, we follow notation conventions outlined in Ref. [12].

Consistent with the critical inflection point hypothesis, we consider the case where $\partial\mathcal{B}_i$ serves as an attractor: atoms are displaced from $(\mathrm{p}_\perp, \varepsilon)$ onto $(\mathrm{p}_\perp, \varepsilon_i)$, implying that $\tau = \frac{\varepsilon_i}{\varepsilon}$.

##### S1.7.1 Order parameter

According to Main Text Eqs. (11) and (31), we have:

$$\frac{\partial \ln(n/K)}{\partial \ln(l)} = \frac{l}{\xi}\Lambda(\ln(n/K)) \implies \Lambda(\ln(n/K)) = \frac{\xi d_f}{l} = \frac{1}{\left(\left(\frac{\varepsilon}{\varepsilon_i}\right)^{-1} + \nu\right)}, \tag{S15}$$

where $n/K \in (0,1)$ is the order parameter associated with the structural transition from the PD to the VSDs.

For $l = l_i$ and $\nu = 1$, (S15) gives:

$$\beta := \Lambda(\ln(n/K))|_{l=l_i, \nu=1} = 0.5, \tag{S16}$$

where $\beta$ is the mean-field order parameter critical exponent (note that $\nu = 1$ implies a mean-field conditioning for the atomic environment around the current point (for details see Main Text 2.1.1.3). Note also that $l = l_i$ implies $\tau = 1$, indicating that the computation pertains to the reduced temperature at the critical point.

##### S1.7.2 Molecular susceptibility

We re-visit Eqs. (22) and (23) appearing in the Main Text, to define:

$$\begin{aligned} &\mathcal{X} \coloneqq \frac{H_{j,\text{int}}}{H_{j,\text{ext}}} \leq \exp(-l/\xi) =: \mathcal{X}_\text{b} \\ &\text{with} \quad H_{j,\text{int}} \coloneqq |h_j| \; [\text{kcal} \times \text{Å}^j], \quad H_{j,\text{ext}} \coloneqq n(1+\kappa_j\xi/s)\varepsilon_0 l^j \; [\text{kcal} \times \text{Å}^j], \end{aligned} \tag{S17}$$

where $H_{j,\text{int}}$ and $H_{j,\text{ext}}$ are the amplitudes of an internal and external field acting on $\mathcal{P}$, respectively. We refer to $\mathcal{X}$ as the molecular susceptibility. $\mathcal{X}_\text{b}$ introduces an upper bound on $\mathcal{X}$, where the subscript 'b' stands for 'bound'.

We emphasize the difference between $\mathcal{X}$ and the composite susceptibility, $\chi_j$ (Main Text Eq. (29)). $\mathcal{X}$ works only at an envelope level, providing a general rule for the internal field responsiveness. In contrast, $\chi_j$ conveys below-the-envelope information, providing insights on the competition between hydrophobic and hydrophilic dipole field amplitudes.

Focusing on $\mathcal{X}_\text{b}$ and utilizing Main Text Eq. (31), we get:

$$\frac{\partial \ln(\mathcal{X}_\text{b})}{\partial \ln(l)} = \frac{l}{\xi}\Lambda(\ln(\mathcal{X}_\text{b})) \implies \overbrace{\Lambda(\ln(\mathcal{X}_\text{b}))}^{=:-\gamma} = -1 \; \forall l, \tag{S18}$$

where $\gamma$ is the unsigned scaling exponent characterizing $\mathcal{X}_\text{b}$. $\gamma$ serves here as the equivalent of the mean-field susceptibility critical exponent.

##### S1.7.3 Order parameter response to external fields

The response of the order parameter to $j$-th-order external influences is described by the $\delta_j$-exponent according to:

$$\delta_j \coloneqq \frac{\partial \ln(H_{j,\text{ext}})}{\partial \ln(n)} = 1 + \frac{j}{d_f} + \lambda_j, \tag{S19}$$

where $\lambda_j$ is the contribution to $\delta_j$ from the $\frac{\partial \ln(1+\xi\kappa_j/s)}{\partial \ln(n)}$-term.

##### S1.7.4 Scaling exponent relationship

For $l = l_i$ and $\nu = 1$, (S19) leads to the following equation:

$$\begin{aligned} &\delta_j = 1 + 2j\xi/l_i + \lambda_j \implies \\ &\delta_j = 1 + j\xi/(l_i\beta) + \lambda_j \implies \\ &\beta \times (\delta_j - (\lambda_j + 1)) = \gamma_j, \quad \gamma_j = j\xi/l_i, \end{aligned} \tag{S20}$$

constituting a generalization of Widom's scaling law. Note that $\gamma_j$ is a re-scaled version of $\gamma = 1$, with a scaling factor of $j\xi/l_i$.

To put (S20) into a structural context, consider an arbitrary large, outward stretching of a VSD leading to a sharp increase in the radial size, $s$ (Eq. (S5)). Recent experiments support that this phenomenon is a plausible and notable occurrence, underlying a so-called 'dissociation' [13] of a VSD from the PD. In the context of (S20), 'dissociation' implies that $\lambda_j \to 0$, since interfacial adsorption forces have become insignificant with diverging $s$. This leads to:

$$\beta \times (\delta_{j,\lambda_j \to 0} - 1) = \gamma_j, \quad \delta_{j,\lambda_j \to 0} \to 1 + 2j\xi/l_i, \tag{S21}$$

where it becomes obvious that if the condition

$$j\xi = l_i \tag{S22}$$

is met, then the standard form of Widom's scaling law is recovered, as $\gamma_j = 1$ and $\delta_{j,\lambda_j \to 0} \to 3$. Note that generally $\xi < l_i$.

As an example, let us consider the case $j = 1$. When $\xi$ approaches $l_i$ from below, PD constituents are experiencing predominantly attractive interactions, while the PD intrinsic dimension approaches its lower bound given by $1/q$ (Main Text Eq. (19)). This suggests that the standard form of Widom's scaling law naturally arises when spatial organization efficiency is maximized, leading to:

$$\begin{aligned} &\beta \times (\delta_{1,\lambda_1 \to 0} - 1) = \gamma \\ &\text{with} \quad \delta_{1,\lambda_1 \to 0} \to 3, \quad \gamma \overset{(S18)}{=} 1, \quad \beta \overset{(S16)}{=} 0.5 \quad \text{for} \quad l = l_i = \xi, \quad \nu = 1. \end{aligned} \tag{S23}$$

##### S1.7.5 Sandpile slope

The number of effective perturbation modes encoded into a structural location (i.e., the geometric center of a residue undergoing mutation) is bounded from above by the following quantity:

$$\frac{\partial \ln(|\phi_j| \neq 0)}{\partial \ln(\tau)} = \xi \frac{\mathcal{I}_j}{\phi_j}. \tag{S24}$$

Another interpretation of (S24) is that it describes the intensification of the geodesic flow associated with $|\phi_j| \overset{(S14)}{=} |\ln(\chi_j)|$ along the radial direction. Note that $\tau = \frac{\varepsilon}{\varepsilon_i}$ (as also used in (S15) and (S17)).

The larger (S24) is, the larger the number of 'shortcuts' connecting $(\mathrm{p}_\perp, -h_{j,-})$ with $(\mathrm{p}_\perp, h_{j,+})$ becomes, implying that the $n$-cluster thermostability can be easily compromised with increasing cluster size.

We postulate that (S24) quantifies the slope of the sandpile at site $(\mathrm{p}_\perp, \varepsilon)$.

In the Main Text and thereafter, 'perturbation potential' is used as a synonym for 'sandpile slope.'

##### S1.8 Data analytics

Let $\mathcal{O}(\mathbf{p}, l_\alpha)$, $l_\alpha = \bar{R} + \frac{a}{N_\alpha}(L - \bar{R})$, $\alpha = 1, 2, \ldots, N_\alpha \in \mathbb{Z}_{>0}$, be some arbitrary two-dimensional observable array consisting of card($\mathcal{P}$) columns and $N_\alpha$ rows.

###### S1.8.1 Data collapse procedures

###### S1.8.1.1 Trace collapsing

Let us consider a family of NaVCh structures. Each NaVCh structure is represented by a single $\mathcal{O}$. $\mathcal{O}$'s are statistically summarized (i.e., 'collapsed') according to procedures described in Alg. 1.

**Algorithm 1** Data collapse

Step 1. For each NaVCh structure, return the $\alpha$-indexed medians of the $\mathcal{O}$-columns and stack them into the rows of a two-dimensional array (let it be $\tilde{\mathcal{O}}$). Each row of $\tilde{\mathcal{O}}$ is associated with a different NaVCh molecule.
Step 2. Return the $\alpha$-indexed mean values of the $\tilde{\mathcal{O}}$-columns and the corresponding standard deviations. The former and the latter serve as collapsed traces and collapsed trace clouds, respectively.

###### S1.8.1.2 Atom ordering and coloring

Atoms are colored according to the scheme presented in Alg. 2.

**Algorithm 2** Atom coloring

**for** $\mathbf{p} \in \mathcal{P}$ **do**
  **for** $\mathbf{a} \in \mathcal{A}$ **do**
    Calculate $n(\mathbf{p}, \mathrm{r_a})/K$ for $\mathrm{r_a} = \|\overbrace{\mathbf{p} - \mathbf{a}}^{=\vec{\mathbf{r}}_\mathrm{a}}\|$ [Å]
  **end for**
**end for**
For the current atom $\mathbf{a}$, return the mean of $n(\mathbf{p}, \mathrm{r_a})/K$ along $\mathcal{P}$, $\langle n(\mathbf{p}, \mathrm{r_a})/K \rangle_\mathrm{p} \in (0, 1)$.

The quantity $\langle n(\mathbf{p}, \mathrm{r_a})/K \rangle_\mathrm{p}$ represents the PD/VSD ordering score for an **a**-atom, informed from the order parameter, $n/K$ (see also Eq. (S15)). If $\langle n(\mathbf{p}, \mathrm{r_a})/K \rangle_\mathrm{p} \approx 0.5$, then the atom is expected to be found somewhere near the PD/VSD interface.

###### S1.8.2 Inertia and conductivity statistical summary indices

The $\hat{\cdot}$-operator used to compute the inertia and conductivity statistical summary indices (Main Text Fig. 5 caption), returns:

$$\mathrm{med}[\{\underset{l_\mathrm{mut}}{\mathrm{med}}(\mathcal{O}_{2k})\}], \quad \mathcal{O}_{2k} = \mathcal{O}_{2k}(\mathbf{p}, l_\mathrm{mut}) \tag{S25}$$

and

$$\text{med}[\{\underset{l_{\text{mut}}}{\text{med}}\,(\mathcal{O}_{2k+1,\perp})\}], \quad \mathcal{O}_{2k+1,\perp} = \mathcal{O}_{2k+1,\perp}(\mathbf{p}, l_{\text{mut}}) \tag{S26}$$

where $k = 0, 1, \ldots, 5$ determines the moment order (i.e., the dimension of the water-mediated interaction space), and $l_{\text{mut}}$ is the distance between a pore point, $\mathbf{p}$, and a structural location. Note that (S25) and (S26) are $\mathbf{p}$-dependent. Meaning that for every $\mathbf{p}$ we first calculate the median over all $l_{\text{mut}}$-distances for each $k$. Finally, the median of these medians is calculated, represented by (S25) and (S26).

The maximum and minimum values of the clouds surrounding the traces of (S25) and (S26) illustrated in Figs. 5(b),(c),(d),(e) appearing in the Main Text are given by:

$$\max[\{\underset{l_{\text{mut}}}{\text{med}}\,(\mathcal{O}_{*})\}] \tag{S27}$$

and

$$\min[\{\underset{l_{\text{mut}}}{\text{med}}\,(\mathcal{O}_{*})\}] \tag{S28}$$

respectively, where $* = \begin{cases} 2k, & \text{or} \\ 2k+1, \perp & \end{cases}$.

##### S1.8.3 Calculation of $\eta_j$-exponents

Power-law convergence is probed by extracting the slopes and corresponding Pearson coefficients from the $\ln(|h_j|)$-vs.-$\ln(l)$ diagram over the intervals $[l_t, l_i)$ and $(l_i, l_q]$. $l_t$ and $l_q$ mark the $l$-values for which the $n$-curvature (Main Text Eq. (5)) maximizes and minimizes, respectively.

##### S1.8.4 Calculation of $\mathcal{I}_j$

The first-order derivative of the logarithmic composite susceptibility, $\phi_j$, is computed according to procedures presented in Alg. 3.

**Algorithm 3** Calculation of $\mathcal{I}_j$

Step 1. Smooth the $\phi_j$-trace. Smoothing of $\phi_j$ takes place in Python [14] with the `savgol_filter` function for a sliding window of size $\xi$.
Step 2. The smoothed $\phi_j$-trace is numerically differentiated.

#### S1.9 Pain-disease-associated mutations

Tab. S3 provides a summary of all pain-disease-associated mutations analyzed in this study.

The files `SCN9A_data_ClinVar.xlsx` and `SCN9A_data_gnomAD.xlsx`, containing comprehensive variant information from the ClinVar and gnomAD databases, are included in the dataset [15].

The data were obtained from the respective databases on 18 November 2024.

##### S1.10 Machine learning procedures

We work with the `sklearn` library in Python.

We consider a dataset of mutations split into two classes: pathogenic (`class_0`) and non-pathogenic (`class_1`). `class_0` may also include mutation events whose electrophysiological signature significantly differs from that of the wild-type but the phenotype is unknown (see LoF column in Tab. S3).

The core of our algorithm is a k-fold cross-validation procedure, implemented via the `StratifiedKFold` function, described in Alg. 4. Stratification in k-fold cross-validation ensures that each fold maintains the same class distribution as the original dataset. To a certain extent, this can mitigate artifacts introduced by class imbalances.

The classification is implemented by a support vector machine classifier (SVC) using a radial basis function kernel, `SVC(kernel='rbf')`, providing some flexibility with handling non-linear decision boundaries.

**Algorithm 4** K-fold cross-validation

**for** call $= 1, 2, \ldots, N_{\mathrm{calls}}$ **do**
  Shuffle the dataset for some $\mathrm{seed}_{\mathrm{shuffle}}$.
  Split the shuffled dataset into $K_{\mathrm{fold}}$ groups (or folds).
  **for** each unique group **do**
    Take the current group as the test dataset.
    Take the remaining groups as the training dataset.
    Apply the `SVC(kernel='rbf')` model with default parameter values.
  **end for**
**end for**

###### S1.10.1 Local learning: learning at a single pore point

We apply Alg. 4 locally, i.e., at each pore point. The rationale for handling each pore point separately is that each one admits a distinct temperature value (Eq. (S9)).

Structural locations are featured as $\{\partial^d \phi_{2k}/\partial l^d\}$ or $\{\partial^d \phi_{2k+1,\perp}/\partial l^d\}$, $d = 0, 1$, $k = 0, 1, \ldots, 5$, depending on whether we probe the violation of inertia or conductivity constraints, respectively.

Structural locations are locally characterized by a median statistic, let it be $\mathscr{L}$, summarizing some arbitrary set of values of cardinality $(K_{\mathrm{fold}} - 1)N_{\mathrm{call}}$ returned by Alg. 4.

For example, $\mathscr{L}$ could represent the median of the local `class_0` probabilities, i.e., the probability that the mutability of a structural location is linked to a disease phenotype.

Specifically, denoting the local `class_0` probability set as $\mathrm{prob}^{\mathtt{class\_0}}_{\mathbf{p},d,\mathrm{DoF}}$, we have:

$$\mathscr{L}^{\mathtt{class\_0}}_{\mathbf{p},d,\mathrm{DoF}} \leftarrow \operatorname*{med}_i(\mathrm{prob}^{\mathtt{class\_0}}_{\mathbf{p},d,\mathrm{DoF}}), \quad i = 1, 2, \ldots, (K_{\mathrm{fold}} - 1)N_{\mathrm{call}}, \tag{S29}$$

where the subscripts '$\mathbf{p}, d$, DoF' indicate that $\mathscr{L}^{\texttt{class\_0}}$ depends on $\mathbf{p}$, $d$, and DoF. DoF admits two values, namely, either 'iner.' or 'cond.', depending on whether the feature input is $\{\partial^d \phi_{2k}/\partial l^d\}$ or $\{\partial^d \phi_{2k+1,\perp}/\partial l^d\}$, respectively.

Procedures are described in Alg. 5.

**Algorithm 5** Local Learning

```
for p ∈ P do
    for d = 0, 1 do
        for DoF = {iner., cond.} do
            if (DoF = iner.) then
                for call = 1, 2, ..., N_call do
                    Execute Alg. 4 with seed_shuffle. Structural locations are featured as {∂^d φ_2k / ∂l^d}_{k=0,1,...,5}
                end for
            else if (DoF = cond.) then
                for call = 1, 2, ..., N_call do
                    Execute Alg. (4) with seed_shuffle. Structural locations are featured as {∂^d φ_2k+1,⊥ / ∂l^d}_{k=0,1,...,5}
                end for
            end if
            Return L^class_0_{p,d,DoF} characterizing each structural location locally
        end for
    end for
end for
```

##### S1.10.2 Global learning: learning from all pore points

We apply Alg. 4 globally, i.e., along $\mathcal{P}$.

Structural locations are featured as:

$$\{\operatorname*{med}_{\mathbf{p}}(\mathscr{L}^{\texttt{class\_0}}_{\mathbf{p},d,\mathrm{DoF}})\} \tag{S30}$$

representing the set of global `class_0` probability values. Note that for $d=0,1$, each structural location admits $2 \times (\max(d)+1) = 4$ feature input values, i.e.,:

$$\{\operatorname*{med}_{\mathbf{p}}(\mathscr{L}^{\texttt{class\_0}}_{\mathbf{p},0,\mathrm{iner.}}), \operatorname*{med}_{\mathbf{p}}(\mathscr{L}^{\texttt{class\_0}}_{\mathbf{p},1,\mathrm{iner.}}), \operatorname*{med}_{\mathbf{p}}(\mathscr{L}^{\texttt{class\_0}}_{\mathbf{p},0,\mathrm{cond.}}), \operatorname*{med}_{\mathbf{p}}(\mathscr{L}^{\texttt{class\_0}}_{\mathbf{p},1,\mathrm{cond.}}),\}$$

.

Procedures are described in Alg. 6.

##### S1.10.3 Machine learning parameters

In Tab. S4 we list machine learning parameters. Note that we did not consider any parameter optimization.

**Algorithm 6** Global learning

**for** call$=1,2,\dots,N_{\text{call}}$ **do**

Execute Alg. 4 with seed$_{\text{shuffle}}$. Structural locations are featured as $\{\underset{\mathbf{p}}{\text{med}}(\mathscr{L}^{\texttt{class\_0}}_{\mathbf{p},d,\text{DoF}})\}$

**end for**

##### S1.11 *N*-curve fitting procedures

We follow [9].

To avoid overfitting, at each pore point, we generate three candidate models:

- General case: $A(1+\nu\exp(-\frac{\nu}{\zeta}(l-l_i)))^{-1/\nu}$ (fitted parameters: $\{A,\zeta,l_i,\nu\}$)
- Logistic case ($\nu\!=\!1$): $A(1+\nu\exp(-\frac{1}{\zeta}(l-l_i)))^{-1}$ (fitted parameters: $\{A,\zeta=\xi,l_i\}$)
- Gompertz case ($\nu\to 0$): $A\exp(-\exp(-(l-l_{i,\zeta\to 0})/\xi))$ (fitted parameters: $\{A,\xi,l_{i,\zeta\to 0}\}$)

and fit them individually to the normalized experimental trace, $N/N_{\text{total}}$ (Main Text Expression (32)). Note that once $A$ is known, the NaVCh structure carrying capacity, $K$, is given by $AN_{\text{total}}$.

The best-fitting candidate model is selected by minimizing an Akaike information criterion. The `L-BGS-B` algorithm in Python [14] was used to fit the candidate model traces.

### S2 Supplementary Results

#### S2.1 Atom packing model quality assessment

We use a two-sample Cramér-von Mises criterion to assess the goodness of the fit of the theoretical cumulative atom number, $n$, to the normalized experimental trace $N/N_{\text{total}}$.

The corresponding P-value is computed in Python using the built-in `cramervonmises` function with argument `method = 'asymptotic'`. We also calculate the mean absolute fitting error (MAFE) characterizing the goodness of the fit of $n$ to $N/N_{\text{total}}$.

The histograms of the P-values and MAFEs are summarized in Figs. S2(a) and (b) for prokaryotes and eukaryotes, respectively. The closer the P-value histogram is to unit, and the closer the MAFE histogram is to zero, the higher the model quality. Specifically, if the P-value exceeds a pre-determined significance level, the null hypothesis – that the observed samples are drawn from the same distribution – cannot be rejected.

The smallness of the MAFEs strongly supports our fitting hypothesis. On the other hand, the P-values are primarily distributed near zero, indicating that the underlying distribution may actually differ. This is expected since the focus is on the coarse-grained form of $N$.

These observations support the dominance but not the uniqueness of the detected inflection point. In fact, there are multiple, less prominent – from a thermostability viewpoint – inflection points distributed along the entire $l$-range, reflecting slowly oscillating modes of $N/N_{\mathrm{total}}$, which may become important under certain metastable conditioning. Specific examples are given in Section S2.4.

##### S2.2 Pore domain resistance to excess entropy

In Figs. S3(a),(b), we demonstrate that, for both prokaryotic and eukaryotic NaVChs, the statistical mechanical entropy, $\mathcal{S}$, approaches its maximum, $\mathcal{S}_{q\to 1}$ (Main Text Eq. (18)), somewhere near the center of the pore, i.e., for $\mathrm{p}_\perp \approx 0$, where the surplus of hydrophobic constituents drives the formation of the CC (e.g., see Fig. 1 in [16]). Simultaneously, $l_i$ is minimized (Figs. S3(a),(b)).

Main Text Eq. (20) prescribes how these events promote a lowering of the intrinsic dimension of the PD (Fig. S1). Structural disorder due to excess DoF is thus mitigated ensuring that the tight packing of atoms around the CC pore region is not compromised.

##### S2.3 Hydropathic dipole field decomposability

We verified that the hydrophobic and hydrophilic components of $h_{j=2k}$ and $h_{j=2k+1,\perp}$, extracted from the high-resolution NaV1.7 channel (PDB code: 7w9k), satisfy the Decomposition Ansatz (Main Text Eq. (24)) for $k \geq 0$ and $k \geq 1$, respectively, for an acceptably small cutoff scale, $l_{\mathrm{cutoff}} \approx l_0$.

As discussed below, although the decomposability of the first-order hydropathic dipole field (HDF), $h_{1,\perp}$, is achievable only for a portion of $\mathcal{P}$, this does not hinder further analysis.

###### S2.3.1 First-order hydropathic dipole field decomposability

Since the first-order hydrophilic, $h_{1,\perp,\mathrm{philic}}$ [kcal×Å], and hydrophobic, $h_{1,\perp,\mathrm{phobic}}$ [kcal×Å], membrane-vertical field components satisfy the Decomposition Ansatz described in Main Text Eq. (24) only partially (i.e., for $\mathrm{p}_\perp < -4.19$ and $\mathrm{p}_\perp > 18.2$ (see Figs. S4(c),(d)), we re-define the composite susceptibility as $\chi_{1,\perp} := \left|\frac{h_{1,\perp,+}}{h_{1,\perp,-}\neq 0}\right|$, where $h_{1,\perp,-} = h_{1,\perp,\mathrm{phobic}}$ and $h_{1,\perp,-} = h_{1,\perp,\mathrm{philic}}$. Evidently, by doing so, we are facing the risk that $\phi_{1,\perp} = \ln(\chi_{1,\perp})$ may blow-up as $h_{1,\perp,-}$ crosses zero. Namely, a continuous zero-crossing transition would result in a unmanageably large value in the $\phi_{1,\perp}$ spectrum, given that the sampling along the radial direction is sufficiently dense. As shown in Fig. S4(b), this is not the case as the distribution of the collapsed $\phi_{1,\perp}$-values remains confined within $[-10.7, 9.1]$.

This finding suggests that changes in the orientation of $h_{1,\perp,-}\hat{\mathbf{e}}_\perp$ are abrupt, discontinuous events generated by $h_{1,\perp,-}$ 'jumping over' (rather than 'crossing smoothly') zero. Singularities are thus mitigated, as their peaks are contained, preventing composite susceptibility infinities, even at doubled sampling rate (Fig. S4 caption). We thus confidently proceed with further statistical analysis of $\phi_{1,\perp}$.

An exemplary trace of $\phi_{1,\perp}$, sampled from the pore region between the selectivity filter (SF) and the CC, is shown in Fig. S4(a). We notice the formation of sharp but nevertheless non-divergent corners (Fig. S4(a)).

In the inset of Fig. S4(a), we illustrate the zero-crossing behavior of $h_{1,\perp,+}$ and $h_{1,\perp,-}$, where it becomes evident that $h_{1,\perp,+}$ and $h_{1,\perp,-}$ are related via an exchange-sign operation, namely, $h_{1,\perp,+} \mapsto -h_{1,\perp,-}$ roughly holds.

Interestingly, $h_{1,\perp,-}$ crosses zero at $l \approx l_i$ revealing that, while transitioning from the SF pore region towards the CC pore region, the HDF induced by the PD on $\mathcal{P}$ is purely driven by hydrophilic constituents.

According to S1.8.2, we obtained a statistical summary of $\phi_{1,\perp}$ across NaV1.7 structural locations where pain-disease-associated mutations and non-pathogenic mutations appear.

At the SF pore region, we notice that the GoF- and LoF-derived $\phi_{1,\perp}$-medians remain relatively close to zero (Fig. S4(c)). On the other hand, the neutral- and benign-associated $\phi_{1,\perp}$-medians are negatively minimized at the extracellular side (ES) of the SF pore region (Fig. S4(c)). Negative minimization of the GoF- and LoF-derived $\phi_{1,\perp}$-medians occurs much later, namely, at the pore region corresponding to the extracellular funnel.

This observation suggests that $\phi_{1,\perp}$-traces associated with structural locations where GoF and LoF mutations appear, are more likely to exhibit double zero-crossings at the SF pore region, compared to those associated with neutral and benign mutations. Pain-disease-associated mutations could thus cause indecisiveness in conductivity directionality and amplitude exactly at the SF pore region, which could potentially affect the ion coordination and selection process.

The GoF- and LoF-derived $\mathcal{I}_{1,\perp}$-medians exhibit oppositely correlated fluctuations, with amplitudes significantly greater than those associated with neutral and benign structural locations, particularly after the CC pore region towards the ES (Figs. S4(d)). This suggests that GoF and LoF mutations appear in structural locations that, when perturbed, are likely to trigger oppositely directed mechanofunctional responses, aligning well with the statistical tendencies illustrated in Main Text Fig. 5(e).

#### S2.4 Spatial organization features of prokaryotic NaVCh molecules

##### S2.4.1 Inflection point geometry, hydropathic energy, and entropy

A prominent inflection point characterizes the spatial organization of prokaryotic NaVCh structures, except in the case of the NaVAe structures describing a PD with a C-terminal domain (CTD) extension (see Figs. S5-S6 and S8-S10(a) versus S7(a)).

The thermostability of the PD/CTD assembly is determined by a pair of prominent inflection points. Our algorithm was still able to fit the experimental $N$-traces extracted from NaVAe structures, however, with low fidelity (Fig. (S7(a)). As we can see in Fig. (S7(b)), the second inflection point marks the transition from the PD towards the CTD mediated by a neck region extending towards a coiled coil (Fig. S7(b)). Experimental evidence suggests that the neck acts as a flexible metastable linker allowing the CTD to exert significant thermodynamic control over the NaVAe gating properties [17]. This insight, combined with findings shown here, suggests that, in principle, multiple inflection points may coexist and collaboratively influence the gating cycle of a NaVCh. NaVCh gating dynamics is then enriched since distributing thermodynamic control throughout several inflection points increases the likelihood of asynchronous and/or oppositely directed mechanofunctional responses accessed for different initial conditions and/or via different perturbation modes.

We propose that optimal thermostability necessitates the dominance of a single inflection point, accompanied by several peripheral inflection points whose thermostability significance hierarchically diminishes. On these grounds, we would expect that inclusion of the VSDs in the NaVAe PD/CTD assembly would allow to refocus on a single, prominent inflection point. Our findings corroborate this idea (see Fig. S8(b)). We also report that a prominent inflection point underlies the thermostability of autonomously folded S5-6 transmembrane helices (Fig. S9(b)), suggesting that prokaryotic PDs can stably exist without the VSDs [18].

$\varepsilon$ offers a good qualitative description of the decaying unsigned atomic hydropathic energy, $|h_0|/N$ [kcal/atom], for increasing $l$ (Figs. S5-S10(c)). This indicates that finite-size effects, modeled with the interfacial adsorption bond length, $\kappa_j \xi$, can be neglected up to a certain level of approximation.

Nonextensive entropic constraining largely persist along prokaryotic pores (Figs. S5-S10(d),(f)), corroborating our understanding of the key role that $q$ plays in shaping $\mathcal{S}$.

##### S2.4.2 Dipole field topological fingerprint of prokaryotic pore domains

In NaChBac and NaVAb channels, we observe that the $h_{1,\perp,\downarrow}$ and $h_{1,\perp,\uparrow}$ HDF-instances induced at the IS and ES of the pore are preferably oriented intracellularly and extracellularly, respectively (Figs. S5,S6(e)).

Generally, this favors the formation of centrally located repeller-like HDF topologies. This expectation is verified for a prototype NaVAb channel captured at a pre-open state (see [16] Fig. 2).

In the NaVMs pore-only structures, the opposite correlation is observed, namely, HDF-instances induced at the ES and IS of the pore are oriented extracellularly and intracellularly, respectively, promoting an attractor-like hydrodynamic profile (Fig. S9(e)).

For the remaining structures, HDF-traces exhibit large-amplitude fluctuations, making it difficult to discern clear trends (Figs. S7,S8,S10(e)), implying a mixture of attractor-vs.-repeller hydrodynamic patters.

##### S2.4.3 Diversification of dipole field scaling behavior beyond prokaryotic pore domains

In Fig. S11(a), we show that prokaryotic $\eta_{1,\perp,<}$-exponents do not exhibit any notable distributional preferences for the ES or the IS of the pore, suggesting that $h_{1,\perp,\downarrow}$ and $h_{1,\perp,\uparrow}$ no longer exhibit a preferred orientation state. This implies that the amplification direction of allosteric signals coupling the PD with the VSDs does not correlate with the ES or the IS in any obvious manner. Perturbations can thus be either amplified or attenuated as they propagate between the PD and the VSDs on both sides of the pore.

#### S2.5 Spatial organization features of eukaryotic NaVChs

##### S2.5.1 Inflection point geometry, hydropathic energy, and entropy

We report an excellent agreement between $N$ and $n$, indicating that the dominance of the PD/VSD inflection point is preserved in all cases (Figs. S12-S22(a)). The addition of the $\beta$ subunits to the PD/VSD assembly does not compromise the quality of the fitting (Figs. S13-S19(b) and S19-S20(b)).

The correlation between the profile of the unsigned atomic hydropathic energy and $\varepsilon$ remains qualitative only (Figs. S12-S22(c)). Interfacial adsorption effects appear to be more pronounced beyond the PD in eukaryotes compared to prokaryotes, as evidenced by the increase in the deviation between $\varepsilon$ and $|h_0|/N$ for $l > l_i$ (see Fig. S20(c) versus S5,S6(c)). Determining whether this reflects VSD low-resolution artifacts or a genuine physical NaVCh property requires further investigation.

Entropic constrains apply stronger in eukaryotes compared to prokaryotes, as the dispersion of $\mathcal{S}$-values in the $(\mathcal{S}, \xi, \zeta)$ space is less pronounced (see Figs. S12-S22(d) versus S5-S10(d)). The dependence of $\mathcal{S}$ on water-mediated interactions is thus enhanced, while simultaneously $\mathcal{S}$ is further decoupled from size constraints, $s$ (see Figs. S12-S22(f) versus S5-S10(f)).

##### S2.5.2 Dipole field topological fingerprint of eukaryotic pore domains

In constrast to prokaryotic structures, we observe that $h_{1,\perp,\downarrow}$ and $h_{1,\perp,\uparrow}$ HDF-instances induced at the IS and ES of the pore tend to be oriented extracellularly and intracellularly, respectively (Figs. S12-S22(e)). This favors the formation of attractor-like hydrodynamic topologies, where ions accumulate in the CC so that their passage through the channel is prevented.

##### S2.5.3 Diversification of dipole field scaling behavior beyond eukaryotic pore domains

For $l > l_i$, $h_{1,\perp,\downarrow}$ and $h_{1,\perp,\uparrow}$ HDF-instances induced at the IS and ES of the pore tend to vanish and increase, respectively (Figs. S12-S22(e)).

This observation is independent of the structural inclusion of the $\beta$ subunits, implying that this pattern is an intrinsic characteristic of the PD/VSD structural transition (compare structures with and without $\beta$ subunits and notice that observed HDF transients remain invariant: Figs. S13-S19,S19-S20(e) vs. S18,S21,S12(e)).

In Fig. S11(b), we demonstrate that, in contrast to their prokaryotic counterparts, eukaryotic $\eta_{1,\perp,>}$-exponents show distinct distributional preferences for either the ES or the IS. A clear preference of positive and negative eukaryotic $\eta_{1,\perp,>}$-exponents for the IS and the ES, respectively, is observed. This finding implies that the amplification direction of allosteric signals, coupling the PD with the VSDs, correlates with the

IS but not with the ES. Perturbations propagating from the PD towards the VSDs are thus expected to be attenuated if they are initiated at the ES, while they are expected to be amplified if they are initiated at the IS. Thus, broken HDF symmetries in eukaryotic NaVChs result in a more specialized (i.e., pore-side-specific) nature for PD/VSD coupling in eukaryotes.

An illustrative example is provided in Fig. S23. There, we show that, for the high-resolution 7w9k NaV1.7 molecule, the $\eta_{1,\perp,>}$-exponent spectrum negatively minimize at the ES of the SF pore region, while positively maximizing near the AG pore region, exhibiting an odd-symmetry profile (notice how the $\eta_{1,\perp,>}$-exponent spectrum is organized with respect to the pseudo(i.e., non-exact)symmetry axis roughly dichotomizing the pore). In contrast, $\eta_{1,\perp,<}$-exponents, describing the scaling behavior of $|h_{1,\perp}|$ [kcal×Å] in the PD, remain close to their mean value of approximately 3.5 (refer to Main Text Fig. 3 for further details), while adhering to an even-symmetry pattern (notice how the $\eta_{1,\perp,<}$-exponent spectrum is organized in relation to the pseudosymmetry axis shown in Fig. S23).

We also notice that the positive maxima and negative minima of the $\eta_{1,\perp,>}$-exponent spectrum coincide with the pore regions where GoF and LoF mutation subsets converge, i.e., tend to minimize their respective Euclidean distances from. Simultaneously, the Pearson coefficients associated with both $\eta_{1,\perp,<}$ and $\eta_{1,\perp,>}$ reach their maximum values (Fig. S23). This observation suggests that the structural locations of GoF and LoF mutations are embedded in distinct allosteric pathways, with GoF mutations linked to the amplification of perturbations in the VSDs, while LoF mutations associated with their attenuation. A complementary observation is that the interaction range ratio, $\nu$ approaches zero and unity, respectively, indicating how the atomic environment transitions from a tightly packed configuration at the ES to a nearly mean-field state at the IS.

#### S2.6 Machine learning experiments

##### S2.6.1 Stable local learning

As deduced from the plots appearing in the left panel of Figs. S24 and S25, the feature inputs $\phi_{j=0,1,\ldots,11}$ result in generally stable machine learning algorithm performance.

We observe a weak but consistent trend of machine learning algorithm performance improvement towards the center of the pore alongside with a noticeable decrease in performance at the ES. The reason for this behavior could be partially attributed to finite-size effects: $\partial n$-shell occupancy decreases as we move towards the IS or the ES, affecting the quality of $\phi_j$ and, consequently, also the learnability of the algorithm.

For the dataset incorporating the classes of pain-disease-associated mutations (GoF∪LoF) and of neutral and benign (Neutr.∪Benign), the algorithm shows a recovery in performance, achieving a stable predictive accuracy also at the ES (Figs. S24(c), S25(c)). This is noteworthy because one might expect the significant imbalance between GoF∪LoF and Neutr.∪Benign classes to negatively affect local learnability of the algorithm. The lack of such an effect is an important finding, supporting the distinct perturbation potential profile for the structural locations where pain-disease-associated mutations appear.

For the feature inputs $\mathcal{I}_{j=0,1,\ldots,11}$ we observe significantly stronger fluctuations in the machine learning algorithm's performance (right panel of Figs. S24 and S25).

This instability arises from the computation method of $\mathcal{I}_j$, which struggles to accurately represent the underlying biophysical phenomenon of interest, namely, the strength and directionality of the interfacial coupling. This is due to the smoothing procedure that $\phi_j$ undergoes before being differentiated to compute $\mathcal{I}_j$ (see Alg. 3), which, while intended to reduce noise and make $\phi_j$ more tractable for differentiation, can filter-out important variations of the interfacial coupling, $\Delta\psi_j$ (Main Text Eq. (27)). A crucial parameter that could be thus fine-tuned in the future is the size of the smoothing window, currently conventionally set to be equal with the attractive interaction range, $\xi$.

##### S2.6.2 Importance of interfacial inertia and conductivity features

In Fig. S26, we display the feature importance of interfacial inertia inputs $\mathcal{I}_{j=0,1,\ldots,11}$. Notably, the feature importance of $\mathcal{I}_0$ is elevated at the CC pore region (Fig. S26(a)). CC's stability is not relying on the precise geometrical arrangement of constituents. Hence, residue substitutions that do not significantly alter the hydrophobic profile of the CC are expected to be tolerated, regardless of any interaction geometry changes they may induce.

At the SF pore region, the significance of $\mathcal{I}_0$ diminishes, along with $\mathcal{I}_2$, while higher-order terms (strikingly, up to tenth-order) dominate (Fig. S26(a)). This suggests that $\partial n$-shell higher-order rotational and vibrational modes are important for the ion selection process. It is thus unlikely that substituting one residue for another would have a neutral effect or even be tolerated, as even small changes in the interaction geometry could have an impact.

$\mathcal{I}_{\perp,1}$ emerges as the most significant interfacial conductivity feature, as shown in Fig. S26(b). However, a pronounced rise in $\mathcal{I}_{\perp,3}$ alongside with a simultaneous decline in $\mathcal{I}_{\perp,1}$ is observed precisely at the extracellular entrance of the SF, as illustrated in Fig. S26(b). This suggests that volumetric constraints introduced by the $\partial n$-shells surrounding the SF could be important for managing the initial dehydration and proper positioning of ions before they move deeper into the narrow SF passage.

##### S2.6.3 Machine learning experimentation with abolished directionality

In Tab. S5 we summarize the findings concerning the machine learning experiments with the feature inputs $\{|\phi_{2k}|, |\mathcal{I}_{2k}|, |\phi_{2k+1,\perp}|, |\mathcal{I}_{2k+1,\perp}|\}$, $k = 0, 1, \ldots, 5$, probing the violation of inertia and conductivity constrains irrespective of the perturbation response directionality.

The values appearing in Tab. S5 are largely similar to those appearing in the Tab. 2 in the Main Text.

The differences in the algorithm's performance for switching between the feature inputs $\{|\phi_{2k}|, |\mathcal{I}_{2k}|\}$ and $\{|\phi_{2k+1,\perp}|, |\mathcal{I}_{2k+1,\perp}|\}$, $k = 0, 1, \ldots, 5$, is only minor, supporting that both feature inputs carry equally weighted, biophysical information.

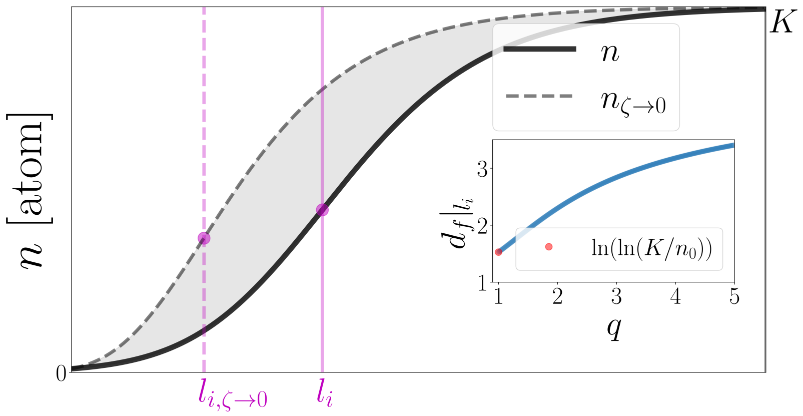


**Fig. S1**: **Tightening of the atomic environment around a pore point**. The cumulative atom number limit case, $n_{\zeta\to 0}$ [atom], is compared to a reference $n$-trace for fixed parameter values $l_0, n_0, \xi, K$. This comparison highlights the effect of a diminished repulsive interaction range ($\zeta \to 0$) on the $n$-trace. In the inset, we illustrate how the intrinsic dimension of the pore domain, $d_f|l_i$, increases monotonically with the so-called nonextensivity index [7], $q = 1 + \nu = 1 + \frac{\zeta}{\xi}$, driven by the increase in $\zeta$. The x-axis locations of the inflection points $l_{i,\zeta\to 0} = \lim_{\zeta\to 0} l_i = \xi \ln(\ln(K/n_0)) + l_0$ [Å] and $l_i = \xi \ln\left(\frac{(n_0/K)^{-\nu} - 1}{\nu}\right) + l_0$ [Å] are also highlighted.

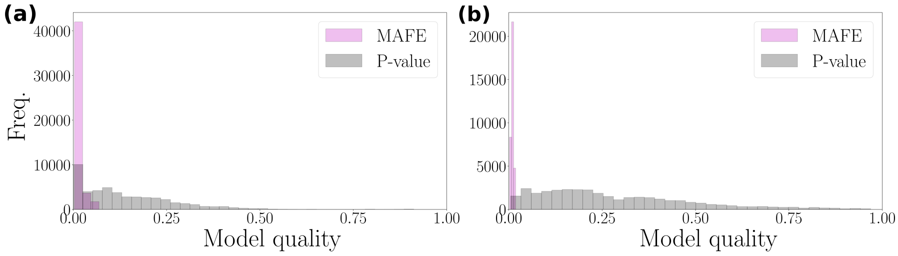


**Fig. S2**: **Assessing the quality of the generalized logistic atom packing model**. P-values and mean absolute fitting errors (MAFEs) are obtained by fitting the theoretical cumulative atom number, $n$ [atom], on the empirical one given by $N/N_{\mathrm{total}}$ (Main Text Eq. (3) and Expression (32), respectively). P-value and MAFE histograms are illustrated for 71 atomic structures of prokaryotic origin and 50 of eukaryotic origin in **(a)** and **(b)**, respectively (see Tab. S1).

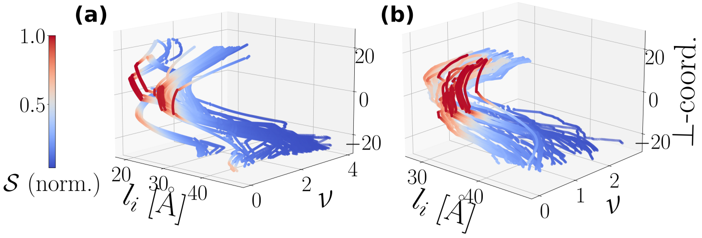


**Fig. S3**: **The geometric shape of the PD**. We illustrate the dependence of the characteristic size of the PD, $l_i$ [Å], on the interaction range ratio, $\nu$, and the statistical mechanical entropy, $\mathcal{S}$ [kcal/($\Theta\times$atom)], for $\mathbf{p}\in\mathcal{P}$ (represented in terms of $p_\perp$). In **(a)** and **(b)**, we overplot empirical traces derived from 71 prokaryotic and 50 eukaryotic NaVCh structures, respectively (Tab. S1). Note that $\mathcal{S}$ is normalized over its maximum value, $\mathcal{S}_{\max}$, given by Main Text Eq. (18).

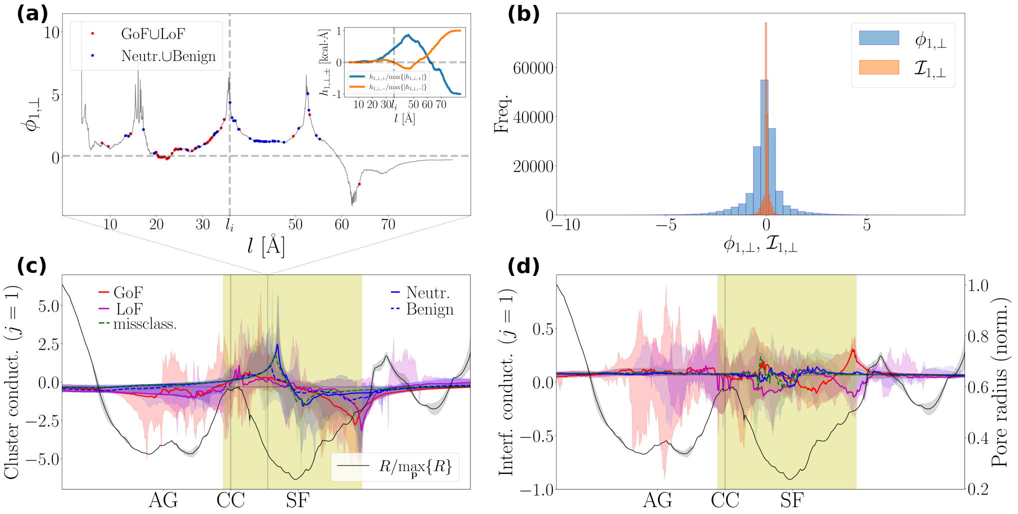


**Fig. S4**: **First-order conductivity profile of structural locations attracting pain-disease-associated mutations**. In **(a)** we show the trace of the membrane-perpendicular first-oder logarithmic composite susceptibility, $\phi_{1,\perp}$, sampled at $\mathrm{p}_\perp = 3.0$. An exceptionally large sampling rate along the radial direction is considered by $\alpha = 1, 2, \ldots, N_\alpha = 1600$. In the inset, we plot the traces of the corresponding hydrophilic and hydrophobic components, denoted as $h_{1,\perp,+}$ and $h_{1,\perp,-}$, respectively. **(b)**, the distribution of the collapsed $\phi_{1,\perp}$ and $\mathcal{I}_{1,\perp}$ values (i.e., of all $\phi_{1,\perp}$ and $\mathcal{I}_{1,\perp}$ values obtained along $\mathcal{P}$ for $\alpha = 1, 2, \ldots, N_\alpha = 1600$). **(c)**, we plot the first-order conductivity cluster index, $\underset{l_{\mathrm{mut}}}{\mathrm{med}}(\phi_{1,\perp}(\mathbf{p} \in \mathcal{P}, l_{\mathrm{mut}}))$ (see (S26)), where $l_{\mathrm{mut}}$ is the Euclidean distance between $\mathbf{p}$ and a structural location where a mutation appears. We consider the subsets of gain-of-function (GoF), loss-of-function (LoF), neutral (Neutr.), benign mutations. Missclassified (missclass.) mutation subset concerns GoF or LoF structural locations which are consistently misclassified by our machine learning procedures (Main Text Fig. 6(b)). **(d)** is equivalent to **(c)**, but instead, it displays the traces of the first-order conductivity interfacial index, $\underset{l_{\mathrm{mut}}}{\mathrm{med}}(\mathcal{I}_{1,\perp}(\mathbf{p}, l_{\mathrm{mut}}))$. Shaded areas highlight the 5%-95% percentile interval around the medians. The light yellow shaded area illustrates the pore region where the Decomposition Ansatz is violated, i.e., $-4.19 \leq \mathrm{p}_\perp \leq 18.2$. Data appearing in **(c)** and **(d)** were created by using the standard resolution used throughout this work, namely, $\alpha = 0, 1, \ldots, N_\alpha = 800$, to maintain consistency in the computation of conductivity statistical summary indices (S26) throughout.

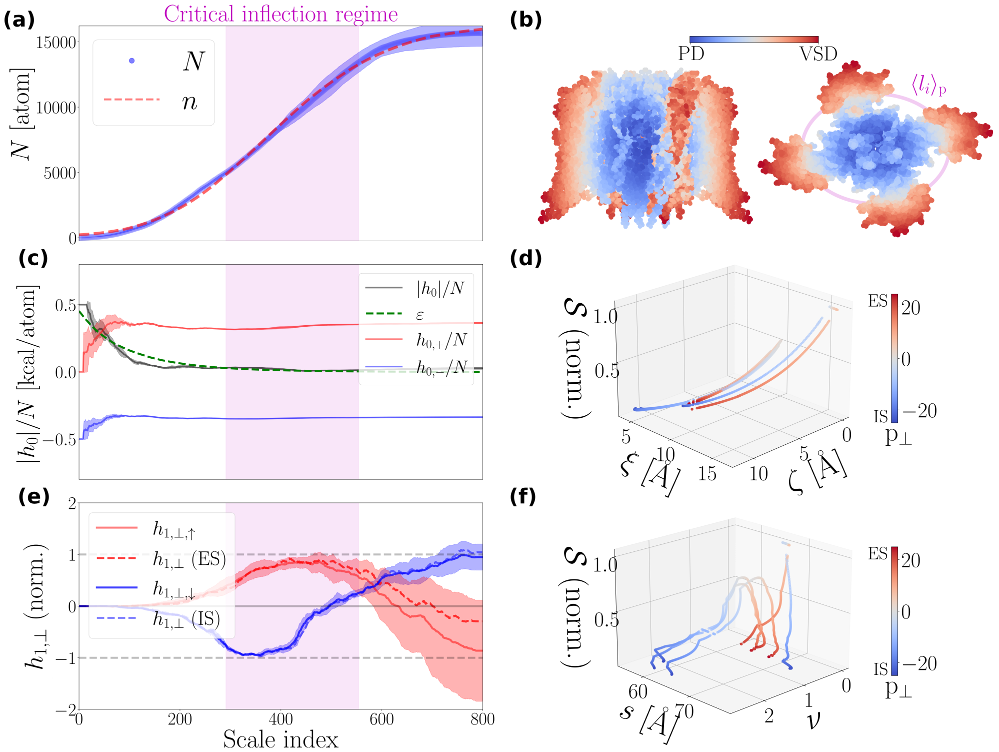


**Fig. S5**: **Spatial organization observables of 4 NaChBac molecules (see Tab. S1)**. Relevant observables were extracted for $\mathbf{p} \in \mathcal{P}$ and scale indices $\alpha = 1, 2, \ldots, N_\alpha = 800$ (Main Text Eq. (12)). **(a)** Empirical, $N$ [atom], and best-fitted theoretical, $n$ [atom], collapsed traces of the cumulative atom number. **(b)**, Illustration of a NaChBac molecule (PDB code: 6vwx). Atoms are colored according to their ordering score (see Alg. 2) and mapped on a membrane-vertical (left) and membrane-parallel (right) plane. $\langle l_i \rangle_\mathrm{p}$ [Å] is the mean value (computed over $\mathcal{P}$) of the characteristic PD size. **(c)**, Empirical, $\frac{|h_0|}{n}$ [kcal/atom], and theoretical, $\varepsilon$ [kcal/atom], collapsed traces of the unsigned atomic hydropathic energies with $h_{0,+} + h_{0,-} = h_0$ (Main Text Eqs. (21),(24)). **(d)**, The interplay between the statistical mechanical entropy, $\mathcal{S}$, and the interaction range pair, $\{\xi, \zeta\}$. **(e)**, Collapsed traces of 'pointing-ES' and 'pointing-IS' instances of $h_{1,\perp}$ [kcal×Å/atom] (Main Text Eq. (21)). $h_{1,\perp}$ is normalized over $|h_{1,\perp}|_{l_i}|$. If $h_{1,\perp}|_{l_i} > 0$, $h_{1,\perp}$ is labeled as a 'pointing-ES' (↑) instance; otherwise, 'pointing-IS' (↓). Dashed lines represent collapsed traces of instances of $h_{1,\perp}$ computer over $\mathrm{p}_\perp < 0$ and $\mathrm{p}_\perp > 0$, associated with the IS and the ES of pore, respectively. **(f)**, The nonextensive profile of $\mathcal{S}$ [kcal/(Θ×atom)], as revealed from its dependence over the molecular radial size, $s$ [Å], and the interaction range ratio, $\nu = \frac{\xi}{\zeta}$. The critical inflection regime highlights the interval from which all $l_i$-values are drawn. Collapsed traces are computed according to procedures described in S1.8.1. $\mathcal{S}$ is normalized over $\mathcal{S}_\mathrm{max}$ (Main Text Eq. (18)).

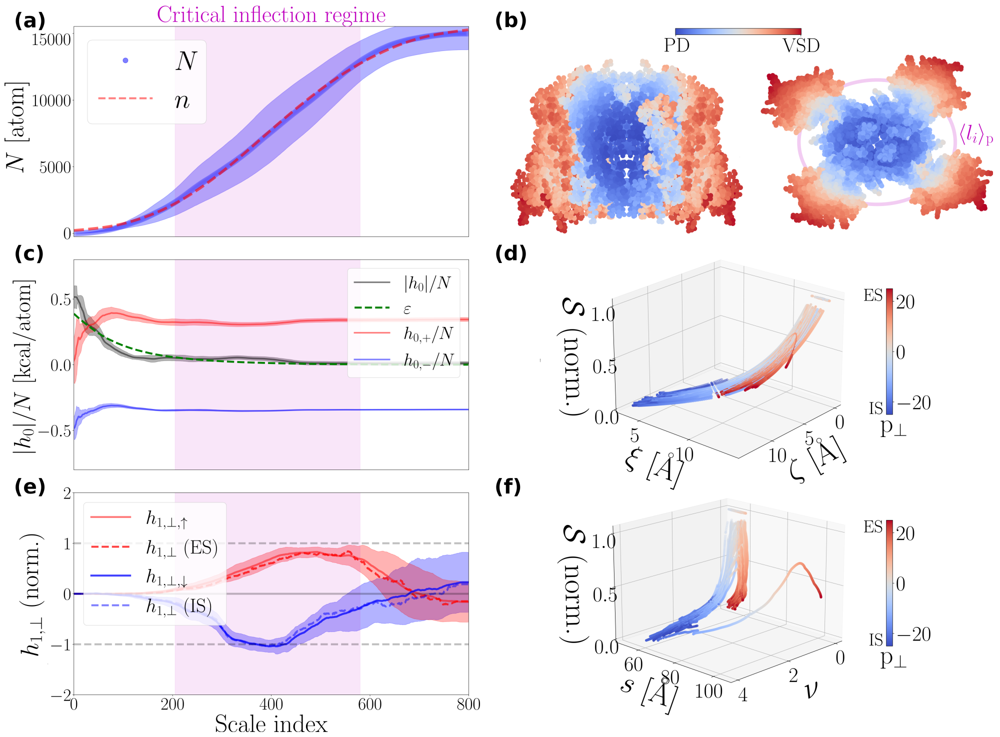


**Fig. S6**: **Spatial organization observables of 33 NaVAb molecules (see Tab. S1).** **(a)**, **(b)**, **(c)**, **(d)**, **(e)**, **(f)** illustrate same type observables as in Fig. (S5), computed using the same data analytics procedure (see S1.8.1), for $\mathbf{p} \in \mathcal{P}$ and $\alpha = 1, 2, \ldots, N_\alpha = 800$. In **(b)**, we illustrate a NaVAb molecule captured in a potentially inactivated state (PDB code: 4ekw).

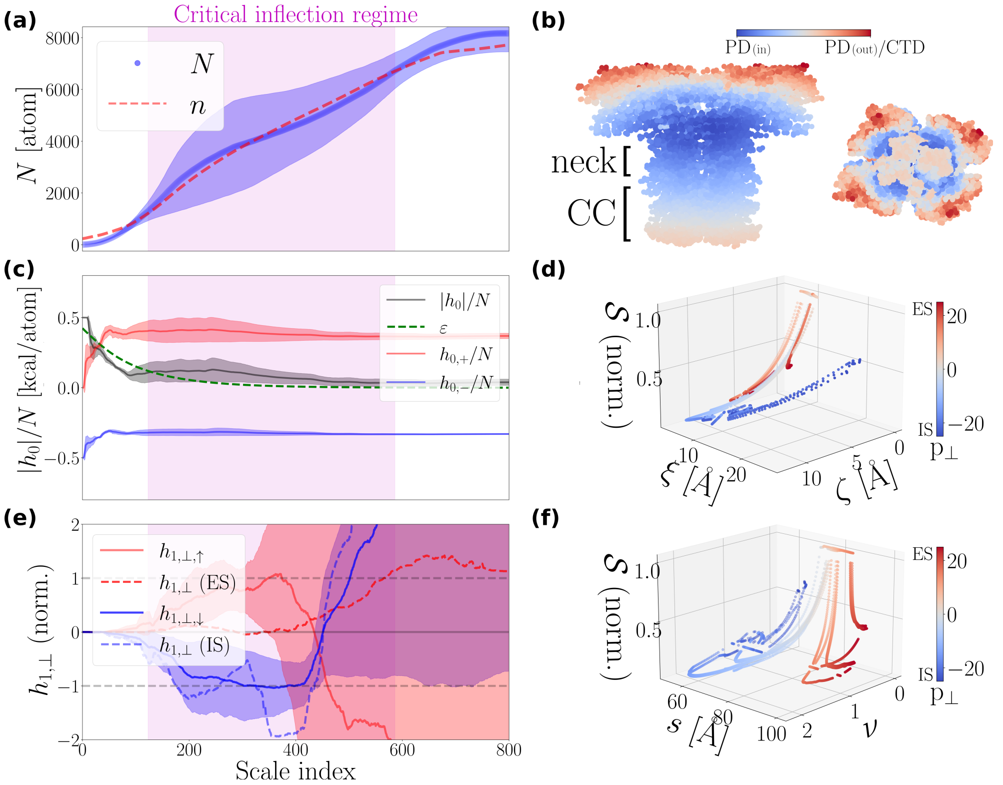


**Fig. S7**: **Spatial organization observables of 7 NaVAe PD/CTD molecular assemblies (see Tab. S1).** **(a)**, **(b)**, **(c)**, **(d)**, **(e)**, **(f)** illustrate same type observables as in Fig. (S5), computed using the same data analytics procedure (see S1.8.1), for $\mathbf{p} \in \mathcal{P}$ and $\alpha = 1, 2, \ldots, N_\alpha = 800$. In **(b)**, we illustrate the NaVAe structure with PDB code 5hk7. NaVAe assemblies comprise a pore domain (PD) extending towards a C-terminal domain (CTD). The CTD consists of a flexible hydrophilic neck connecting the PD with a four-helix coiled-coil (CC). Note that $\langle l_i \rangle_{\mathrm{p}}$ [Å] is not shown here as it does not convey any useful information. Instead, two prominent inflection points emerge: one governing the transition from the core (denoted as $\mathrm{PD}_{(\mathrm{in})}$) towards the CTD, and a second one governing the transition from the core towards the molecular periphery (denoted as $\mathrm{PD}_{(\mathrm{out})}$).

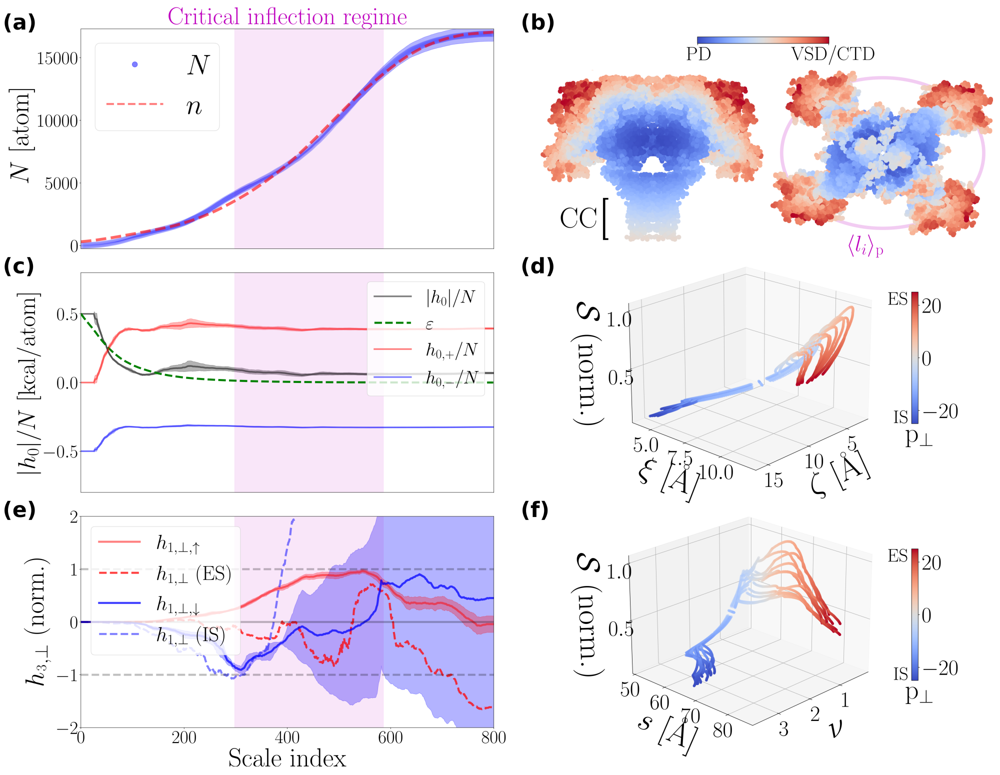


**Fig. S8**: **Spatial organization observables of 11 NaVMs PD/VSD/CTD molecular assemblies (see Tab. S1)**. **(a)**, **(b)**, **(c)**, **(d)**, **(e)**, **(f)** illustrate same type observables as in Fig. (S5), computed using the same data analytics procedure (see S1.8.1), for $\mathbf{p} \in \mathcal{P}$ and $\alpha = 1, 2, \ldots, N_\alpha = 800$. In **(b)**, we illustrate the NaVMs molecule with PDB code 5hvx. It consists of a pore domain (PD) coupled with four voltage-sensor domains (VSDs). The PD extends towards a C-terminal domain (CTD) with a four-helix coiled-coil (CC). Notable, incorporating the CTD does not diminish the prominence of the inflection point governing the transition from the PD to the VSDs.

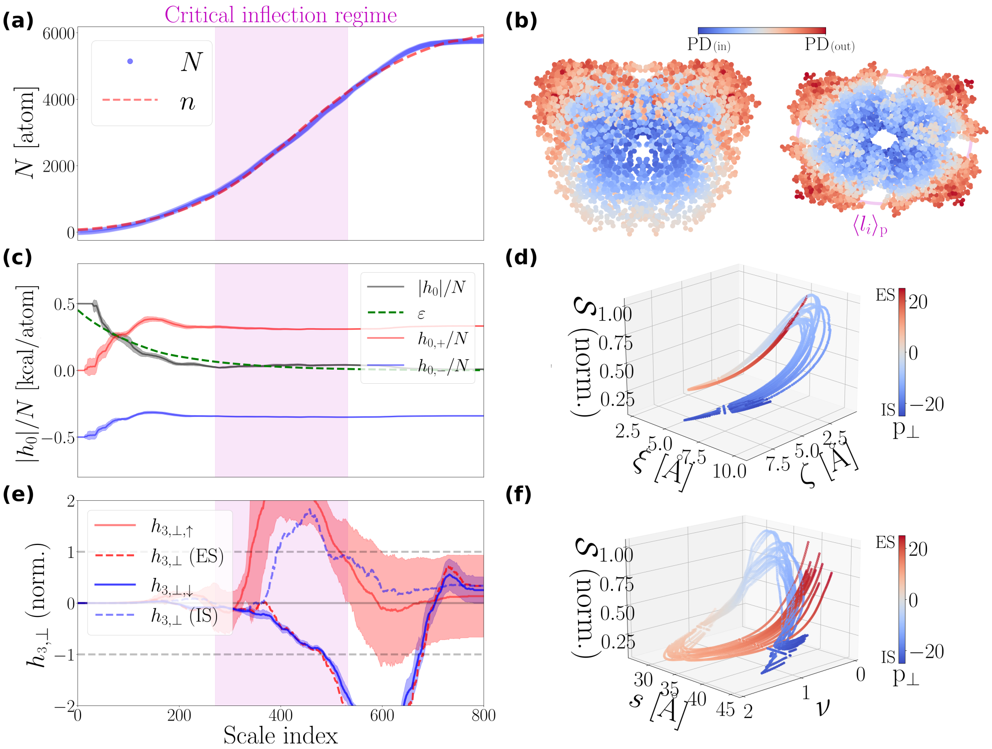


**Fig. S9**: **Spatial organization of 15 NaVMs pore-only molecules (see Tab. S1)**. **(a)**, **(b)**, **(c)**, **(d)**, **(e)**, **(f)** illustrate same type observables as in Fig. (S5), computed using the same data analytics procedure (see S1.8.1), for $\mathbf{p} \in \mathcal{P}$ and $\alpha = 1, 2, \ldots, N_\alpha = 800$. In **(b)**, we illustrate the NaVMs pore-only structure with PDB code 4cbc. Note that in **(e)**, we use the third-order hydropathic moment to capture critical transients, since the first-order moment suffers from strong fluctuations.

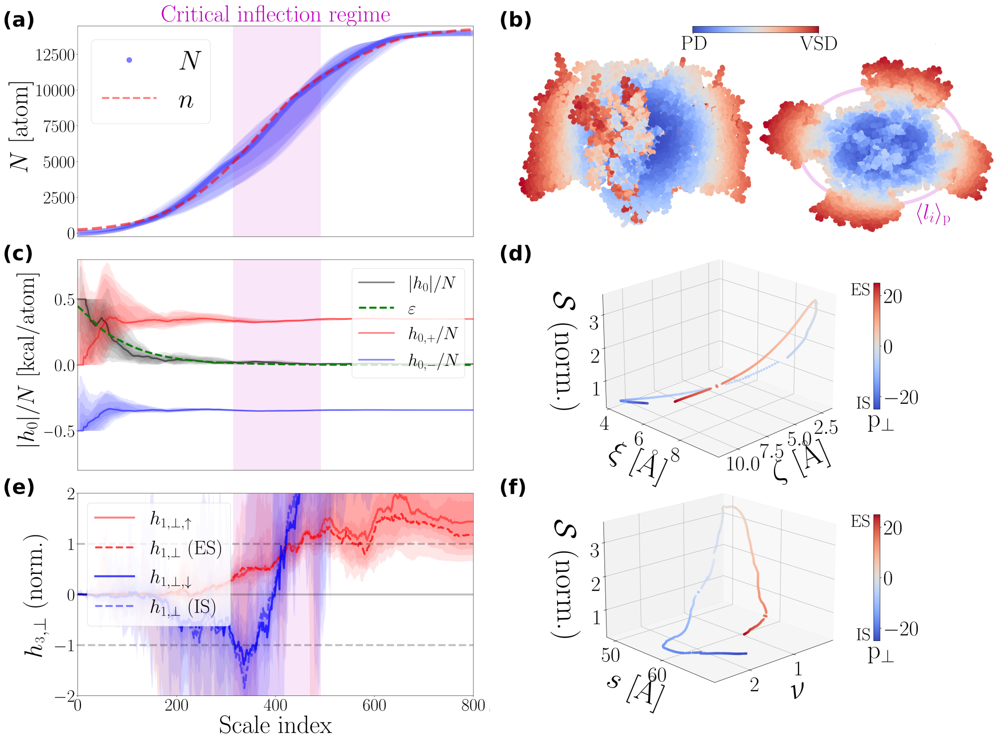


**Fig. S10**: **Spatial organization of a single NaVRh molecule (PDB code: 4dxw)**. **(a)**, **(b)**, **(c)**, **(d)**, **(e)**, **(f)** illustrate same type observables as in Fig. (S5), computed using the same data analytics procedure (see S1.8.1), for $\mathbf{p} \in \mathcal{P}$ and $\alpha = 1, 2, \ldots, N_\alpha = 800$.

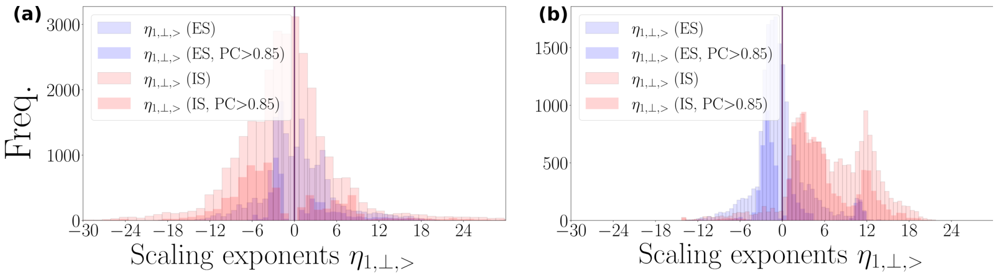


**Fig. S11**: **Distributional preferences of prokaryotic and eukaryotic $\eta_{1,\perp,>}$-exponents**. We split the parent $\eta_{1,\perp,>}$-distribution into two sub-distributions, denoted with ES and IS, accounting for $\eta_{1,\perp,>}$-exponents characterizing the scaling behavior of hydropathic dipole field amplitude, $|h_{1,\perp}|$ [kcal×Å], acting at the extracellular side (ES) of the (i.e., $p_\perp > 0$) and at the intracellular side (IS) of the pore (i.e., $p_\perp < 0$), respectively. We remind that $\eta_{1,\perp,>}$-exponents characterize the scaling behavior of $|h_{1,\perp}|$ in the VSDs. **(a)** and **(b)** show the results for 71 prokaryotic and 50 eukaryotic NaVCh structures (Tab. S1). PC stands for Pearson coefficient.

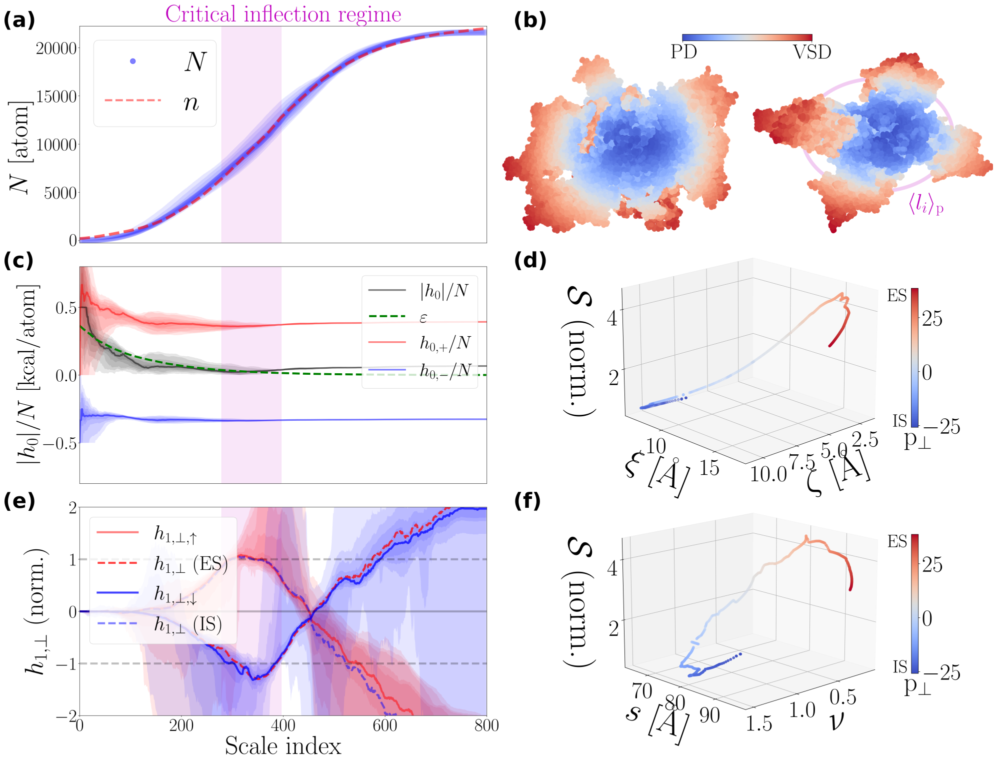


**Fig. S12**: **Spatial organization of a single NaVPas molecule (PDB code: 6a95)**. **(a)**, **(b)**, **(c)**, **(d)**, **(e)**, **(f)** illustrate same type observables as in Fig. (S5), computed using the same data analytics procedure (see S1.8.1), for $\mathbf{p} \in \mathcal{P}$ and $\alpha = 1, 2, \ldots, N_\alpha = 800$.

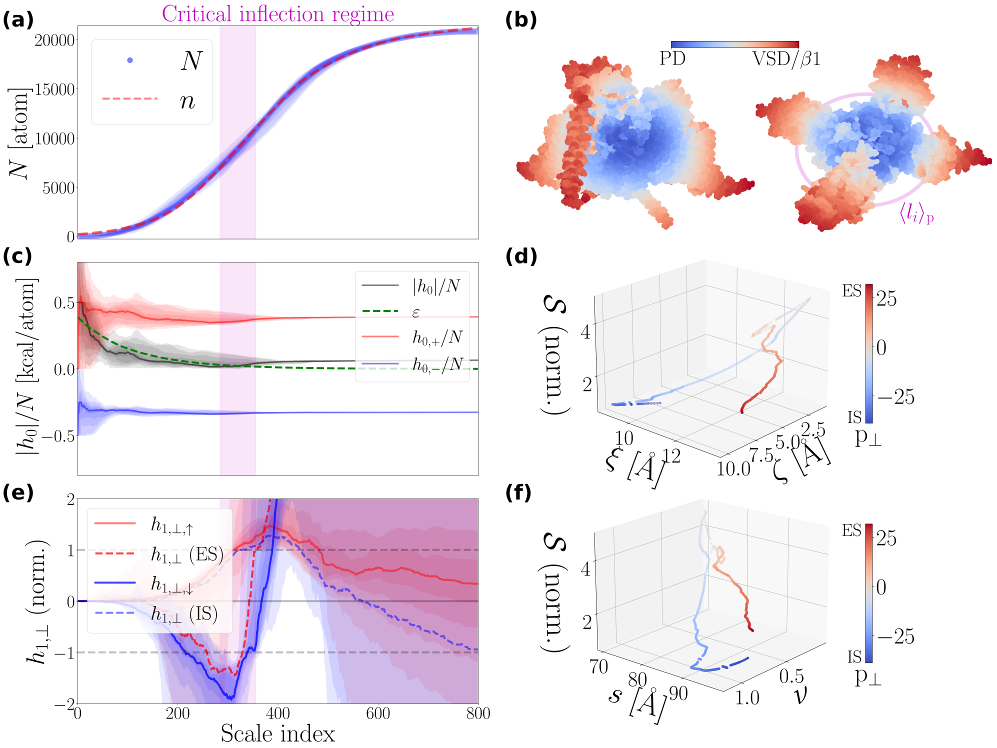


**Fig. S13**: **Spatial organization of a single NaVEe1 molecule (PDB code: 5xsy)**. **(a)**, **(b)**, **(c)**, **(d)**, **(e)**, **(f)** illustrate same type observables as in Fig. (S5), computed using the same data analytics procedure (see S1.8.1), for $\mathbf{p} \in \mathcal{P}$ and $\alpha = 1, 2, \ldots, N_\alpha = 800$. Note the structural inclusion of the $\beta 1$ subunit.

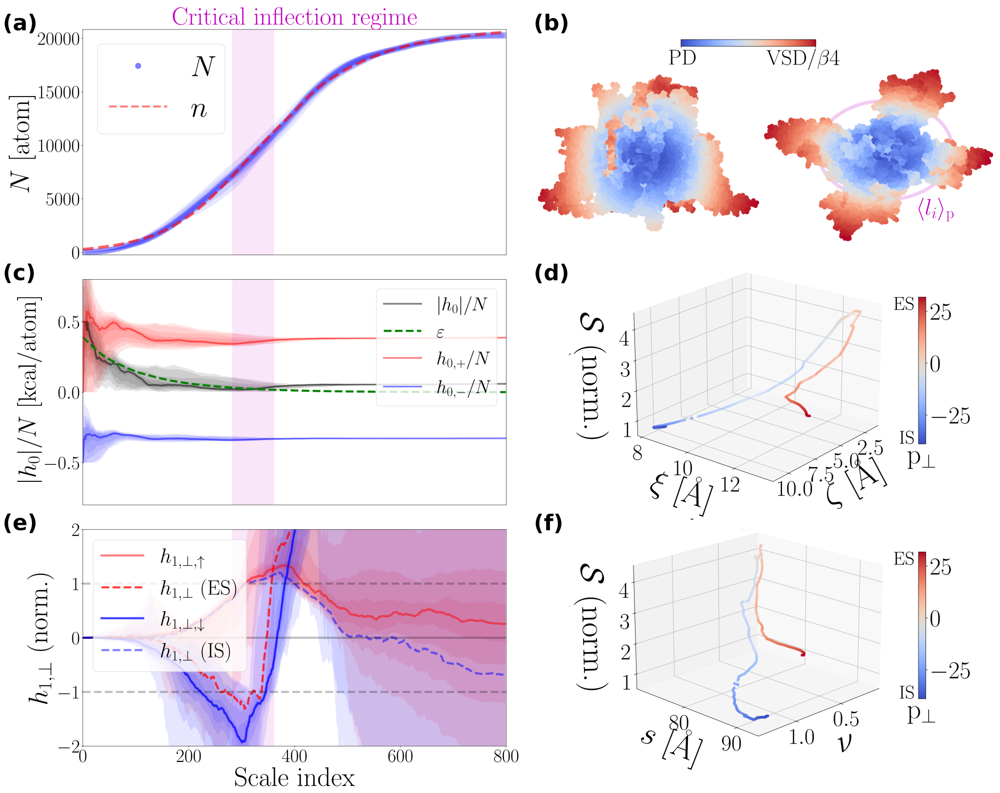


**Fig. S14**: **Spatial organization of a single NaV1.1 molecule (PDB code: 7dtd)**. **(a)**, **(b)**, **(c)**, **(d)**, **(e)**, **(f)** illustrate same type observables as in Fig. (S5), computed using the same data analytics procedure (see S1.8.1), for $\mathbf{p} \in \mathcal{P}$ and $\alpha = 1, 2, \ldots, N_\alpha = 800$. Note the structural inclusion of the $\beta 4$ subunit.

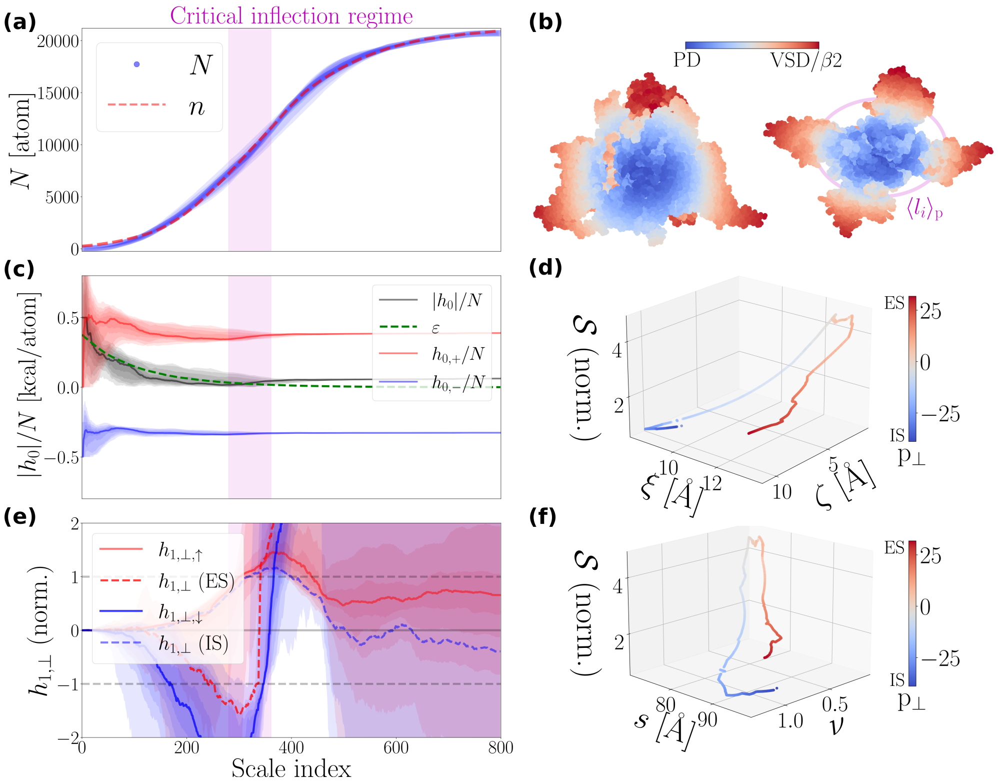


**Fig. S15**: **Spatial organization of a single NaV1.2 molecule (PDB code: 6j8e)**. **(a)**, **(b)**, **(c)**, **(d)**, **(e)**, **(f)** illustrate same type observables as in Fig. (S5), computed using the same data analytics procedure (see S1.8.1), for $\mathbf{p} \in \mathcal{P}$ and $\alpha = 1, 2, \ldots, N_\alpha = 800$. Note the structural inclusion of the $\beta 2$ subunit.

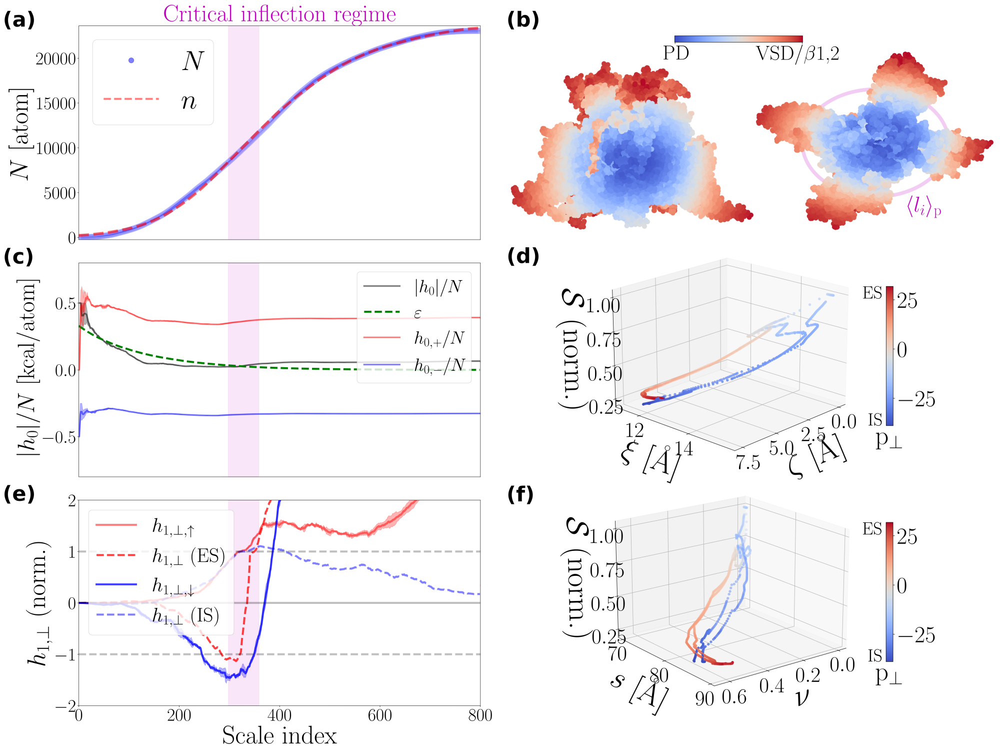


**Fig. S16**: **Spatial organization of 2 NaV1.3 molecules (PDB codes: 7w77, 7w7f)**. **(a)**, **(b)**, **(c)**, **(d)**, **(e)**, **(f)** illustrate same type observables as in Fig. (S5), computed using the same data analytics procedure (see S1.8.1), for $\mathbf{p} \in \mathcal{P}$ and $\alpha = 1, 2, \ldots, N_\alpha = 800$. In **(b)**, we illustrate the NaV1.3 molecule with PDB code 7w7f. Note the structural inclusion of the $\beta 1, 2$ subunits.

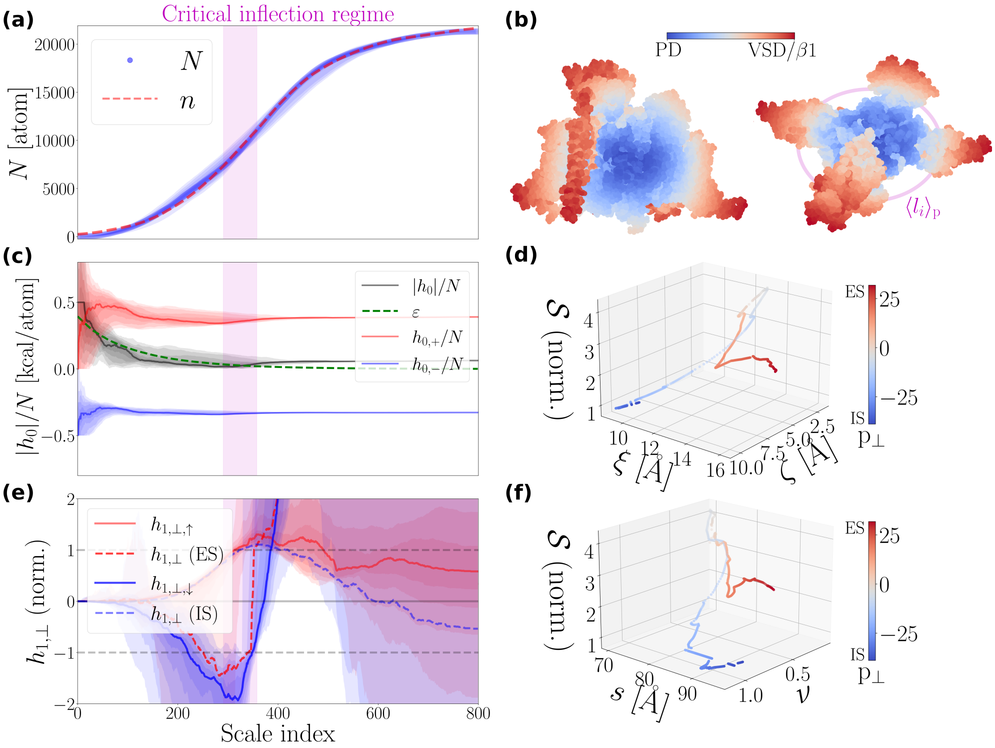


**Fig. S17**: **Spatial organization of a single NaV1.4 molecule (PDB code: 6agf)**. **(a)**, **(b)**, **(c)**, **(d)**, **(e)**, **(f)** illustrate same type observables as in Fig. (S5), computed using the same data analytics procedure (see S1.8.1), for $\mathbf{p} \in \mathcal{P}$ and $\alpha = 1, 2, \ldots, N_\alpha = 800$. Note the structural inclusion of the $\beta 1$ subunit.

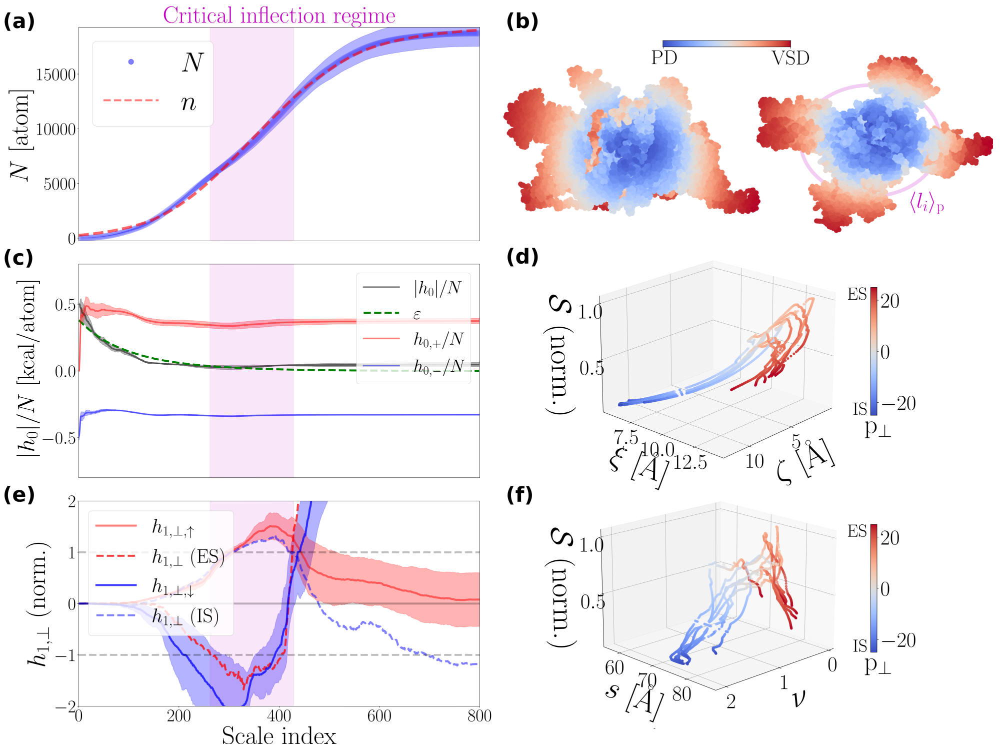


**Fig. S18**: **Spatial organization of 7 NaV1.5 molecules (see Tab. S1)**. **(a)**, **(b)**, **(c)**, **(d)**, **(e)**, **(f)** illustrate same type observables as in Fig. (S5), computed using the same data analytics procedure (see S1.8.1), for $\mathbf{p} \in \mathcal{P}$ and $\alpha = 1, 2, \ldots, N_\alpha = 800$. In **(b)** we illustrate the molecule with PDB code 7k18.

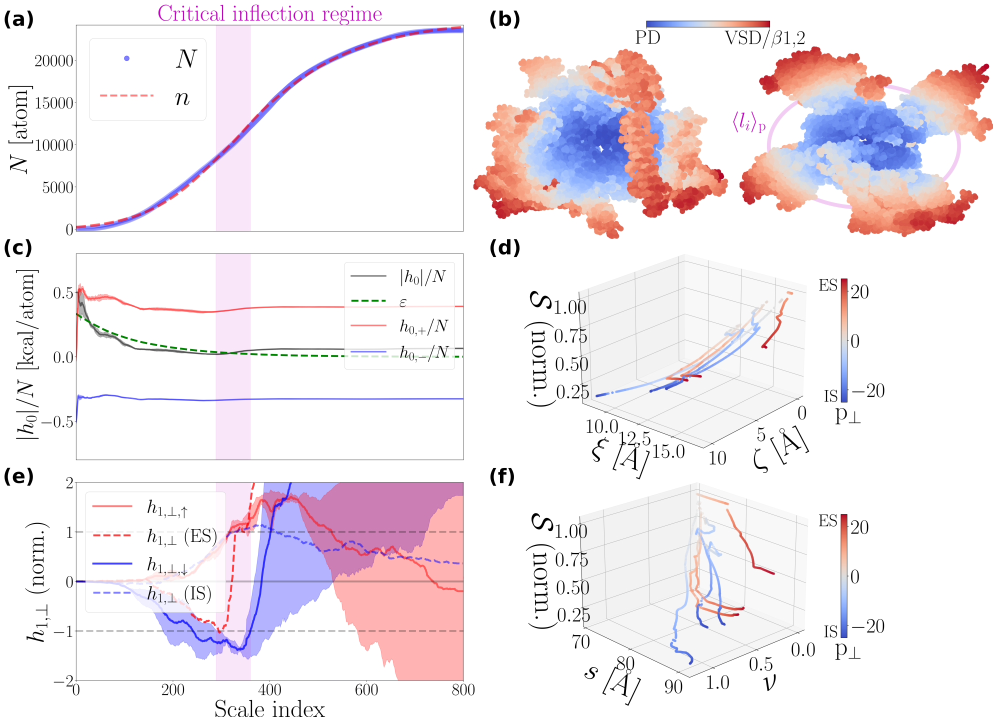

**Fig. S19**: **Spatial organization of 3 NaV1.6 molecules (see Tab. S1)**. **(a)**, **(b)**, **(c)**, **(d)**, **(e)**, **(f)** illustrate same type observables as in Fig. (S5), computed using the same data analytics procedure (see S1.8.1), for $\mathbf{p} \in \mathcal{P}$ and $\alpha = 1, 2, \ldots, N_\alpha = 800$. In **(b)** we illustrate the molecule with PDB code 8fhd. The 8fhd structure incorporates the $\beta1$ subunit, while the structures 8gz1 and 8gz2 incoroporate both $\beta1$ and $\beta2$ subunits.

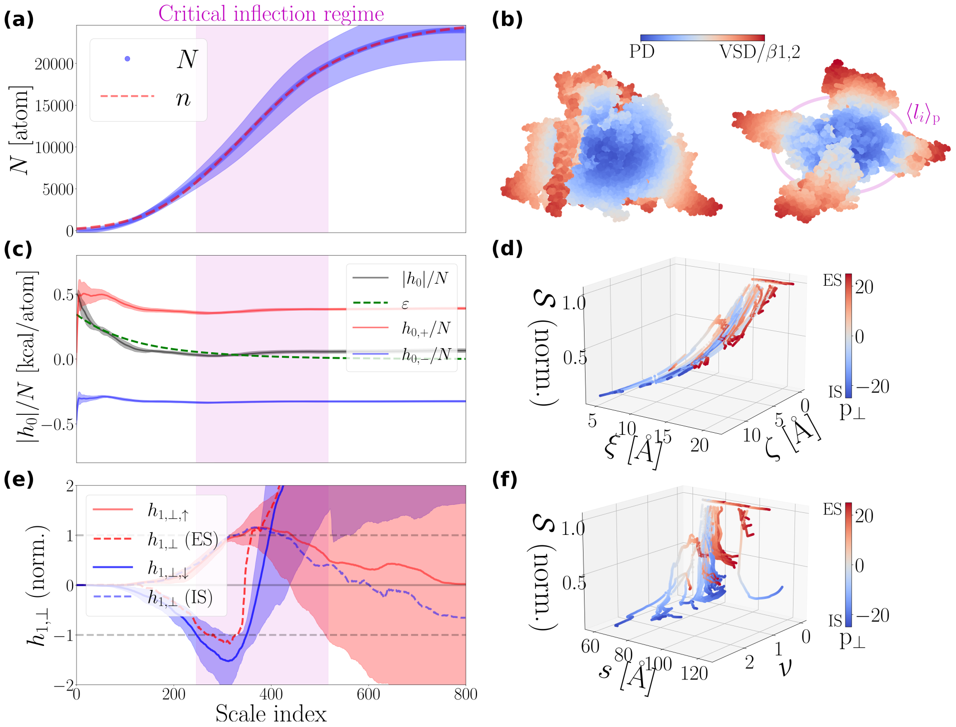


**Fig. S20**: **Spatial organization of 28 NaV1.7 molecules (see Tab. S1)**. **(a)**, **(b)**, **(c)**, **(d)**, **(e)**, **(f)** illustrate same type observables as in Fig. (S5), computed using the same data analytics procedure (see S1.8.1), for $\mathbf{p}\in\mathcal{P}$ and $\alpha=1,2,\dots,N_\alpha=800$. In **(b)** we illustrate the molecule with PDB code 6j8j. All structures incorporate the $\beta$1 and $\beta$2 subunits except for 5ek0, 6ntq, 6nt3, 6nt4, 8f0p, 8f0q, 8f0r, 8f0s, which are 'naked' of $\beta$ subunits. 8j4f 'wears' only the $\beta$1 subunit.

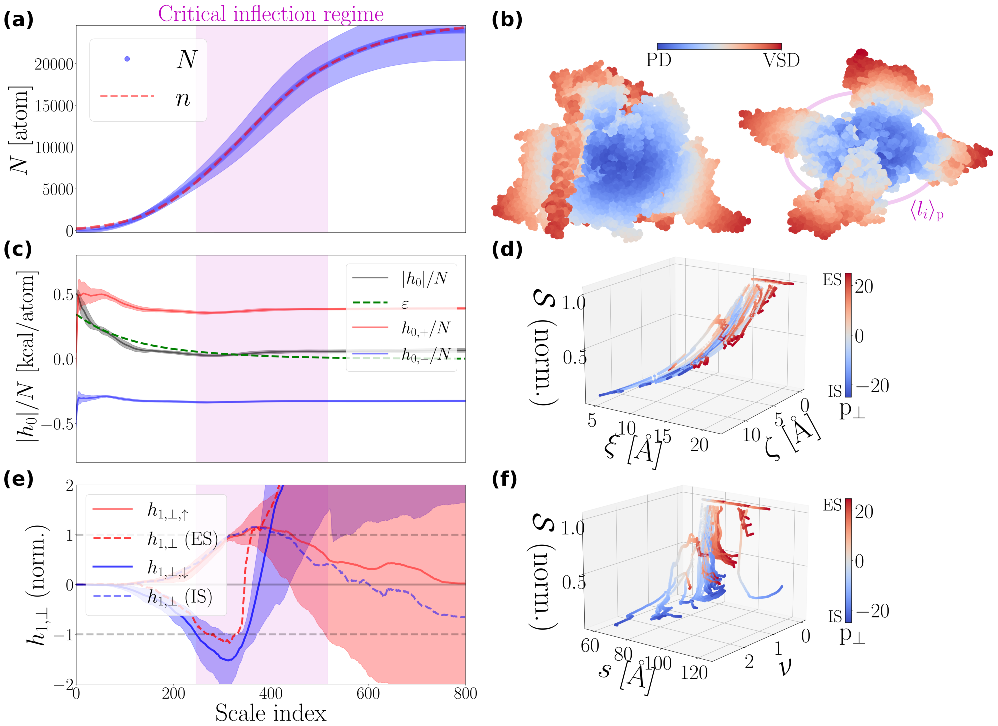


**Fig. S21**: **Spatial organization of 4 NaV1.8 molecules (see Tab. S1).** **(a)**, **(b)**, **(c)**, **(d)**, **(e)**, **(f)** illustrate same type observables as in Fig. (S5), computed using the same data analytics procedure (see S1.8.1), for $\mathbf{p} \in \mathcal{P}$ and $\alpha = 1, 2, \ldots, N_\alpha = 800$. In **(b)** we illustrate the molecule with PDB code 7we4.

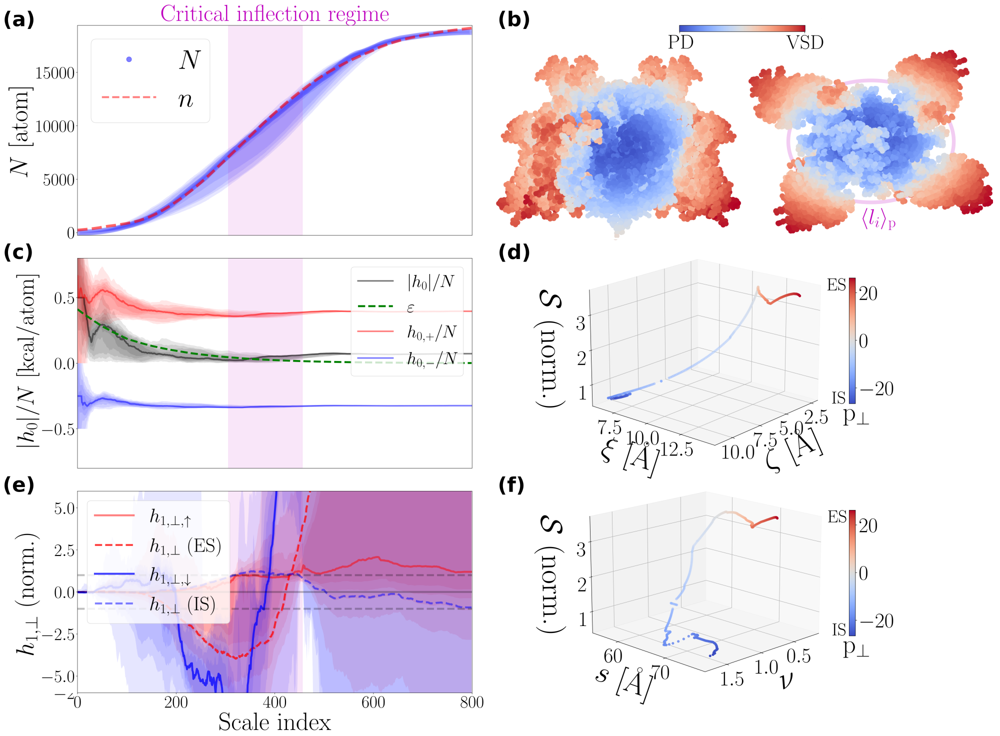


**Fig. S22**: **Spatial organization of a single NaVEh molecule (PDB code: 7x5v)**. **(a)**, **(b)**, **(c)**, **(d)**, **(e)**, **(f)** illustrate same type observables as in Fig. (S5), computed using the same data analytics procedure (see S1.8.1), for $\mathbf{p} \in \mathcal{P}$ and $\alpha = 1, 2, \ldots, N_\alpha = 800$.

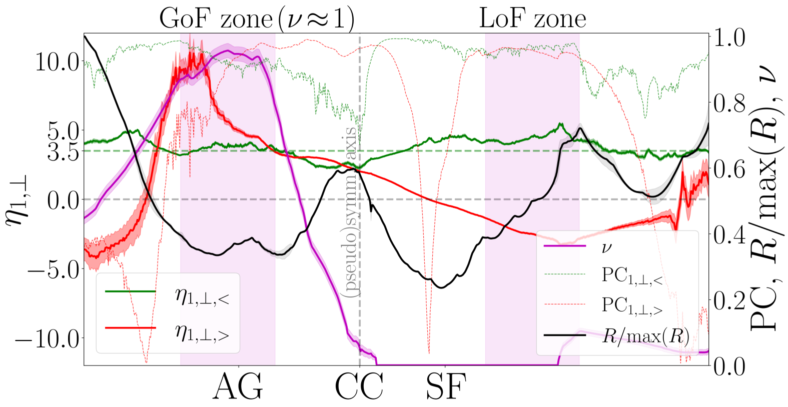


**Fig. S23**: **Exponent spectra of a NaV1.7 molecule (PDB code: 7w9k)**. We plot the pair of the hydropathic dipole field (HDF) exponents, $\{\eta_{1,\perp,<}, \eta_{1,\perp,>}\}$, with $\{PC_{1,\perp,<}, PC_{1,\perp,>}\}$ being their corresponding Pearson coefficients (PCs) for $\mathbf{p} \in \mathcal{P}$. $\eta_{1,\perp,<}$ and $\eta_{1,\perp,>}$ inform about the scaling behavior of HDF amplitude, $|h_{1,\perp}|$ [kcal×Å], in the PD (i.e., $l < l_i$) and beyond it (i.e., $l > l_i$). Moreover, we show the trace of the interaction range ratio, $\nu$, and of the normalized pore radius, $R/\max\{R\}$. In light magenta, we highlight the pore regions, i.e., subsets of $\mathcal{P}$, toward which gain-of-function (GoF) and loss-of-function (LoF) mutation clouds tend to converge, i.e., minimize their Euclidean distance (see also Main Text Figs. 4(g),(f)). To highlight exponent spectrum symmetries we introduce a pseudosymmetry axis placed approximately in the center of the pore. Light magenta-colored zones highlight the pore regions where the distance to the structural locations of GoF and LoF mutations is minimized in a statistical sense.

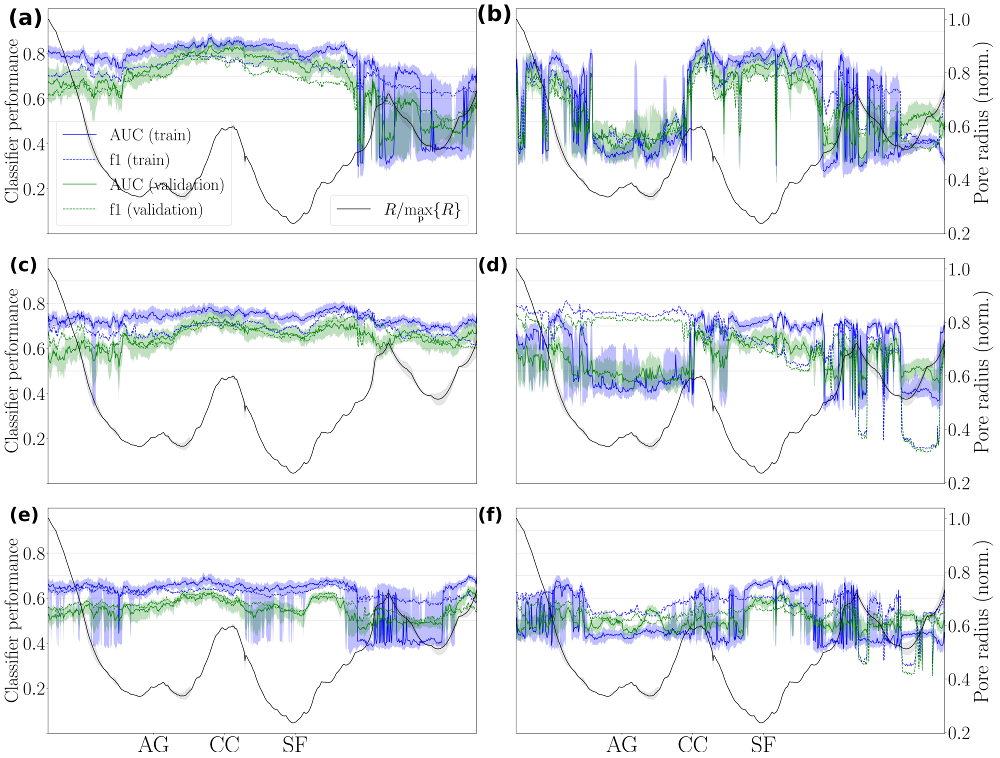


**Fig. S24**: **Local learning of structural location perturbation potential based on inertia constraints**. We plot the medians of the area under the curve (AUC) and the f1 scores obtained after running Alg. 5. Note that at each pore point, $\mathbf{p} \in \mathcal{P}$, the medians were obtained by compiling $(N_{\text{call}} = 20) \times (K_{\text{fold}} - 1 = 2) = 40$ score values (Tab. S4). The shaded area around a median trace illustrates the 25%-75% percentile interval. In **(a)**, **(c)**, and **(e)** we illustrate the median traces obtained by the cluster inertia inputs $\phi_{2k}$, $k = 0, 1, \ldots, 5\}$, for different machine learning experiments processing datasets I, II, and III. Dataset I, II, and III contains the classes $\{$`class_0`: GoF∪LoF, `class_1`: Neutr$\}$, $\{$`class_0`: GoF∪LoF, `class_1`: Neutr.∪Benign$\}$, and $\{$`class_0`: Path, `class_1`: Neutr.∪Benign$\}$, respectively. **(b)**, **(d)**, and **(f)**: equivalent to **(a)**, **(c)**, and **(e)**, respectively, but for the interfacial inertia inputs $\mathcal{I}_{2k}$, $k = 0, 1, \ldots, 5$. AG, CC, and SF stand for activation gate, central cavity, and selectivity filter, respectively. GoF and LoF stand for gain-of-function and loss-of-function, respectively, and represent mutations associated with increased or diminished pain sensation (listed in SI Tab. S3). Neutrals (Neutr.) are carefully selected human pain disease negative controls, sourced from [10, 19]. Benign are generally not expected to be associated with a disease phenotype. Note that the pathogenic (path.) class contains both pain-disease-associated and pain-disease-unrelated pathogenic mutations (Main Text Paragraph 3.2).

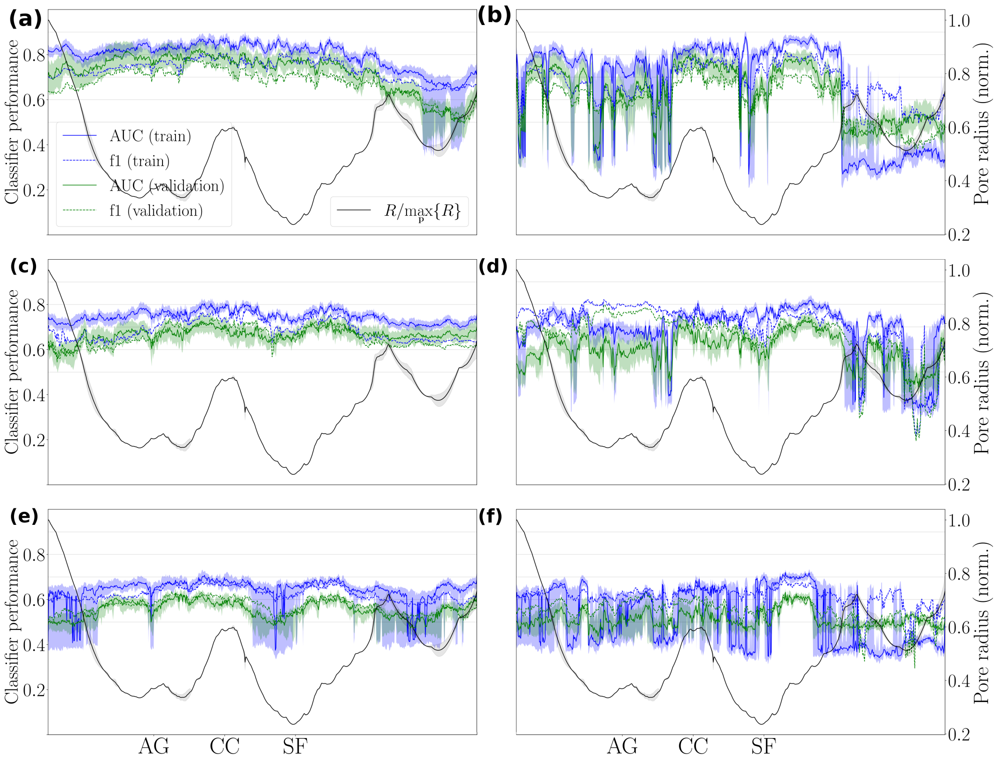


**Fig. S25**: **Local learning of structural location perturbation potential based on conductivity constraints**. We maintain the same format and compile the same information concerning mutation datasets as in Fig. S24, but instead, in **(a)**, **(c)**, and **(e)** we illustrate the median traces obtained from the cluster conductivity feature inputs $\phi_{2k+1,\perp}$, $k=0,1,\ldots,5$, while in **(b)**, **(d)**, and **(f)** we illustrate the interfacial conductivity feature inputs $\mathcal{I}_{2k+1,\perp}$, $k=0,1,\ldots,5$.

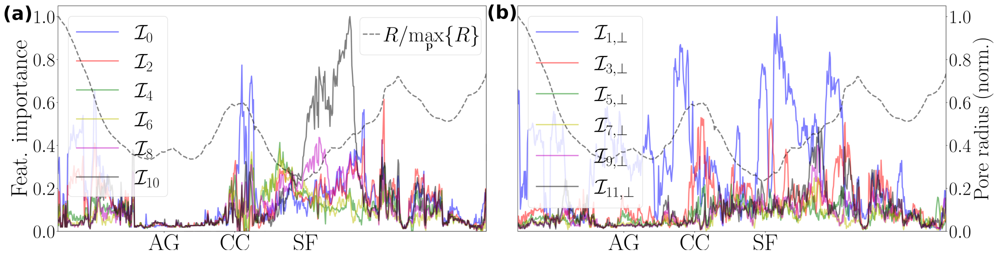


**Fig. S26**: **Interfacial feature input importance**. The local (i.e., **p**-dependent) importance of a feature input is evaluated in terms of the median of the score values returned by the `permutation_importance` function found in the `sklearn` library. Executing Alg. 5 generates to total of $(N_{\text{perm.}}=10)\times(N_{\text{call}}=20\times(K_{\text{fold}}-1=2)=400$ score values (see Tab. S4 for parameter value listing), which are then aggregated to derive local median statistics. In **(a)** and **(b)** we illustrate the median traces for the interfacial inertia, $\mathcal{I}_{2k}$, and interfacial conductivity, $\mathcal{I}_{2k+1,\perp}$, respectively, feature inputs. We consider the dataset of pain-disease-associated (GoF$\cup$LoF) and neutral mutations. GoF and LoF stand for gain-of-function and loss-of-function, respectively, and represent mutations associated with increased or diminished pain sensation (listed in SI Tab. S3). Neutrals (Neutr.) are carefully selected human pain disease negative controls, sourced from [10, 19]. Note that the medians are normalized over the maximum value in their dataset. Note that Figs. (6)(c) and 6(d) appearing in the Main Text illustrate the equivalent of **(a)** and **(b)** but for $\phi_{2k}$ and $\phi_{2k+1,\perp}$, respectively.

**Table S2**: **Atomic parameters.** The CVECT parameter specifies the direction of the pore with respect to $(\hat{\mathbf{e}}_x, \hat{\mathbf{e}}_y, \hat{\mathbf{e}}_z)$. The SAMPLE parameter, $\Delta z$ [Å], sets the distance between consecutive $xy$-planes. The parameters MCKT and MCDISP define the initial Boltzmann factor and maximum step length for the Metropolis Monte Carlo simulated annealing optimization routine of HOLE, respectively. The ENDRAD parameter specifies the maximum pore radius value at which the pore ends. The chosen value for the ENDRAD parameter leads to an average of approximately 690 pore points per structure, implying that the average length of a NaVCh pore is approximately $690 \times \Delta z = 69$ Å. The RASEED parameter sets the seed for the $k$-th HOLE routine call. CPOINT refers to the initial point, defined as the molecular center, **c**, of the 'clean', protonated, and oriented structure. Van der Waals atomic radii are imported from Ref. [20].

| HOLE routine parameter | Value | Atom | Van der Waals radius [Å] |
|---|---|---|---|
| CVECT | 0.0 0.0 1.0 | C | 1.65 |
| MCDISP | 0.05 | O | 1.4 |
| MCKT | 0.2 | S | 1.8 |
| SAMPLE | 0.1 Å | N | 1.55 |
| ENDRAD | 8.0 Å | H | 1.0 |
| RASEED | $k \times 1000$ | P | 1.8 |
| CPOINT | $c_x$ $c_y$ $c_z$ | | |

**Table S3**: **Pain-disease-associated mutations.** We consider a set of gain-of-function (GoF) and loss-of-function (LoF) mutations. GoF and LoF mutations are typically interlinked with increased and decreased, respectively, ionic net currents. GoF mutations are associated with one of the three following pain phenotypes: inherited erythromelalgia (IEM), paroxysmal extreme pain disorder (PEPD), and small fiber neuropathy (SFN). LoF mutations typically cause pain insensitivity (pain insens.). If the disease phenotype of a LoF mutation is unknown, but only the electrophysiological signature is known, the mutation is indicated with an asterisk (*).

| GoF | | | LoF |
|---|---|---|---|
| IEM | PEPD | SFN | Pain insens. or unknown |
| I136V [21, 22] | V1298F [23, 24] | R185H [25] | A99H [26] |
| S211P [27] | V1299F [24, 28] | I228M [29] | L172R [30] |
| F216S [31] | I1461T [32] | I720K [33] | F344I* [34] |
| I234T [35] | G1607R [36] | I739V [25] | R896Q [37] |
| S241T [38, 39] | L1612P [40] | M932L [33] | M899I [41] |
| L245V [42] | M1627K [43] | R1279P [44] | W917G [26] |
| N395K [45] | A1632E [46] | T1596I [47] | F1378I* [34] |
| V400M [48] | | | F1670I* [34] |
| L823R [49] | | | A1687P [30] |
| F826Y [50] | | | C1719R [51] |
| I848T [52, 53] | | | |
| G856R [54] | | | |
| G856D [55] | | | |
| L858H [56, 57] | | | |
| L858F [58] | | | |
| A863P [59] | | | |
| V872G [60] | | | |
| Q875E [61] | | | |
| L955Del [62] | | | |
| P1308L [24] | | | |
| V1316A [63, 64] | | | |
| F1449V [65] | | | |
| W1538R [66] | | | |
| A1632T [67] | | | |
| A1632G [68] | | | |
| A1746G [66] | | | |

**Table S4**: **Machine learning experiments parametrization**. $K_{\text{fold}}$ is the number of groups (folds) considered during the k-fold cross-validation algorithm illustrated in Alg. 4. $N_{\text{call}}$ is the number of Alg. 4 is called (see Algs. 5,6). $\text{seed}_{\text{classif}}$ is the seed used for calling the SVC. $\text{seed}_{\text{shuffle}}$ is the seed used for shuffling (see Alg. 4). $N_{\text{perm}}$ is the number of permutations performed during feature importance estimation. Feature importance estimated with the `permutation_importance` function found in the `sklearn` library.

| | Local learning | Global Learning |
|---|---|---|
| Parameter | Value | Value |
| $K_{\text{fold}}$ | 3 | 3 |
| $N_{\text{call}}$ | 20 | 50 |
| $\text{seed}_{\text{classif}}$ | 42 | 42 |
| $\text{seed}_{\text{shuffle}}$ | $i-1$ | $i-1$ |
| $N_{\text{perm}}$ | 10 | 10 |

**Table S5**: **Machine-learning experiments summary with abolished directionality.** The first and second number of the pair $(\cdot,\cdot)$ represent median values, rounded to two decimal digits, obtained from model training and testing (validating) during the final classification round, respectively. Undirectional inertia (iner.) and conductivity (cond.) constraints are probed by the unsigned feature inputs $\{|\phi_{2k}|, |\mathcal{I}_{2k}|\}$ and $\{|\phi_{2k+1,\perp}|, |\mathcal{I}_{2k+1,\perp}|\}$, respectively. *SCN9A*-gene mutation datasets I, II, and III contain the classes {`class_0`: GoF∪LoF, `class_1`: Neutr.}, {`class_0`: GoF∪LoF, `class_1`: Neutr.∪Benign}, and {`class_0`: Path., `class_1`: Neutr.∪Benign}, respectively. GoF and LoF stand for gain-of-function and loss-of-function, respectively, and represent mutations associated with increased or diminished pain sensation (listed in SI Tab. S3). Neutrals (Neutr.) are carefully selected human pain disease negative controls, sourced from [10, 19]. Benign are generally not expected to be associated with a disease phenotype. Note that the pathogenic (path.) class contains both pain-disease-associated and pain-disease-unrelated pathogenic mutations (Main Text Paragraph 3.2).

| | PDB: 7w9k, res.: 2.2 Å | | | PDB: 6j8j, res.: 3.2 Å | | |
|---|---|---|---|---|---|---|
| Pert. modes | iner./cond. | iner. | cond. | iner./cond. | iner. | cond. |
| AUC (dataset I) | (0.90, 0.84) | (0.86, 0.80) | (0.87, 0.85) | (0.91, 0.84) | (0.87, 0.82) | (0.88, 0.84) |
| f1 (dataset I) | (0.83, 0.79) | (0.81, 0.78) | (0.80, 0.79) | (0.80, 0.76) | (0.78, 0.74) | (0.80, 0.76) |
| AUC (dataset II) | (0.86, 0.76) | (0.82, 0.75) | (0.83, 0.77) | (0.87, 0.77) | (0.83, 0.74) | (0.79, 0.73) |
| f1 (dataset II) | (0.78, 0.74) | (0.77, 0.74) | (0.72, 0.70) | (0.81, 0.75) | (0.79, 0.75) | (0.74, 0.71) |
| AUC (dataset III) | (0.78, 0.66) | (0.73, 0.64) | (0.73, 0.65) | (0.78, 0.63) | (0.73, 0.62) | (0.71, 0.62) |
| f1 (dataset III) | (0.72, 0.64) | (0.69, 0.64) | (0.67, 0.63) | (0.71, 0.61) | (0.68, 0.59) | (0.69, 0.63) |
